# Genome evolution and glycosyltransferase diversification shape asperosaponin VI biosynthesis in *Dipsacus asperoides*

**DOI:** 10.64898/2026.11.03.763632

**Authors:** Run Yang, Changhao Fu, Xiu Zhao, Yan Zhao, Weiping Huang, Mingyu Li, Guisheng Xiang, Jinglei Xiong, Yina Wang, Geng Chen, Qize Liu, Shengchao Yang, Guanghui Zhang, Simei He, Yanli Liang

**Author notes:** These authors have contributed equally to this work (Run Yang, Changhao Fu, Xiu Zhao, Yan Zhao). Correspondence: Yanli Liang, Simei He, Guanghui Zhang, Shengchao Yang.

## Abstract

Deciphering the genomic and enzymatic features associated with high-abundance specialized metabolism is important for understanding and engineering bioactive natural product biosynthesis. Asperosaponin VI, a hederagenin-based saponin, accumulates at exceptionally high levels in Dipsacus asperoides roots and is by far the predominant triterpenoid saponin. Here, by integrating a chromosome-level genome assembly with comparative genomics, transcriptomics, metabolite profiling, and biochemical characterization, we reconstructed a major biosynthetic route to asperosaponin VI and identified diversification of triterpenoid-associated gene families. Functional analyses revealed an ordered glycosylation sequence in which DaUGT73GZ2 preferentially accepts C-3-glycosylated substrates retaining a free C-28 carboxyl group, thereby establishing a C-3-before-C-28 glycosylation order during asperosaponin VI biosynthesis. Phylogenetic and biochemical analyses further uncovered functional differentiation among UGT85CA enzymes involved in C-3 arabinosylation, while genomic and expression analyses revealed lineage-specific patterns of pathway gene duplication and transcriptional regulation. Together, these findings identify genomic, enzymatic, and transcriptional features associated with the distinctive, high-abundance triterpenoid saponin profile of D. asperoides roots and provide a framework for understanding and engineering triterpenoid saponin biosynthesis.

**Short Summary:** Integrated genomic and biochemical analyses of Dipsacus asperoides link the predominant accumulation of asperosaponin VI to gene duplication, glycosyltransferase diversification, ordered glycosylation, and root-biased expression. These findings connect genome evolution with specialized metabolic output and identify genes and enzymes useful for engineering triterpenoid saponin biosynthesis.

## INTRODUCTION

Plants synthesize an extraordinary diversity of specialized metabolites, many of which accumulate in highly lineage-, tissue- or organ-specific patterns (Pichersky and Lewinsohn, 2011; Huang and Dudareva, 2023). In some species, one compound or a narrow set of structurally related metabolites can reach exceptionally high abundance, generating dominant metabolic profiles with strong pathway directionality and product specificity (Johnson et al., 2017; Hordyjewska et al., 2019). Such lineage-specific metabolic outputs often arise through multilayered genomic and biochemical innovations, including gene duplication, copy-number variation, metabolic gene clustering, enzyme neofunctionalization, altered substrate specificity, and coordinated transcriptional regulation (Cheng et al., 2025). Classical examples illustrate different aspects of this process: glucosinolate diversification in Brassicaceae has been shaped by gene duplication and changes in enzyme substrate specificity, whereas avenacin biosynthesis in oat represents a well-characterized triterpenoid pathway encoded by a biosynthetic gene cluster (Qi et al., 2004; Li et al., 2021). Similar principles have also been observed in opium poppy and withanolide-producing Solanaceae species, where pathway gene clustering, copy-number variation and lineage-specific enzyme recruitment contribute to the evolution of alkaloid and withanolide biosynthesis (Li et al., 2020; Hakim et al., 2025). Nevertheless, how genome evolution, enzyme functional diversification, and transcriptional deployment are integrated to shape lineage-specific specialized metabolic outputs remains incompletely understood.

*Dipsacus asperoides* is a perennial herb in the Caprifoliaceae family and the source plant of the traditional Chinese medicinal material Xuduan (Tao et al., 2020) (Figure.1A). Its roots accumulate triterpenoid saponins as major specialized metabolites, with asperosaponin VI, a hederagenin-based saponin, representing the predominant constituent (Xu et al., 2022; Pan et al., 2023). Asperosaponin VI has been reported to reach approximately 15% of root dry weight and represents a dominant triterpenoid saponin in *D. asperoides* roots (Yang et al., 2025). Asperosaponin VI biosynthesis follows the general logic of triterpenoid saponin formation, beginning with 2,3-oxidosqualene cyclization and followed by oxidative tailoring and glycosylation reactions mediated by oxidosqualene cyclases, cytochrome P450 monooxygenases and UDP-dependent glycosyltransferases. Our previous biochemical studies have identified several key enzymes involved in asperosaponin VI biosynthesis in *D. asperoides*, including the *β*-amyrin synthases DaOSC2 and DaOSC5, the *C*-23 oxidases DaCYP714E29 and DaCYP714E30, and the *C*-3 arabinosyltransferase DaUGT85J2, thereby establishing a partial enzymatic framework for hederagenin-based saponin formation in *D. asperoides* (Yao et al., 2024; Niu et al., 2025; Yang et al., 2025). However, the genomic organization and evolutionary diversification of this pathway remain unresolved, particularly how pathway gene diversification, enzyme specialization, and tissue-specific expression are associated with asperosaponin VI biosynthesis and its predominant accumulation in roots. Key enzymatic and evolutionary questions also remain unanswered, including how sequential glycosylation is controlled and how UGT activities diversify during hederagenin-based saponin biosynthesis.

Here, we report the first chromosome-level genome assembly of *D. asperoides* and integrate comparative genomics, transcriptomics, metabolite profiling, and biochemical characterization to explore the molecular and evolutionary features associated with the preferential accumulation of asperosaponin VI. We reconstructed a major biosynthetic route to asperosaponin VI and identified an ordered glycosylation sequence governed by substrate-selective UGT activities. Comparative genomic and functional analyses revealed diversification of triterpenoid-associated gene families and functional differentiation among UGT85CA enzymes involved in *C*-3 arabinosylation. We further uncovered lineage-specific patterns of pathway gene duplication and transcriptional differentiation that correlate with the distinctive triterpenoid saponin profile of *D. asperoides* roots. Together, these findings establish a genomic and biochemical framework for understanding how gene duplication, enzyme specialization, and transcriptional regulation may contribute to high-abundance, lineage-specific triterpenoid saponin metabolism, while also providing valuable genetic and enzymatic resources for the engineering of triterpenoid saponin biosynthesis.

## RESULTS

### Asperosaponin VI dominates the root triterpenoid profile of *D. asperoides*

To determine whether triterpenoid metabolism is lineage- and/or organ-specifically enriched in *D. asperoides*, we performed LC–MS-based profiling of 18 pentacyclic triterpenoids in *D. asperoides* and two closely related Caprifoliaceae species, *Triplostegia glandulifera* and *Lonicera japonica*. Hederagenin-based saponins were markedly enriched in *D. asperoide*s, whereas only trace levels were detected in *L. japonica* and *T. glandulifera* (Supplementary Figures. 1-4 and Supplementary Table 1). By contrast, *T. glandulifera* displayed three prominent characteristic signals matching the calculated [M-H]^-^ions of triploside A (m/z 1407.68), triploside B (m/z 1275.63) and triploside C (m/z 1011.55), respectively (Ma et al., 1991) (Supplementary Figures. 5 and 6), suggesting the accumulation of species-specific oleanolic acid-type triterpenoid saponins.

We then quantified seven hederagenin-based saponins, including cauloside A (**6**), HN saponin F (**7**), asperosaponin VI (**8**), *α*-hederin (**9**), dipsacoside B (**10**), macranthoidin A (**11**) and macranthoidin B (**12**). Asperosaponin VI was the predominant saponin in *D. asperoides* roots (Figures. 1A-D), whereas downstream rhamnosylated derivatives, including dipsacoside B and macranthoidin A, were mainly enriched in flowers. We further quantified asperosaponin VI in roots from 209 *D. asperoides* accessions collected from different geographic regions, revealing an average content of 96.35 ± 31.40 mg g^−1^ DW (Figure. 1E and Supplementary Table 2). Together, these results indicate that *D. asperoides* roots represent a naturally high-flux triterpenoid system in which hederagenin-based saponin metabolism is strongly concentrated around a single dominant product, asperosaponin VI.

**Figure 1.**
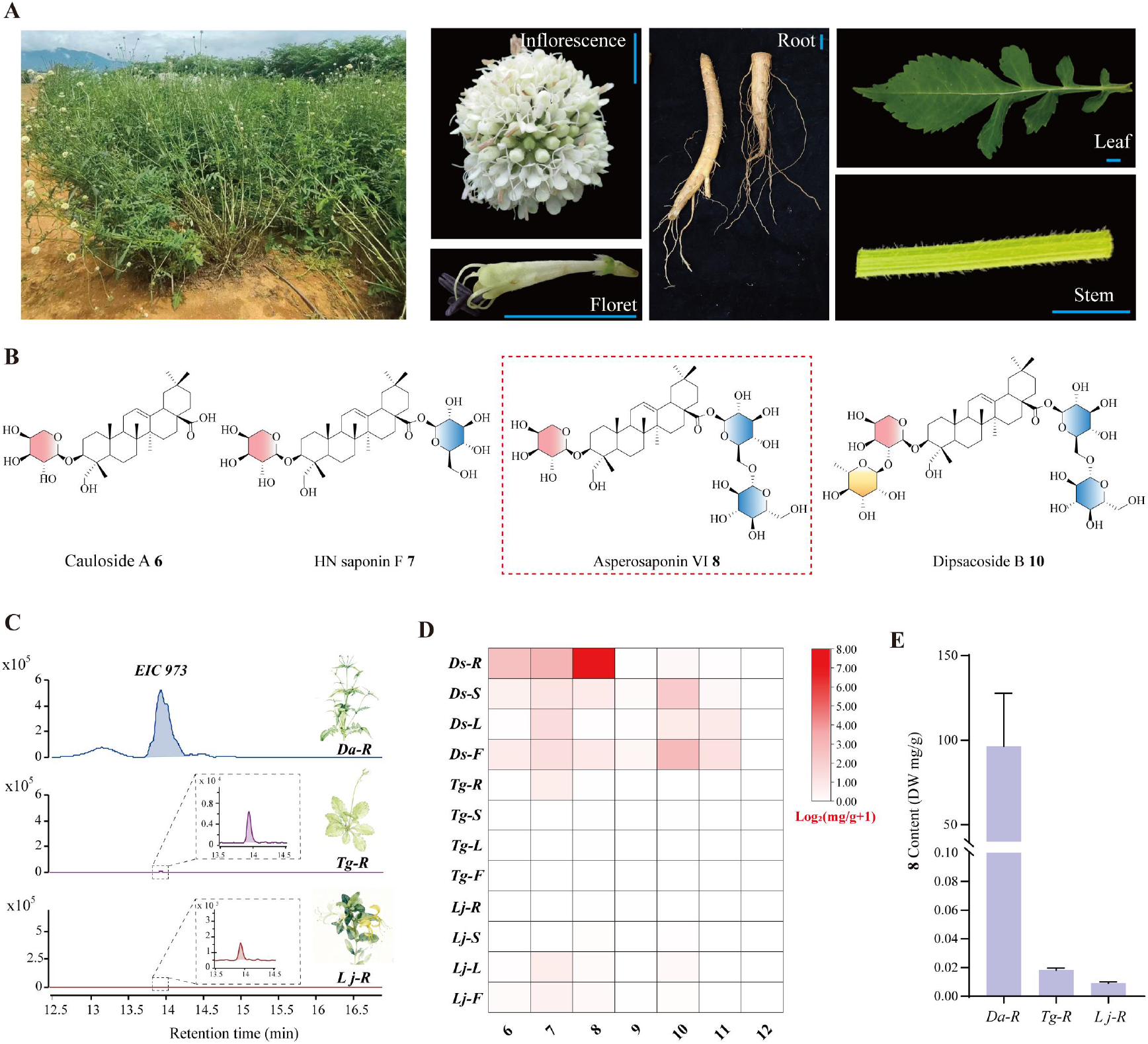
Morphological and chemotypic profiles of *D. asperoides*. **(A)** Morphological and tissue characteristics of *D. asperoides. Blue scale bars indicate 1 cm*. **(B)** Chemical structures of several major hederagenin-based saponins identified. (**C**) Extracted ion chromatograms (*EIC* m/z 973, corresponding to asperosaponin VI) of root extracts from the three representative Caprifoliaceae species (*D. asperoides, L. japonica, and T. glandulifera*). (**D**) Heatmap illustrating the distribution and absolute contents of hederagenin-based saponins across different tissues of *D. asperoides* (*Ds*) and two related species from the same family, *T. glandulifera* (*Tg*) and *L. japonica* (*Lj*). Abbreviations: *R*, root; *S*, stem; *L*, leaf; *F*, flower. (**E**) Absolute content of Asperosaponin VI in the roots of the three plant species.

### LTR expansion shapes the enlarged *D. asperoides* genome

To establish a genomic framework for investigating asperosaponin VI accumulation, we selected the same high-producing *D. asperoides* individual used for transcriptome profiling in our previous study (Yang et al., 2025) and generated the first chromosome-level reference genome for this species using PacBio HiFi and Hi-C sequencing. Genome survey analysis estimated a diploid genome size of 5.29 Gb with a heterozygosity rate of 0.57% (Supplementary Figures. 7 and 8). The final polished assembly was 5.23 Gb, with a contig N50 of 32.45 Mb and a scaffold N50 of 545.80 Mb, and 97.60% of the assembly was anchored onto nine pseudochromosomes (Supplementary Table 3 and Supplementary Figure. 9). Merqury analysis estimated a genome-wide quality value (QV) of 57.36, corresponding to an estimated consensus error rate of 1.84 × 10⁻⁶, indicating high base-level accuracy of the assembly. Telomeric repeats were detected at both ends of four pseudoch romosomes and at one end of each remaining pseudochromosome (Supplementary Table 4). BUSCO analysis recovered 99.1% of complete orthologs, and the LAI value of 21.62 indicated gold-standard assembly quality, supporting the high continuity and completeness of the *D. asperoides* genome assembly (Table 1 and Supplementary Table 5).

**Table 1.** The statistics of assembly and annotation of *D. asperoides*.

| Assembly and annotation | *D. asperoides* |
|---|---|
| Total number of contigs | 1,796 |
| Assembly size (bp) | 5,227,414,451 |
| N50 contig length (bp) | 32,451,949 |
| Total scaffold number | 414 |
| N50 scaffold length (bp) | 545,804,658 |
| Total number of pseudo-chromosomes | 9 |
| Sequence on chromosomes (bp) | 5,101,940,005 |
| Complete BUSCO for genome assembly | 99.1 |
| Number of protein-coding genes | 42856 |
| Complete BUSCO for annotation | 97.1% |
| LAI | 21.62 |

Comparative repeat annotation showed that the enlarged *D. asperoides* genome was largely associated with extensive transposable element accumulation. Repetitive sequences accounted for 91.13% of the genome, with LTR retrotransposons representing 79.27% and Gypsy elements alone comprising 54.27% (Supplementary Table 6). Compared with *T. glandulifera* and *L. japonica*, *D. asperoides* exhibited substantially higher TE and LTR-RT contents (Pu et al., 2020; Zhang et al., 2024), and LTR insertion-time analysis revealed a recent burst of LTR-RT amplification approximately 0.1 million years ago (Supplementary Figure. 10 and Supplementary Tables 7 and 8). These results suggest that recent LTR-RT proliferation was a major contributor to the approximately five-fold larger genome of *D. asperoides* relative to the two related species.

Genome annotation identified 42,856 protein-coding genes, 96.98% of which could be functionally annotated (Supplementary Tables 9-12). We detected 39,197 duplicated genes in *D. asperoides*, exceeding the numbers observed in *T. glandulifera* and *L. japonica* (Pu et al., 2020; Zhang et al., 2024) (Supplementary Figure. 11). Duplication-mode classification indicated substantial contributions from whole-genome, transposed and tandem duplications, with tandemly duplicated genes showing relatively low *Ks* values and elevated *Ka/Ks* ratios, consistent with recent duplication and accelerated sequence divergence (Supplementary Figure. 11). Functional enrichment analysis further showed that rapidly evolving tandem duplicates were enriched in secondary metabolite biosynthetic pathways (Supplementary Figure. 12), suggesting that gene duplication may have contributed to the expansion and diversification of specialized metabolic gene repertoires in *D. asperoides*. We therefore next examined whether triterpenoid saponin-related gene families were specifically expanded and recruited in this enlarged genome.

### Gene duplication expands triterpenoid pathway gene families

Phylogenomic analysis of *D. asperoides* and 15 other plant species showed that the three Caprifoliaceae species clustered into a single clade. Within this clade, Caprifolioideae and Dipsacoideae diverged approximately 58.79 million years ago, whereas *Dipsacus* and *Triplostegia* were recovered as sister genera and diverged approximately 31.45 million years ago (Figure. 2B and Supplementary Figure. 13). Consistent with its enlarged genome, *D. asperoides* contained the higher number of multi-copy genes among the examined species and harbored 810 species-specific gene families (Supplementary Figures. 14 and 15), indicating substantial lineage-specific expansion of its gene repertoire.

**Figure 2.**
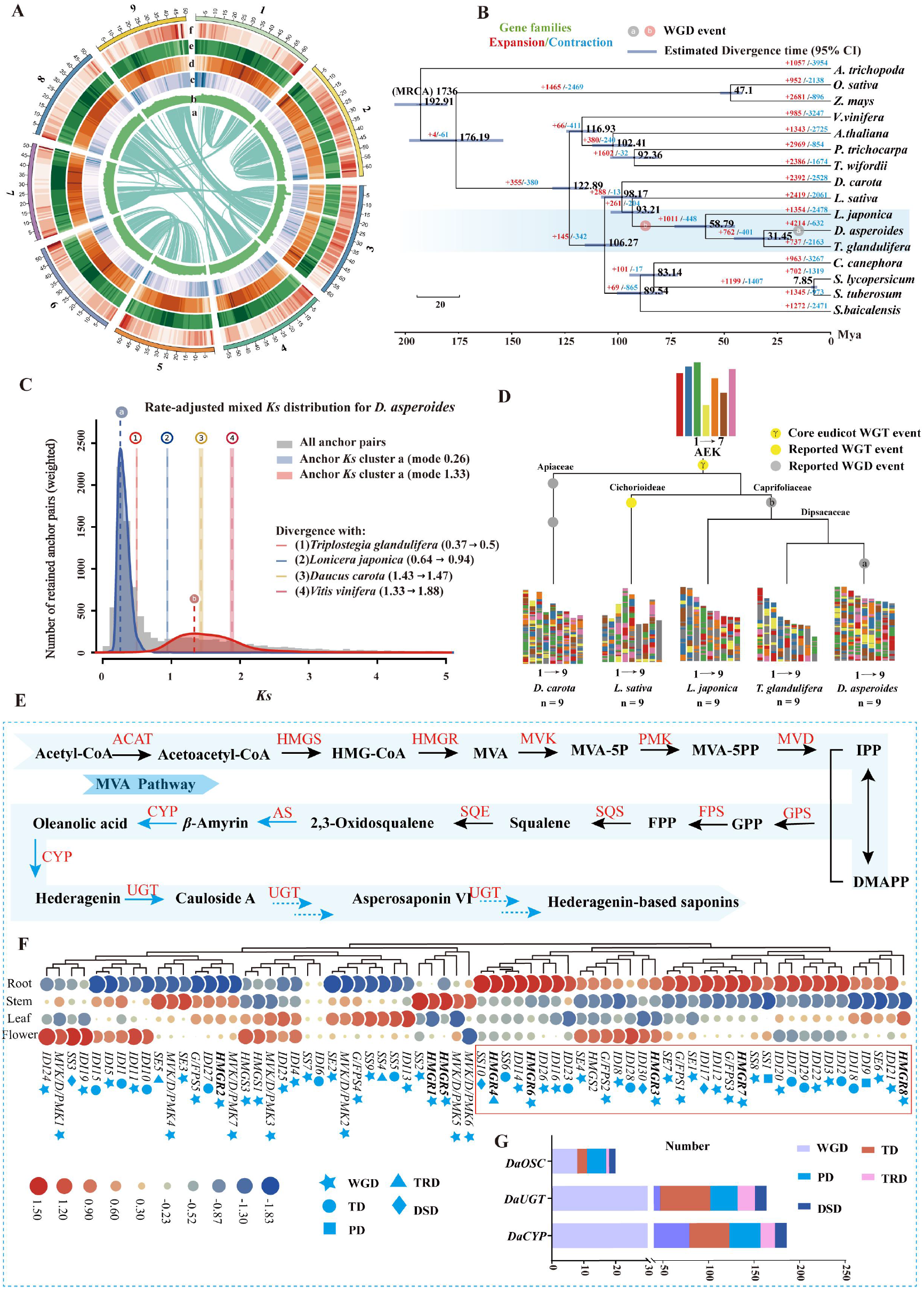
Evolutionary history of the *D. asperoides* genome and the genetic basis for its high asperosaponin VI content. **(A)** Characteristics of the 9 chromosomes of *D. asperoides*. a, Collinearity; b, GC density; c, Copia repetitive sequence density; d, Gypsy repetitive sequence density; e, Repetitive sequence density; f, Gene density. **(B)** Orthologous genes and phylogenetic analysis among *D. asperoides* and 15 other plant species. Blue bars at nodes represent 95% credibility intervals of the estimated dates. The red and cyan numbers represent the expansion and contraction of gene families among 15 other plant species. The ellipse in the branch represents the duplication events and divergence time. **(C)** *K_s_* distributions of anchor pairs for the paralogs of *D. asperoides*, and for the orthologs between *T. glandulifera* and *L. japonica*, and *D. carota* and *V.vinifera* and *A. trichopoda.* Colored dashed lines with letters (a, b, c) represent whole-genome duplication (WGD) events, while those with numbers (1, 2, 3, 4) correspond to the divergence events between *D. asperoides*, *T. glandulifera*, *L. japonica*, *D. carota*, and *V. vinifera*, respectively. **(D)** Chromosomal rearrangements and structural evolutionary patterns from the Ancestral Eudicot Karyotype (AEK, n = 7) to *D. asperoides* and other species. Chromosomes are color-coded to illustrate the evolutionary fragmentation history of the common AEK, which comprises seven proto-chromosomes. **(E)** AACT acetyl-CoA C-acetyltransferase, HMGS hydroxymethylglutaryl-CoA synthase, HMGR hydroxymethylglutaryl-CoA reductase, MVK mevalonate kinase, PMK phosphomevalonate kinase, MVD diphosphomevalonate decarboxylase, IDI isopentenyl-diphosphate Delta-isomerase, GPS geranyl diphosphate synthase, FPS farnesyl diphosphate synthase; SQS squalene synthase, SQE squalene epoxidase, AS amyrin synthase, CYP cytochrome P450 enzymes, UGT UDP-glycosyltransferase. **(F)** Expression profiles and duplication types of genes extracted from expanded gene families involved in the MVA and squalene synthesis pathways. Heatmap shows expression levels across various tissues, with duplication origins indicated on the right. **(G)** Expansion scales and duplication types of key enzyme gene families (OSCs, CYPs, and UGTs) in the *D. asperoides* genome.

Whole-genome duplication events can rapidly expand gene copy numbers and represent important evolutionary events that provide abundant genetic variation for species adaptive evolution (Wang et al., 2022; Yang et al., 2023; Xu et al., 2024; An et al., 2024). Intraspecific and interspecific synteny analysis revealed that *D. asperoides* has undergone multiple whole-genome duplication events (Supplementary Figures. 16 and 17). Syntenic depth ratios of 4:2 between *D. asperoides* and *L. japonica* and 2:1 between *D. asperoides* and *T. glandulifera*, together with two major *Ks* peaks at *Ks* ≈ 1.33 and *Ks* ≈ 0.26, supported an ancestral *Dipsacales*-wide D-WGD event and a more recent *D. asperoides*-specific Da*β* WGD event, estimated to have occurred approximately 91.30 and 17.85 million years ago (Mya), respectively (Figures. 2B-D and Supplementary Figures. 18 and 19). Reconstruction against the Ancestral Eudicot Karyotype further indicated that multiple chromosomal fission and fusion events shaped the current nine chromosomes of *D. asperoides* (Figure. 2D).

Gene family analysis revealed extensive gene family turnover in *D. asperoides*, including 4,214 expanded and 632 contracted gene families (Figure. 2B). Functional enrichment analysis revealed that genes in expanded gene families were primarily enriched for UDP-glycosyltransferase activity and pathways associated with secondary metabolite biosynthesis, whereas genes in contracted gene families were enriched for secondary metabolism-related pathways and cytochrome P450-associated functions (Supplementary Figures. 20-22). We therefore examined gene families involved in the MVA-to-2,3-oxidosqualene pathway and downstream triterpenoid saponin biosynthesis (Figure. 2E). Multiple pathway-related gene families showed expansion, including HMGR and other MVA-pathway genes, as well as genes involved in scaffold formation, oxidative tailoring and glycosylation (Figures. 2F and 2G). Duplication-mode and expression analyses further showed that a subset of duplicated triterpenoid-associated genes was retained and expressed in roots (Figures. 2F and 2G). Together, these results suggest that WGD-associated and local duplications expanded the gene repertoire available for recruitment into triterpenoid saponin biosynthesis in *D. asperoides*.

### Biochemical reconstitution defines the major route to asperosaponin VI

To define the backbone-forming steps of asperosaponin VI biosynthesis, we selected *D. asperoides* OSC candidates from the *β*AS and MAS clades and CYP716A-family candidates using protein length >350 aa and TPM >5 as criteria, and functionally characterized them by heterologous expression in yeast. Beyond the previously reported DaOSC (Yang et al., 2025), two additional *β*AS-clade enzymes, Da*β*AS2 and Da*β*AS3, also produced *β*-amyrin, whereas three MAS-clade enzymes, DaMAS1-3, produced both *β*-amyrin and *α*-amyrin (Figures. 3A and 3B and Supplementary Figures. 23-24). In addition, three CYP716 enzymes, DaOAS1-3, catalysed sequential *C*-28 oxidation of *β*-amyrin to produce oleanolic acid (4) (Figures. 3A and 3C and Supplementary Figures. 25-26). Together with the previously identified *C*-23 oxidases DaCYP714E29/DaCYP714E30 (Das09G003295.t1/Das03G003375.t1) (Yang et al., 2025), these results define the core backbone-forming module for hederagenin-based saponin biosynthesis in *D. asperoides*.

**Figure 3.**
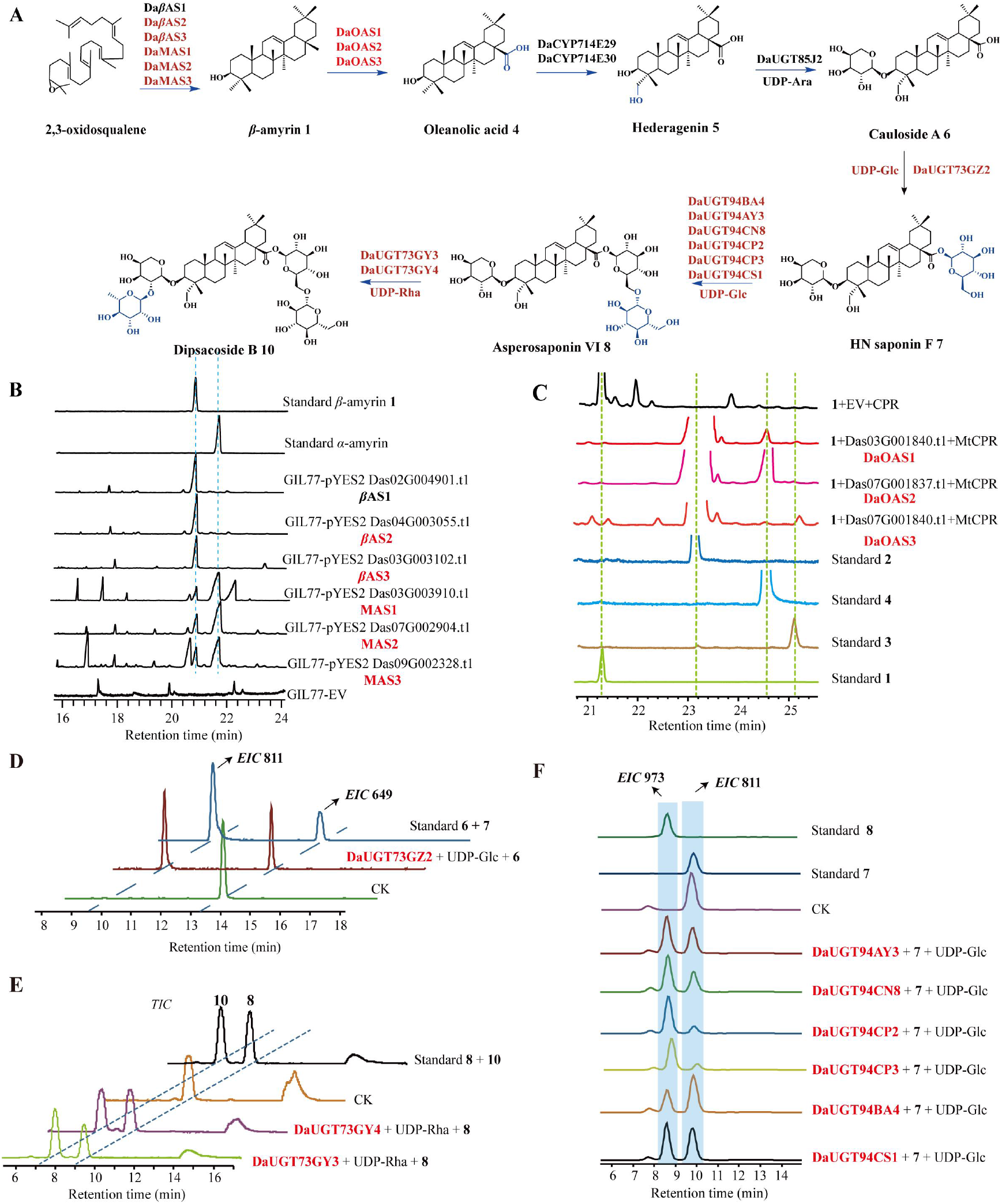
Elucidation of the biosynthetic pathway from 2,3-oxidosqualene to asperosaponin VI and dipsacoside B in *D. asperoides*. **(A)** Biosynthetic pathway from 2,3-oxidosqualene to Dipsacoside B (**10**). Enzyme names shown in red indicate enzymes functionally characterized in this study, whereas those shown in black indicate enzymes characterized previously. **(B)** GC-MS chromatograms of yeast extracts expressing candidate *D. asperoides* OSC genes (*DaβAS*1-3 and DaMAS1-3). **(C)** GC-MS EICs of yeast strains expressing DaOAS1-3, showing the production of intermediates erythrodiol (**2**), oleanolic aldehyde (**3**), and oleanolic acid (4). **(D)** Reaction scheme and LC-MS validation (*EIC* 649/811) of DaUGT73GZ2 converting cauloside A (**6**) to HN saponin F (**7**). **(E)** Reaction scheme and LC-MS validation of DaUGT73GY3/DaUGT73GY4 catalyzing the formation of dipsacoside B (**10**) from asperosaponin VI (**8**). **(F)** Reaction scheme and LC-MS validation (*EIC* 973/811) of various DaGlcGT members (DaUGT94AY3, Da UGT94CN8, DaUGT94CP2, DaUGT94CP3, DaUGT94BA4, DaUGT94CS1) converting HN saponin F (**7**) to asperosaponin VI (**8**).

To identify genes involved in the downstream glycosylation of hederagenin-based saponins in *D. asperoides*, we performed co-expression analysis using *β*AS, CYP714E29 and DaUGT85J2 as bait genes and identified 13 candidate UGTs (Supplementary Figure. 27). UGT73 and UGT94 families are frequently associated with triterpenoid saponin glycosylation. Therefore, we selected members of the UGT73 and UGT94 families for further functional characterization (Supplementary Figures. 28 and 29). In vitro assays showed that the UGT73 enzyme DaUGT73GZ2 catalysed the conversion of **6** to **7** (Figure. 3D and Supplementary Figures. 30 and 31). Two additional UGT73 enzymes, DaUGT73GY4 and DaUGT73GY3, rhamnosylated **6** and **8** to produce **9** and **10**, respectively (Figure. 3E and Supplementary Figures. 30, 32 and 33). By contrast, the UGT94 enzymes DaUGT94BA4 and DaUGT94CP3 catalyzed the subsequent conversion of **7** to **8** (Figure. 3F and Supplementary Figures. 30 and 34). We further cloned and assayed additional members of the UGT94 family, among which four enzymes, DaUGT94CP2, DaUGT94CS1, DaUGT94AY3 and DaUGT94CN8, also catalyzed the conversion of **7** to **8** (Figure. 3F and Supplementary Figures. 29 and 34). Finally, transient co-expression of the six most highly expressed *D. asperoides* pathway genes-Da*β*AS2, DaOAS2, DaCYP714E29, DaUGT85J2, DaUGT73GZ2 and DaUGT94BA4-with oat tHMGR in *Nicotiana benthamiana* leaves resulted in detectable asperosaponin VI production, supporting pathway reconstruction in planta (Supplementary Figure. 35).

### *C*-3 glycosylation primes DaUGT73GZ2-mediated *C*-28 glucosylation in asperosaponin VI biosynthesis

Having identified DaUGT73GZ2 as the enzyme converting cauloside A (**6**) to HN saponin F (**7**), we next asked whether this reaction imposed directionality on asperosaponin VI glycosylation. Kinetic analysis using **6** and UDP-Glc as substrate showed that DaUGT73GZ2 catalyzed *C*-28 glucosylation with a catalytic efficiency of 2.94 x10^-4^ μM^-^¹ s^-1^ (Supplementary Figure. 36), and donor-selectivity assays indicated that UDP-Glc was the preferred sugar donor (Supplementary Figure. 37). We then compared its activity toward a panel of 19 pentacyclic triterpenoids that differed in *C*-3 and *C*-28 substitution status. DaUGT73GZ2 accepted only substrates containing both a *C*-3 glycosyl group and a free *C*-28 carboxyl group, including **6**, **9**, **20**–**21** and **23**–**27**, whereas aglycones and *C*-28-substituted saponins were not detectably glycosylated (Figure. 4A-B and Supplementary Figures. 38–45 and Supplementary Table 13). Thus, prior *C*-3 glycosylation appears to license DaUGT73GZ2-mediated *C*-28 glycosylation, establishing a *C*-3-before-*C*-28 glycosylation sequence that directs intermediates toward asperosaponin VI formation.

**Figure 4.**
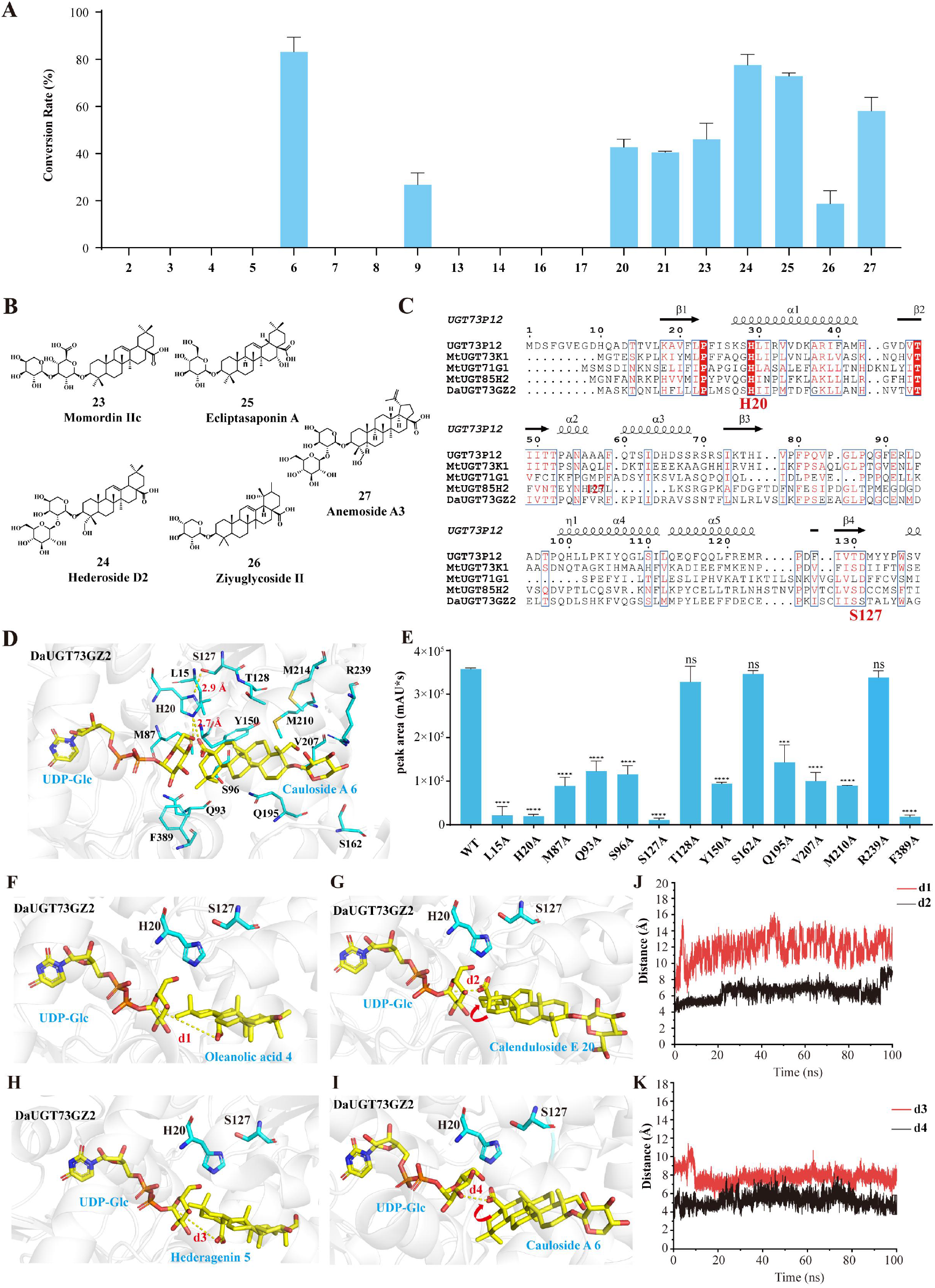
Biochemical substrate preference and structural model of DaUGT73GZ2. **(A)** Conversion rates (%) of glycosylated products from 19 substrates with different C-3 and C-28 substitution patterns using UDP-Glc as the sugar donor. **(B)** Chemical structures of substrates **23**–**27** used to assess substrate promiscuity. **(C)** Multiple sequence alignment of DaUGT73GZ2 with structurally characterized plant UGTs. **(D)** Molecular docking models of DaUGT73GZ2 complexed with cauloside A (6) and UDP-Glc. Both substrates are highlighted in yellow, with yellow dashed lines indicating hydrogen bonds. Key amino acid residues within a 4 Å radius of the substrates in the binding pocket are displayed. **(E)** Alanine scanning mutagenesis. Key residues within a 4 Å radius of Cauloside A were mutated to alanine. Data are presented as mean ± SD (n = 3) from three biological replications. Statistical significance was determined using Student’s t-test. ns, not significant, ***P < 0.001, \*\*\*\**P* < 0.0001. (**F–I)** Molecular docking models of DaUGT73GZ2 with UDP-Glc and different acceptor substrates, including oleanolic acid, Calenduloside E, hederagenin, and cauloside A. H20 is shown as the key catalytic residue. Yellow dashed lines indicate the distances between the *C*-28 carboxyl oxygen atom of the acceptor substrate and the anomeric carbon of UDP-Glc, denoted as d1–d4. **(J-K)** Distance changes of d1–d4 during 100-ns molecular dynamics simulations.

To explore the structural basis for the preference of DaUGT73GZ2 toward *C*-3-glycosylated substrates, we combined sequence alignment, molecular docking, site-directed mutagenesis and molecular dynamics simulations. Unlike many plant UGTs that contain a canonical His-Asp catalytic dyad, sequence alignment identified a non-canonical H20-S127 arrangement in DaUGT73GZ2 (Yao et al., 2022) (Figure. 4C). Docking of DaUGT73GZ2 with UDP-Glc and cauloside A (**6**) placed the *C*-28 carboxyl group of 6 near both H20 and the anomeric carbon of UDP-Glc, with H20 forming hydrogen bonds with S127 and the *C*-28 carboxyl group (2.9 Å and 2.7 Å, respectively; Figure. 4D). Consistently, H20A and S127A variants showed markedly reduced activity, and mutations of several residues surrounding the acceptor pocket also decreased enzymatic activity (Figure. 4E).

Comparative docking further showed that, within the DaUGT73GZ2 active-site pocket, the *C*-28 carboxyl groups of aglycones **4** and **5** failed to adopt a favourable orientation toward the catalytic residue H20 and the anomeric carbon of UDP-Glc. By contrast, their *C*-3-glycosylated derivatives **20** and **6** displayed an approximately 120° reorientation of the *C*-28 carboxyl group, placing it in a catalytically competent conformation relative to H20 and the UDP-Glc anomeric carbon (Figure. 4F-I). Other *C*-3-glycosylated substrates, including **9** and **23–27**, similarly adopted catalytically competent orientations, whereas aglycones **15** and **16** remained unfavourably oriented (Supplementary Figure. 46). Molecular dynamics simulations supported this distinction: the *C*-28 carboxyl groups of aglycones **4** and **5** remained farther from both H20 and the UDP-Glc anomeric carbon, whereas those of the *C*-3-glycosylated substrates **20** and **6** maintained shorter distances to the two reaction-relevant positions throughout the simulation (Figure. 4J-K and Supplementary Figures. 47 and 48). Together, these structural analyses suggest a model in which *C*-3 glycosylation may favor positioning of the *C*-28 carboxyl group near H20 and the UDP-Glc anomeric carbon, providing a plausible structural basis for the preference of DaUGT73GZ2 toward *C*-3-glycosylated substrates.

### Evolutionary and functional diversification of UGT85CA enzymes in Caprifoliaceae

*C*-3 arabinosylation is a characteristic modification of hederagenin-based saponins in Caprifoliaceae, but the evolutionary origin of the glycosyltransferase activity responsible for this modification remains unclear (Liu et al., 2011; Yu et al., 2019; Yin et al., 2023). We previously identified DaUGT85J2 as a *C*-3 arabinosyltransferase acting on hederagenin in *D. asperoides* (Yang et al., 2025); this enzyme was subsequently designated DaUGT85CA7 by the UGT Nomenclature Committee. Phylogenetic analysis of UGTs from 16 angiosperm species showed that UGT85CA homologs from the three sampled Caprifoliaceae species formed a distinct lineage containing DaUGT85CA7 (Figure. 5A and Supplementary Figure. 49 and Supplementary Table 14). Branch-wise *Ka/Ks* analysis further showed that most branches within the UGT85CA lineage had *Ka/Ks* ratios below 1, indicating that the lineage has been predominantly subject to purifying selection. To further examine the taxonomic distribution of UGT85CA, we searched the NCBI non-redundant protein and OneKP databases using stringent criteria (E-value ≤ 1e^−5^ and protein identity > 60%). No homologs satisfying these criteria were identified outside Caprifoliaceae, suggesting that UGT85CA represents a lineage largely restricted to Caprifoliaceae and has undergone lineage-specific diversification within the family.

**Figure 5.**
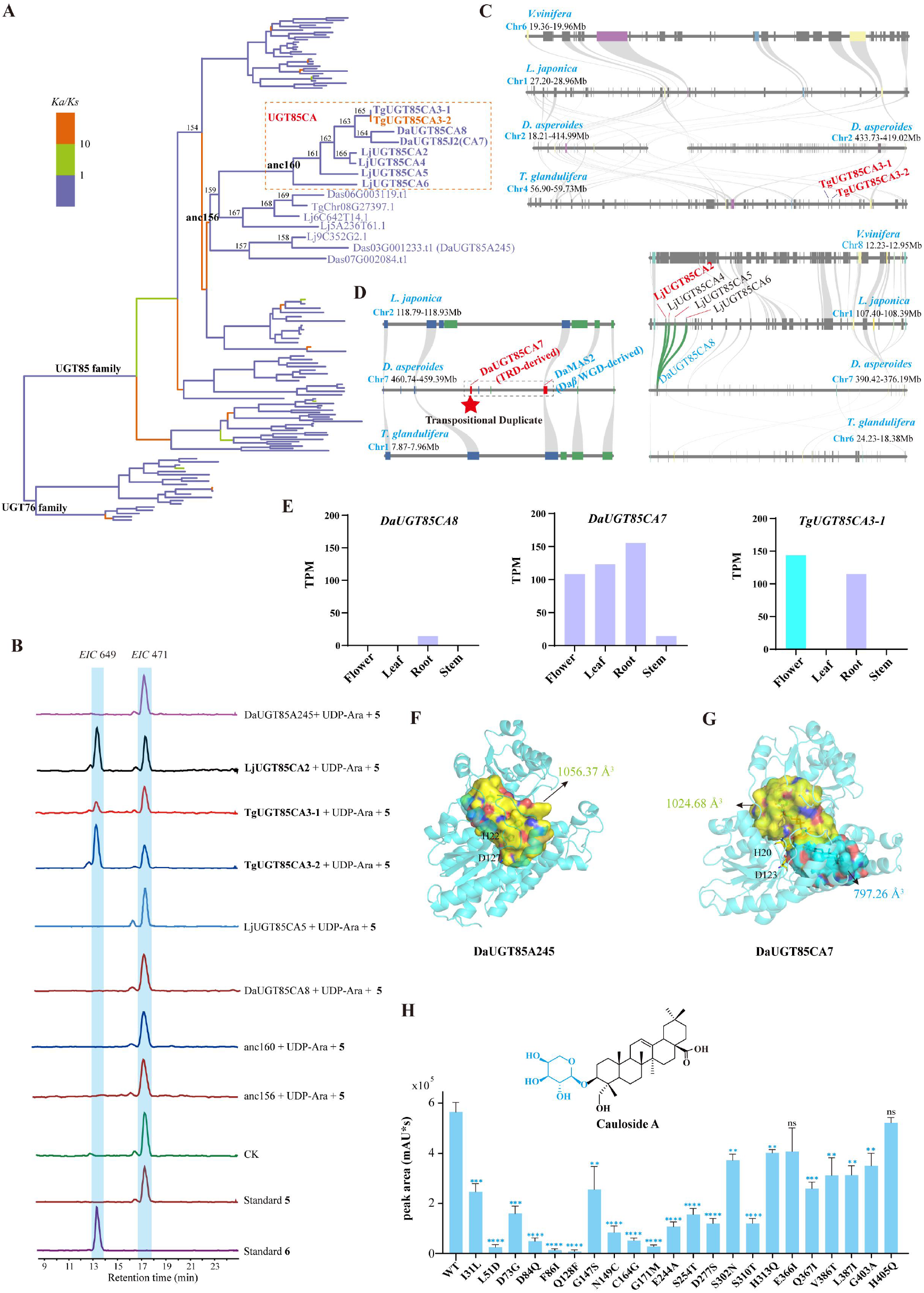
Evolutionary origin, genomic recruitment and functional specialization of *C*-3 arabinosyltransferase activity in the Caprifoliaceae UGT85CA lineage. **(A)** Phylogenetic and selection pressure analysis of the UGT85 family. *Ka/Ks*, nonsynonymous (*Ka*) with synonymous (*Ks*) substitution rates. **(B)** Extracted ion chromatograms (EIC) of compounds formed in enzymatic reactions of candidate UGTs with two substrates: UDP-Ara and hederagenin. Standard **5** and standard **6** represent authentic standards of hederagenin and cauloside A, respectively. **(C-D)** Synteny analysis of the UGT85CA subfamily in *T. glandulifera*, *D. asperoides*, and *L. japonica*. (**E)** Expression levels of *DaUGT85CA8*, *DaUGT85CA7*, and *TgUGT85CA3-1* across different tissues. (**F-G)** Predicted active-site pockets of DaUGT85A245 and DaUGT85CA7. The sugar-donor pocket is shown in yellow, and the sugar-acceptor pocket is shown in cyan. **(H)** Catalytic activities of DaUGT85J2 and its 20 site-directed mutants. Data are presented as mean ± SD (n = 3) from three biological replications. Statistical significance was determined using Student’s t-test. ns, not significant, **P < 0.01, \*\*\**P* < 0.001. ****, P < 0.0001.

We next investigated the functional diversification of the UGT85CA lineage. Full-length cDNAs were successfully obtained for five UGT85CA genes, whereas the remaining homologs exhibited extremely low transcript abundance in the sampled tissues based on transcriptome data and could not be recovered as complete cDNAs. Biochemical characterization showed that TgUGT85CA3-1, TgUGT85CA3-2, and LjUGT85CA2 exhibited *C*-3 arabinosyltransferase activity, whereas no corresponding activity was detected for DaUGT85CA8, LjUGT85CA5, Anc160, or Anc156 under the same assay conditions (Figure. 5B and Supplementary Figures. 50–52). Together with the previously characterized activity of DaUGT85CA7, *C*-3 AraGT activity showed a discontinuous distribution among the extant UGT85CA enzymes examined and was not detected in either reconstructed ancestral protein. This pattern is consistent with lineage-specific acquisition of *C*-3 AraGT activity in multiple UGT85CA branches and suggests possible parallel functional evolution within the lineage.

Comparative synteny and expression analyses revealed lineage-specific diversification in the genomic organization and transcriptional profiles of UGT85CA genes across Caprifoliaceae (Figures. 5C-E). UGT85CA genes in *L. japonica* and the syntenic DaUGT85CA8 copy in *D. asperoides* generally showed low transcript abundance. TgUGT85CA3-1 and TgUGT85CA3-2 retained orthologous relationships with the *L. japonica* UGT85CA genes but were located in non-syntenic genomic regions, indicating lineage-specific changes in genomic context; among them, TgUGT85CA3-1 showed relatively high expression. In contrast, DaUGT85CA7 lacked a clear orthologous or syntenic counterpart in the other examined species and was located in a TE-rich genomic region, where it was classified as a transposed duplicate (Supplementary Figures. 53 and 54 and Supplementary Table 15). DaUGT85CA7 is located adjacent to DaMAS2, an OSC gene involved in triterpenoid scaffold formation, forming a *D. asperoides*-specific adjacent pair of pathway-related genes (Figure. 5D). Notably, DaUGT85CA7 showed high transcript abundance, with its highest expression detected in roots (Figure. 5E). Together, these results indicate substantial lineage-specific genomic and transcriptional diversification of UGT85CA genes, with the functional DaUGT85CA7 representing a distinctive, highly expressed transposed copy in *D. asperoides*.

Most characterized UGT85 enzymes preferentially act on relatively small-molecule acceptors, whereas activity toward bulky pentacyclic triterpenoids is uncommon (Kurze et al., 2022; Del Giudice et al., 2022). To investigate the structural basis underlying the unusual pentacyclic triterpenoid glycosyltransferase activity of DaUGT85CA7, we compared it with DaUGT85A245, its closest non-UGT85CA homolog in *D. asperoides* (Supplementary Figure. 55). Structural modelling predicted that both proteins contain conserved sugar-donor-binding pockets, whereas DaUGT85CA7 possesses an additional spacious *N*-terminal acceptor pocket that providing sufficient space to accommodate bulky pentacyclic triterpenoid substrates (Figure. 5F and 5G). Similar acceptor-pocket features were also predicted in the biochemically characterized active Caprifoliaceae UGT85CA enzymes, including TgUGT85CA3-1 and LjUGT85CA2 (Supplementary Figure. 56). Sequence comparison identified 24 conserved but divergent residues potentially associated with the formation and remodeling of this acceptor pocket (Supplementary Figure. 57). Systematic substitution of these residues in DaUGT85CA7 revealed that several residues located within the predicted *N*-terminal acceptor pocket region, particularly L51, F86, Q128, and G171, markedly reduced product formation when substituted (Figure. 5H). These results support the importance of *N*-terminal acceptor-pocket residues in the recognition and catalytic transformation of bulky pentacyclic triterpenoid substrates by DaUGT85CA7. Furthermore, the structural model positioned the conserved catalytic residue H21 within this acceptor pocket, where it could form a hydrogen bond with the *C*-3 hydroxyl group of hederagenin, while D123 was located nearby. Alanine substitution of either residue nearly abolished DaUGT85CA7 activity, providing functional support for the predicted active-site configuration and substrate-binding model (Supplementary Figure. 58). Together, these results indicate that the expanded N-terminal acceptor pocket and its associated residues are closely linked to the specialization of DaUGT85CA7 toward bulky pentacyclic triterpenoid substrates.

### Pathway-gene expansion and root-biased expression support asperosaponin VI accumulation

Comparative genomic analyses showed that several gene families contributing to hederagenin-based saponin biosynthesis were expanded in *D. asperoides* relative to closely related species. The *β*AS, MAS and CYP714E families showed lineage-specific expansion, whereas the CYP716A family was contracted overall (Supplementary Figure. 59). Notably, however, the functionally characterized CYP716A copies responsible for *C*-28 oxidation were retained and amplified through Da*β* WGD and proximal duplication, indicating retention and further duplication of pathway-relevant oxidases despite contraction of the broader CYP716A family. Duplication-mode, collinearity and *Ks* analyses further showed that *β*AS and MAS expansions were mainly associated with the Da*β* WGD event in the *D. asperoides* lineage, whereas CYP genes involved in *C*-28 and *C*-23 oxidation originated through both Da*β* WGD and proximal duplication (Figure. 6A; Supplementary Figures. 60-62 and Supplementary Tables 16-17). Downstream glycosylation genes were also expanded: UGT94 and UGT73 family members involved in hederagenin-based saponin glycosylation were amplified through Da*β* WGD-derived syntenic blocks and tandem duplication, forming local clusters on chromosomes 3 and 9 (Figure. 6B, C and Supplementary Figures. 63-66). In particular, a 291-kb UGT94 cluster on chromosome 9 indicates extensive tandem amplification following WGD.

**Figure 6.**
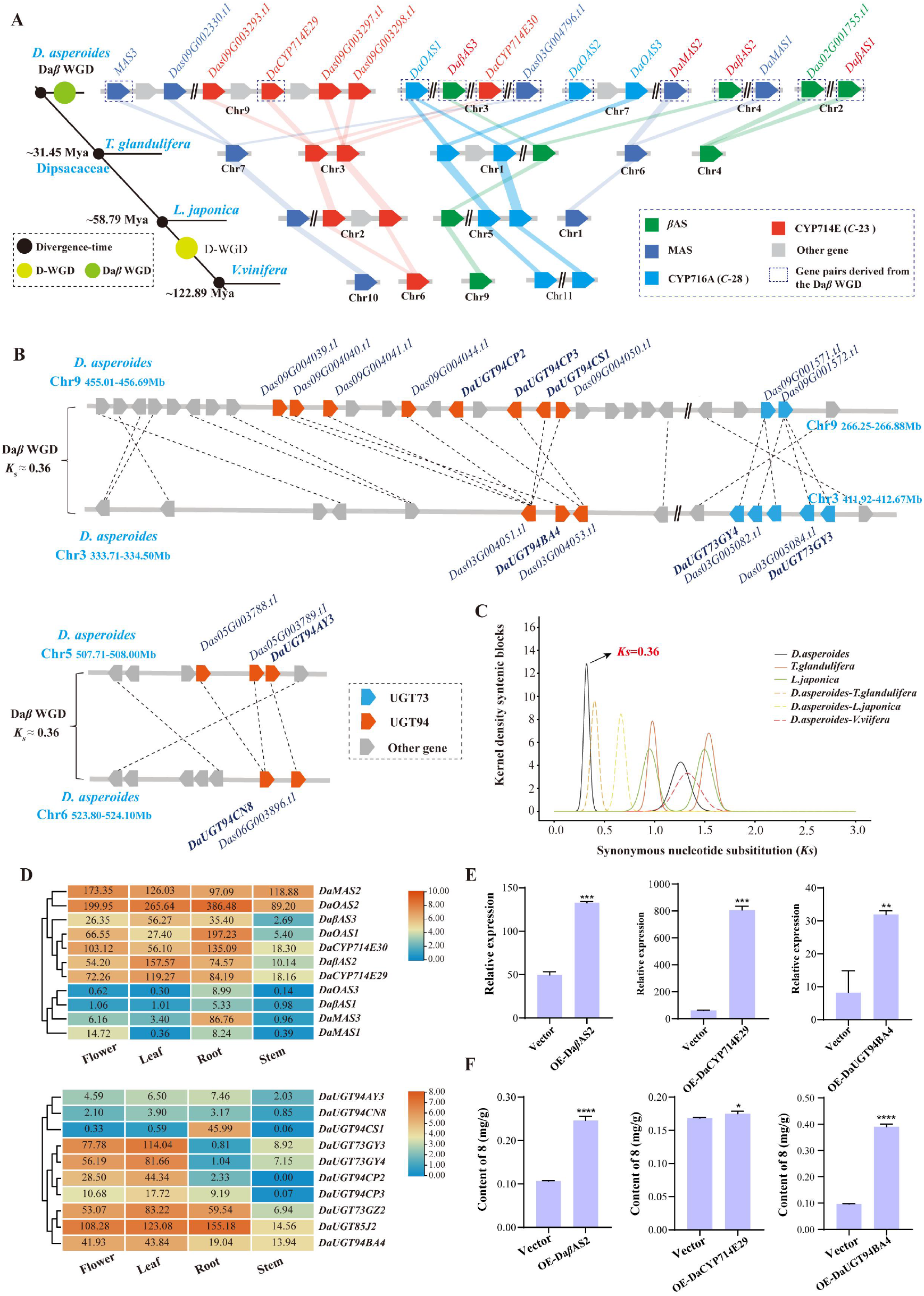
Evolutionary expansion and transcriptional patterns of saponin biosynthetic genes in *D. asperoides*. **(A)** Microsynteny analysis of genes involved in triterpenoid backbone biosynthesis across *D. asperoides*. **(B)** Synteny analysis of UGTs involved in hederagenin-based saponin biosynthesis in *D. asperoides*. The UGT genes responsible for the hederagenin-type saponin pathway are highlighted in red, with dashed lines indicating their syntenic relationships. **(C)** Distribution of synonymous substitution rates (*Ks*) for orthologous gene pairs. The *Ks* values among *D. asperoides*, *T. glandulifera*, *L. japonica*, *D. carota*, and *V. vinifera* were calculated using the WGDI tool. **(D)** Tissue-specific expression profiles of key biosynthetic genes. **(E)** qRT-PCR analysis of ***DaβAS2***, ***DaCYP714E29*** and ***DaUGT94BA4*** transcript levels in *D. asperoides* leaves transiently overexpressing the corresponding pathway genes. **(F)** LC-MS quantification of asperosaponin VI (**8**) content in *D. asperoides* leaves transiently overexpressing ***DaβAS2***, ***DaCYP714E29*** or ***DaUGT94BA4***. Data are presented as mean ± SD (n = 3) from three biological replications. Statistical significance was determined using Student’s t-test. *P < 0.05, **P < 0.01, \*\*\**P* < 0.001. ****, P < 0.0001.

Beyond copy-number expansion, expression profiling showed that the functionally characterized genes involved in asperosaponin VI biosynthesis were among the most highly expressed members of their respective subfamilies in *D. asperoides*, particularly in roots (Figure. 6D and Supplementary Figures. 67-69). This expression pattern suggests that duplicated OSC, CYP and UGT copies were not only retained, but also transcriptionally recruited into a root-active hederagenin-based saponin pathway. By contrast, the downstream rhamnosyltransferases DaUGT73GY4/DaUGT73GY3, which convert asperosaponin VI to dipsacoside B, showed very low expression in roots (Figure. 6D). This expression pattern is consistent with limited downstream conversion of asperosaponin VI and may contribute to its predominant accumulation in roots.

To further test whether increased expression of representative pathway genes could promote asperosaponin VI accumulation, Da*β*AS2, DaCYP714E29 and DaUGT94BA4 were transiently overexpressed in *D. asperoides* leaves. qRT-PCR analysis confirmed that the transcript levels of the corresponding genes were significantly increased in overexpression tissues compared with empty-vector controls (Figure. 6E). Quantitative analysis further showed that transient overexpression of Da*β*AS2, DaCYP714E29 or DaUGT94BA4 significantly increased the asperosaponin VI content, with the largest increase observed in leaves overexpressing DaUGT94BA4 (Figure. 6F and Supplementary Figure. 70). These results show that increased expression of individual pathway genes can enhance asperosaponin VI accumulation in the transient leaf expression system. Together, the observed patterns of gene duplication, root-biased expression and pathway-gene activity are associated with the high-abundance asperosaponin VI profile of *D. asperoides* roots.

## DISCUSSION

Specialized metabolic pathways often diversify among related plant lineages, but how plants organize shared precursor pools toward high-abundance, organ-dominant metabolic outputs remains poorly understood. In this study, we show that the dominant accumulation of asperosaponin VI in *D. asperoides* roots is associated with a multilayered pathway architecture involving precursor loading, ordered glycosylation and downstream product retention. Genome expansion and pathway-gene duplication enlarged the triterpenoid biosynthetic gene pool, whereas root-biased expression of functional OSC, CYP and UGT copies supported the production of hederagenin-based intermediates. Downstream, DaUGT85J2-mediated *C*-3 arabinosylation and DaUGT73GZ2-mediated *C*-28 glucosylation established an ordered glycosylation sequence that favors asperosaponin VI formation, while low root expression of DaUGT73GY4/ DaUGT73GY3 is consistent with reduced conversion to downstream rhamnosylated products. Together, these findings suggest that *D. asperoides* evolved a high-abundance triterpenoid metabolic profile through the integration of genomic expansion, enzymatic gating and transcriptional restriction.

Genome expansion in *D. asperoides* appears to have provided an enlarged evolutionary substrate for triterpenoid pathway assembly rather than acting as a direct explanation for asperosaponin VI accumulation. Gene duplication exerts vital functions in organisms’ responses to environmental stress through gene dosage effect, subfunctionalization, neofunctionalization and pseudogenization (Panchy et al., 2016; Peng et al., 2024; Wang et al., 2026). The recent LTR-associated genome expansion and the *D. asperoides*-specific Da*β* WGD generated a large duplicated gene repertoire, including multiple OSC, CYP and UGT copies associated with hederagenin-based saponin biosynthesis. However, only a subset of these duplicates was retained, expressed and functionally recruited into the asperosaponin VI pathway. This is illustrated by the CYP716A family: although the family contracted overall, *C*-28 oxidases relevant to hederagenin formation were preserved and amplified. Thus, genome expansion likely increased the available gene pool, whereas pathway-specific retention and transcriptional recruitment determined which duplicated copies contributed to the root-active triterpenoid pathway.

*C*-3 arabinosylation represents a key early glycosylation step in hederagenin-based saponin biosynthesis. Functional analyses of the sampled UGT85CA enzymes revealed a discontinuous distribution of *C*-3 arabinosyltransferase activity, indicating substantial functional diversification within this lineage and raising the possibility that this activity was acquired independently in different branches. However, because only a limited number of extant UGT85CA enzymes and reconstructed ancestral proteins were functionally characterized, the current data cannot distinguish repeated independent gains from an alternative scenario in which *C*-3 AraGT activity arose earlier and was subsequently lost in some branches. Broader taxon sampling and functional characterization of additional extant and ancestral enzymes will therefore be required to resolve the evolutionary history of this activity. The occurrence of analogous *C*-3 arabinosyltransferase activity in the phylogenetically distinct oat UGT99D1 further suggests that similar glycosylation functions can evolve in different UGT lineages during triterpenoid saponin evolution (Louveau et al., 2018) (Supplementary Figure. 71). Beyond this functional diversification, DaUGT85CA7 in D. asperoides also occurs in a distinct genomic context. It is located adjacent to MAS2, a triterpenoid-related OSC gene, and shows relatively high expression in roots. This genomic arrangement illustrates the reorganization accompanying UGT85CA diversification in *D. asperoides*, although the functional or regulatory significance of the proximity between DaUGT85CA7 and MAS2 remains to be established.

DaUGT73GZ2 provides a biochemical basis for ordered glycosylation during asperosaponin VI biosynthesis. In contrast to the broad substrate promiscuity reported for some triterpenoid UGTs, DaUGT73GZ2 preferentially accepts substrates that are already glycosylated at *C*-3 and retain a free *C*-28 carboxyl group. This substrate preference favors *C*-28 glucosylation after DaUGT85CA7-mediated *C*-3 arabinosylation, thereby limiting *C*-28 glucosylation of aglycones and establishing a *C*-3-before-*C*-28 reaction sequence. Structural modelling and mutagenesis further suggest that the non-canonical H20–S127 arrangement and surrounding acceptor-pocket residues may contribute to this substrate preference by favoring catalytically competent orientations of *C*-3-glycosylated substrates. Thus, the ordered glycosylation of asperosaponin VI is encoded, at least in part, by the substrate preference of DaUGT73GZ2, with its active-site architecture providing a plausible structural basis for this selectivity.

Transcriptional regulation may represent an additional layer associated with asperosaponin VI accumulation in roots. Functional OSC, CYP and UGT genes involved in asperosaponin VI biosynthesis show relatively high expression in roots, whereas the downstream rhamnosyltransferases DaUGT73GY3 and DaUGT73GY4 are expressed at very low levels. This expression pattern is consistent with limited downstream conversion of asperosaponin VI and may represent one factor contributing to its predominant accumulation in *D. asperoides* roots. In addition, the low expression of syntenic pathway orthologs in *L. japonica* and the apparent enrichment of oleanolic acid-type saponins in *T. glandulifera* suggest that the sampled Caprifoliaceae species differ in their triterpenoid metabolic profiles, potentially reflecting distinct combinations of enzyme activity and transcriptional deployment (Supplementary Figures. S72-75).

In summary, our study supports a multilayered model associated with the predominant accumulation of asperosaponin VI in *D. asperoides* roots. LTR proliferation contributed substantially to genome expansion, whereas WGD and local gene duplications expanded the repertoire of triterpenoid-associated genes, from which functional OSC, CYP and UGT copies were retained and expressed in roots. DaUGT85CA7 mediates *C*-3 arabinosylation of hederagenin-type aglycones, whereas the substrate preference of DaUGT73GZ2 favors a *C*-3-before-*C*-28 glycosylation sequence during asperosaponin VI biosynthesis. In contrast, the low root expression of the downstream rhamnosyltransferases DaUGT73GY3 and DaUGT73GY4 is consistent with limited conversion of asperosaponin VI to downstream rhamnosylated products and may contribute to its predominant accumulation in roots. Together, these genomic, enzymatic and transcriptional features provide an integrated framework for understanding the distinctive, high-abundance triterpenoid saponin profile of *D. asperoides* (Figure. 7). Nevertheless, direct genetic perturbation of key pathway genes will be required to establish the causal contributions of gene duplication and transcriptional regulation to asperosaponin VI accumulation in planta.

**Figure 7.**
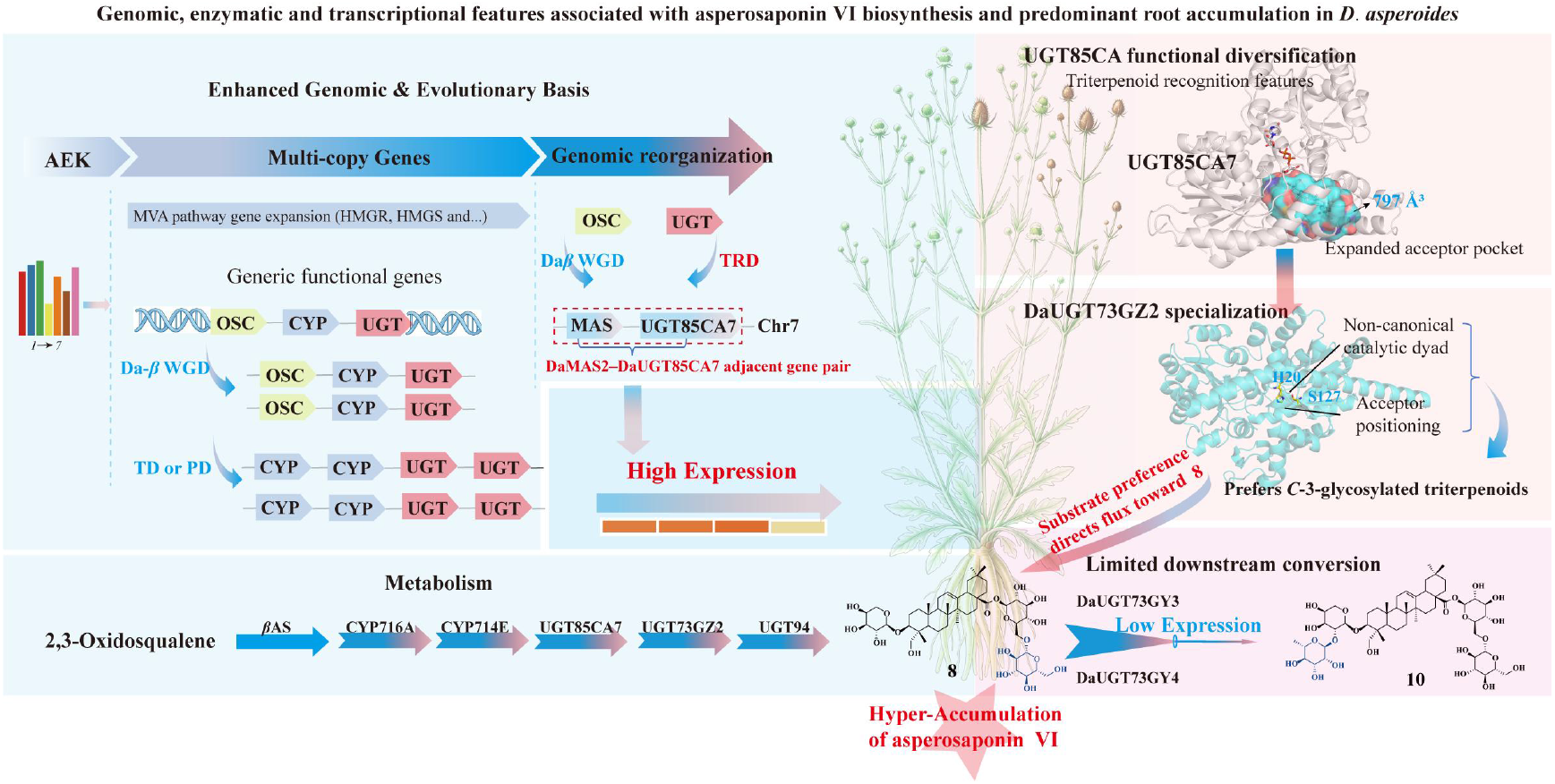
Integrated model of genomic, enzymatic and transcriptional features associated with asperosaponin VI biosynthesis and predominant root accumulation in *D. asperoides*. WGD and local duplication events expanded the repertoire of triterpenoid-associated genes in *D. asperoides*. DaUGT85CA7 represents a transposed UGT85CA copy in a distinct genomic context and catalyzes *C*-3 arabinosylation during hederagenin-based saponin biosynthesis. DaUGT73GZ2 preferentially accepts *C*-3-glycosylated substrates retaining a free *C*-28 carboxyl group, supporting a *C*-3-before-*C*-28 glycosylation sequence. Low root expression of DaUGT73GY3 and DaUGT73GY4 is consistent with limited downstream conversion of asperosaponin VI. Together, these genomic, enzymatic and transcriptional features are associated with the predominant accumulation of asperosaponin VI in *D. asperoides* roots.

## METHODS

### Plant materials and chemical reference standards

*D. asperoides* plants were obtained from a cultivated base in Baoshan City, Yunnan Province, China. Fresh leaves were harvested, immediately frozen in liquid nitrogen, and stored at −80 °C for genome sequencing. Roots, stems, leaves, and flowers of *D. asperoides*, *T. glandulifera*, and *L. japonica* were collected for the detection of triterpenoids. All chemical standards were purchased from BioBioPha Co., Ltd.

### LC–MS analysis of metabolites

All samples (in triplicate) were dried in an oven at 50 °C and ground into fine powder. Subsequently, 0.5 g of the powder was weighed into a 50-mL centrifuge tube, followed by the addition of 30 mL of methanol. The mixture was subjected to ultrasonic extraction (240 W, 40 kHz) for 30 minutes. After extraction, 1 mL of the clarified supernatant was collected, filtered through a 0.22-μm membrane, and transferred into a 1.5-mL vial for subsequent analysis.

Samples were analyzed using an Agilent 1290 UPLC system coupled with a 6540 Q-TOF mass spectrometer equipped with an Agilent C18 column (5 μm, 4.6 × 250 mm). The mobile phase consisted of water containing 0.01% formic acid (A) and acetonitrile (B), with the following linear gradient program: 0–9.5 min, 30% B; 9.5–13.5 min, 30% to 88% B; 13.5–25 min, 88% B. The flow rate was 0.8 mL/min and the column temperature was maintained at 30 °C. Mass spectrometry was performed in negative ion mode with the following settings: capillary voltage, 3500 V; fragmentor voltage, 135 V; skimmer voltage, 60 V; octopole RF voltage, 750 V; and scan range, 50–1700 m/z.

### Genome sequencing, survey, and assembly

High-molecular-weight genomic DNA (gDNA) was extracted from fresh leaves of *D. asperoides* using modified CTAB and SDS protocols, respectively, and purified using the Qiagen Genomic DNA Kit. The quality and concentration of the extracted gDNA were assessed using a NanoDrop One spectrophotometer and a Qubit 3.0 Fluorometer, respectively. Genome survey sequencing was performed on the MGI DNBSEQ-T7 platform. A PCR-free SMRTbell library was constructed from high-quality long-fragment DNA and sequenced using PacBio HiFi technology. Hi-C library preparation and sequencing were conducted on the BGI platform.

All second-generation sequencing data (including Survey and Hi-C data) were filtered using Trimmomatic v0.39 (Bolger et al., 2014). K-mer frequency analysis (k = 17) was performed with Jellyfish v2.3.1 (Marçais and Kingsford, 2011), and genome characteristics including genome size, heterozygosity, and repeat content were estimated using GCE v1.0.2 (Wang et al., 2020). Ploidy level was determined with Smudgeplot v0.2.5 (Ranallo-Benavidez et al., 2020). De novo genome assembly was conducted from PacBio HiFi reads using Hifiasm v0.21.0 (Cheng et al., 2021). Organellar genomes were assembled using Oatk v1.0 (Zhou et al., 2024). Genomic reads were aligned to chloroplast and mitochondrial sequences via Minimap2 v2.28 (Li, 2021), and sequences with over 90% coverage were removed. Subsequent realignment against third-generation sequencing data facilitated the filtering of low-depth regions. Redundant haplotigs were purged with Purge_Dups v1.2.6 (https://github.com/dfguan/purge_dups), and potential contaminant sequences were identified and removed using FCS-GX v0.5.4 (https://github.com/ncbi/fcs-gx). Hi-C data were processed with Pairtools v1.1.0 (Open2C et al., 2024). Contigs were preliminarily anchored and ordered using YaHS v1.2.2 (Zhou et al., 2023) and Juicer v1.6 (Durand et al., 2016). Misassemblies and orientation errors were manually corrected and validated in Juicebox v1.11.08 (https://github.com/aidenlab/Juicebox). Chromosomal interaction heatmaps were generated with HapHiC v1.0.6 (Zeng et al., 2024). Telomeric and centromeric regions were predicted using quarTeT v1.2.2 (Lin et al., 2023). The genome assembly was polished with NextPolish2 v0.2.0 (https://github.com/Nextomics/NextPolish2), and completeness was assessed with BUSCO v5.4.7 (Manni et al., 2021) against the eukaryota_odb10 database.

### Genome annotation

Genomic repetitive sequences were annotated using HiTE v3.2.0 (Hu et al., 2024). Protein-coding genes were predicted through a combination of transcriptome-based, homology-based, and ab initio prediction approaches. Transcriptomic data were obtained from previous work conducted by our research group (PRJNA1049523). The gene sets derived from these three methods were integrated using EGAPx v0.3.2-alpha (https://github.com/ncbi/egapx). Transposable elements and misannotated sequences were filtered out to produce the final gene annotation. Genome annotation completeness was assessed with BUSCO v5.4.7 (Manni et al., 2021), and the LAI (LTR Assembly Index) was evaluated using LAI vbeta 3.2 (Ou et al., 2018). Protein sequences were aligned against the UniProt, NCBI non-redundant (Nr), KOG, Pfam, InterPro, and KEGG databases to infer putative biological functions and metabolic pathways based on sequence similarity. Whole-genome self-collinearity blocks were identified with JCVI v1.5.1 (Tang et al., 2024) and visualized using Circos v0.69-9 (Krzywinski et al., 2009). Non-coding RNAs were annotated as follows: miRNAs and snRNAs were identified with Infernal v1.1.5 (Nawrocki and Eddy, 2013) by searching against the Rfam v15.0 database; tRNAs were predicted using tRNAscan-SE v2.0.12 (Chan et al., 2021); and rRNAs were predicted with Barrnap v0.5.1 (https://github.com/tseemann/barrnap). Protein domains were annotated using PfamScan v1.6 (http://ftp.ebi.ac.uk/pub/databases/Pfam/Tools). Gene Ontology (GO) terms were assigned via InterProScan (Blum et al., 2021). Putative metabolic pathways were inferred by aligning genes to the KEGG database (release 84.0) using BLAST with an E-value cutoff of 1×10⁻⁵.

Raw RNA reads were processed using Trimmomatic v0.39 (Bolger et al., 2014) to generate clean reads through filtering and trimming. Subsequently, expression levels were quantified in transcripts per million (TPM) using Salmon v1.10.1. TPM values were used to visualize gene expression patterns across tissues, and heatmaps were generated using the Heatmap Illustrator function in TBtools-II (Chen et al., 2023a). The transcriptome data used in this study are available under the following accession numbers: PRJNA1049523 for *D. asperoides*, PRJCA019252 for *T. glandulifera*, and GSE81949 for *L. japonica*.

### Comparative genomics

Orthologous groups among *D. asperoides* and 15 other sequenced plant species— including *T. glandulifera* (Zhang et al., 2024), *Arabidopsis thaliana* (Swarbreck et al., 2007), *Oryza sativa* (PRJCA011659), *Solanum tuberosum* (GCA_020169585.1), *L. japonica* (Pu et al., 2020), *Tripterygium wilfordii* (GCA_013401445.1), *Daucus carota* (Iorizzo et al., 2016), *Zea mays* (GCA_000005005.6), *Coffea canephora* (GCA_900059795.1), *Scutellaria baicalensis* (PRJCA016803), *Lactuca sativa* (GCA_900198505.1), *Populus trichocarpa(v4.1)* (Tuskan et al., 2006), *Vitis vinifera* (Shi et al., 2023), *Solanum lycopersicum* (GCA_000181095.1), and the outgroup *Amborella trichopoda* (Amborella Genome Project, 2013)—were identified using OrthoFinder v2.3.11 (Emms and Kelly, 2019). Single-copy orthologous genes were selected to reconstruct a maximum likelihood phylogenetic tree with d RAxML-NG v1.2.2 (Kozlov et al., 2019). Divergence times were estimated using the MCMCTree program in PAML v4.10.7 (Yang, 2007), incorporating fossil calibration points. Gene family expansion and contraction analyses were performed with CAFE5 v1.1 (Mendes et al., 2021). Functional enrichment analysis of expanded gene families was conducted based on Gene Ontology (GO) and KEGG pathways using clusterProfiler v4.6.2 (Wu et al., 2021) in R v4.4.1.

### Analysis of whole-genome duplication

Potential homologous gene pairs within the *D. asperoides* genome and between *D. asperoides* and the genomes of *Triplostegia glandulifera*, *Lonicera japonica*, and Vitis vinifera were identified using BLASTP+ v2.15.0. The BLAST results were subsequently used as input for WGDI v0.6.5 (Sun et al., 2022) or JCVI v1.4.16 (Tang et al., 2024) to generate dot plots. The synonymous substitution rates (*Ks*) of paralogous and orthologous genes were calibrated with ksrates v1.1.4 (Sensalari et al., 2022) to accurately identify whole-genome duplication events and divergence times among the species examined. Synteny analysis between *D. asperoides* and *T. glandulifera*, *L. japonica*, and *V. vinifera* was performed using JCVI v1.4.16 (Tang et al., 2024).

Genome-wide duplication events in *D. asperoides*, *T. glandulifera*, and *L. japonica* were classified into five categories using DupGen_Finder v1.12 (Qiao et al., 2019) under default parameters. The non-synonymous (*Ka*) and synonymous (*Ks*) substitution rates, as well as *Ka/Ks* ratios, were calculated for duplicated gene pairs using the YN model implemented in *KaKs*_Calculator2 (Wang et al., 2010). Functional enrichment analysis for each duplication type was performed based on Gene Ontology (GO) and KEGG pathways using clusterProfiler v4.6.2 (Wu et al., 2021) in R v4.4.1.

### Identification of *OSC*, *CYP*, and *UGT* family members

Genes encoding oxidosqualene cyclases (OSCs), cytochrome P450s (CYPs), and UDP-glycosyltransferases (UGTs) were identified from the protein-coding genes of *D. asperoides*, *T. glandulifera*, and *L. japonica* based on Pfam annotations: those containing domains PF13249 or PF13243 were classified as OSCs, PF00067 as CYPs, and PF00201 as UGTs. A subset of the identified UGTs was officially named by the UGT Nomenclature Committee. Multiple sequence alignment was performed using MAFFT v7.520 (Rozewicki et al., 2019), followed by trimming with trimAl v1.4.1 (Capella-Gutiérrez et al., 2009). A maximum-likelihood phylogenetic tree was reconstructed with IQ-TREE v2.2.2.7 (Nguyen et al., 2015) under 1000 bootstrap replicates to analyze the phylogenetic relationships among OSCs, CYPs, and UGTs from *D. asperoides*, *T. glandulifera*, *L. japonica*, and other functionally characterized homologs.

### Gene cloning of candidate *OSC*, *CYP*, and *UGT* genes

Candidate genes encoding OSC, CYP, and UGT were cloned into the pYES2, y33, pET-28a, and pEAQ vectors, respectively, using primer pairs listed in Supplementary Table 18. The constructed plasmids were subsequently transformed into *E. coli* DH5α for Sanger sequencing. After verification, the validated plasmids were extracted and stored at −20 °C for further use.

### Functional characterization of candidate *OSC* and *CYP* genes

The recombinant plasmid pYES2-OSC was transformed into the lanosterol-deficient yeast strain GIL77. Transformants were cultured in 50 mL of uracil-deficient synthetic complete (SC-Ura) medium supplemented with 2% glucose, ergocalciferol (20 μg/mL), hemin (13 μg/mL), and Tween-80 (5 mg/mL) at 30 °C for 48 hours. The culture was then induced by replacing glucose with 2% galactose and incubated for an additional 48 hours (Chen et al., 2023b). Cells were harvested by high-speed centrifugation and resuspended in a solution of 20% KOH in 50% ethanol, followed by heating at 100 °C for 10 min. The mixture was extracted three times with petroleum ether. After evaporation of the solvent, the residue was derivatized with 100 μL trimethylsilyl cyanide (TMSCN) at 70 °C for 30 min. The derivatized samples were analyzed using an Agilent 7890A gas chromatograph equipped with an HP-5 MS quartz capillary column (30 m × 0.25 mm i.d., 0.25 μm film thickness). The injector, ion source, and transfer line temperatures were set to 250 °C. The oven temperature program was as follows: hold at 80 °C for 2 min; increase to 290 °C at 20 °C/min; hold at 290 °C for 30 min. Helium was used as the carrier gas at a flow rate of 1.2 mL/min. Samples were injected in splitless mode with an injection volume of 3 μL. Mass spectrometry was performed using an Agilent 6540 Accurate-Mass Q-TOF system.

The recombinant plasmid y33-CYP was transformed into the *β*-amyrin-producing Saccharomyces cerevisiae strain BA-BY4742 or the oleanolic acid-producing strain OA-BY4742. Positive transformants were inoculated into synthetic complete (SC) medium lacking histidine, leucine, and uracil, supplemented with 2% glucose, and cultured at 30 °C with shaking at 200 rpm for 96 hours. Cells were harvested by high-speed centrifugation and resuspended in 10 mL of 50% aqueous methanol, followed by ultrasonication for 30 minutes. The suspension was extracted three times with ethyl acetate. The ethyl acetate phase was evaporated to dryness, and the residue was reconstituted in 1 mL of methanol, filtered, and subjected to GC–MS or LC–MS analysis. GC–MS analysis was performed under the same conditions as those used for OSC functional characterization. LC–MS analysis was carried out using an Agilent 1290 UPLC system coupled with a 6540 Q-TOF mass spectrometer. Mass spectrometry was conducted in negative ion mode with the following settings: capillary voltage, 3500 V; fragmentor voltage, 135 V; skimmer voltage, 60 V; octopole RF voltage, 750 V; scan range, 50–1700 m/z; and scanning mode, SRM. Chromatographic separation was achieved on an Agilent Extend-C18 column (250 mm × 4.6 mm, 5 μm) maintained at 30 °C. The mobile phase consisted of methanol with 0.01% (v/v) formic acid (A) and acetonitrile (B) with the following gradient program: 0–12 min, 85% to 33% A; 12–35 min, 33% to 15% A; 35–40 min, 15% to 0% A. The total run time was 40 min, the injection volume was 10 μL, and the flow rate was 0.8 mL/min.

### Functional characterization of candidate *UGT* genes

The recombinant plasmid pET-28a-UGT was transformed into *E. coli* BL21(DE3) cells. Positive colonies were selected and cultured in 500 mL of LB medium containing kanamycin (50 μg/mL) at 37 °C until the OD_600_ reached 0.6–0.8. The temperature was then reduced to 16 °C, and protein expression was induced with 0.5 mM IPTG for 12 h. After induction, cells were harvested by centrifugation and resuspended in Tris-HCl buffer (pH 7.5, containing 200 mM NaCl and 50 mM Tris) for protection. The cells were lysed by ultrasonication, and the target protein was purified using His-Tag affinity chromatography. The purified protein was analyzed by SDS-PAGE, and its concentration was determined using a BCA protein assay kit. A 100 μL reaction system was prepared containing 1 mM UDP-sugar donor, 1 mM substrate, 2% (v/v) Tween-80, and 50 μg of enzyme, followed by incubation at 40 °C for 12 h. The reaction was terminated by adding 100 μL methanol, and the mixture was filtered for HPLC–MS/MS analysis. For substrate-screening assays, apparent conversion rates were estimated from substrate depletion relative to the heat-inactivated enzyme control. The purified enzyme was heat-inactivated by boiling and used as the negative control under otherwise identical reaction conditions. The apparent conversion rate was calculated as follows: apparent conversion rate (%) = (Acontrol − Areaction) / Acontrol × 100, where Acontrol and Areaction represent the integrated LC–MS peak areas of the substrate in the heat-inactivated control and active-enzyme reaction, respectively. Analyses were performed on an Agilent 1290 UPLC system coupled with a 6540 Q-TOF mass spectrometer using an Agilent Extend-C18 column (250 mm × 4.6 mm, 5 μm). The mobile phase consisted of water containing 0.01% formic acid (A) and acetonitrile (B). A gradient elution program was applied as follows: 0–3 min, 85% to 60% A; 3–7 min, 60% to 35% A; 7–9.5 min, 35% to 12% A; 9.5–25 min, 12% A. The total run time was 25 min, the injection volume was 10 μL, and the flow rate was 0.5 mL/min.

### Transient expression of candidate genes in *N. benthamiana*

The candidate genes were cloned into the pEAQ-HT vector and transformed into Agrobacterium tumefaciens strain (GV3101). A. tumefaciens strains harboring recombinant plasmids containing the candidate genes were inoculated into 15 mL of LB liquid medium supplemented with 25 μg/mL rifampicin and 50 μg/mL kanamycin and cultured at 28°C for 24 h. The bacterial cells were then collected by centrifugation at 5,000 rpm for 10 min and resuspended in 5 mL of induction buffer containing 10 mM 2-(N-morpholino)ethanesulfonic acid (MES; pH 5.6), 10 mM MgCl₂, and 100 μM acetosyringone. The bacterial suspension was diluted with induction buffer to an optical density at 600 nm (OD₆₀₀) of 0.2 and incubated at room temperature for 2 h. Subsequently, the suspension was infiltrated into the abaxial side of leaves from 4-week-old Nicotiana benthamiana plants using a 1-mL needleless syringe. After 4 d of incubation, the infiltrated leaves were harvested for subsequent metabolite analysis.

### Protein structure modeling and docking

Protein sequence alignment was performed using Jalview. Protein three-dimensional structure prediction and molecular docking were conducted with Boltz-1 (https://github.com/jwohlwend/boltz). Prediction of catalytic pockets on the target protein was performed using the Proteins Plus web service (https://proteins.plus/pages/about). Resulting docking models were visualized using PyMOL. Site-directed mutagenesis PCR targeting selected residues was carried out with primers listed in the Supplementary Tables 19 and 20. Protein purification and enzyme activity assays were performed as described in the previous section.

### Classical Molecular Dynamics (MD) Simulations

The protein and small molecules were described using the Amber99sb-ildn and GAFF force fields, respectively. The complex system was solvated with the TIP3P water model in a cubic box, and Na⁺/Cl⁻ ions were added to neutralize the system. After energy minimization, NVT equilibration, and NPT equilibration, all positional restraints were removed, followed by a 100 ns unrestrained production simulation under the NPT ensemble. Bond lengths involving hydrogen atoms were constrained using the LINCS algorithm, long-range electrostatic interactions were treated using the PME method, and a time step of 2 fs was applied. The resulting trajectories were used for subsequent analyses of system stability and molecular interactions.

### Phylogenetic and selection pressure analyses of UGT76 and UGT85 genes

UGT family members were retrieved from the Pfam annotations of the *D. asperoides*, *T. glandulifera*, and *L. japonica* genomes. Based on phylogenetic classification, members of the UGT76 and UGT85 subfamilies were selected for subsequent selection pressure analysis. Protein sequences were aligned using MAFFT v7.520 and trimmed using trimAl v1.4.1 to remove gaps and poorly aligned regions. A maximum-likelihood phylogenetic tree was constructed using IQ-TREE v2.2.2.7 with 1,000 bootstrap replicates. Codon-based alignments were generated from the protein alignments and corresponding coding sequences (CDSs) using PAL2NAL. The resulting codon alignments and phylogenetic tree were used for selection pressure analysis using CODEML implemented in the PAML package. The one-ratio model (M0) and two-ratio branch model (M2) were used to estimate branch-specific *Ka/Ks* ratios. Selected internal branches were sequentially designated as foreground branches to estimate branch-specific *Ka/Ks* values, which were subsequently mapped onto the phylogenetic tree. Branches with *Ka/Ks* > 1, *Ka/Ks* = 1, and *Ka/Ks* < 1 were considered to show signatures consistent with positive selection, neutral evolution, and purifying selection, respectively.

### Ancestral sequence reconstruction

For ancestral sequence reconstruction of the UGT85CA-related lineage, the protein sequences of TgUGT85CA3-1, TgUGT85CA3-2, DaUGT85CA8, DaUGT85J2 (CA7), LjUGT85CA2, LjUGT85CA4, LjUGT85CA5, LjUGT85CA6, Das06G003119.t1, TgChr08G27397.1, Lj6C642T14.1, Lj5A236T61.1, Lj9C352G2.1, Das03G001233.t1 (DaUGT85A245), and Das07G002084.t1, together with their corresponding CDS sequences, were collected. Protein sequences were aligned using MAFFT and trimmed using trimAl. A maximum-likelihood phylogenetic tree was constructed using IQ-TREE with 1,000 bootstrap replicates. Codon-based alignments were generated from the protein alignments and corresponding CDS sequences using PAL2NAL. Ancestral sequences at selected internal nodes were subsequently reconstructed using CODEML implemented in PAML. The reconstructed ancestral sequences were annotated at the corresponding internal nodes of the phylogenetic tree.

### Agrobacterium-mediated transient expression in *D. asperoides* leaves

Agrobacterium tumefaciens GV3101 strains carrying pEAQ-*β*AS2, pEAQ-DaCYP714E29, or pEAQ-DaUGT94BA4 were cultured on LB agar medium for 2 days. Before infiltration, the cells were resuspended in MES buffer and adjusted to an OD₆₀₀ of 0.6. For transient expression, one half of each selected leaf from 2-month-old Dipsacus asperoides plants was infiltrated with Agrobacterium carrying the target gene, while the other half was infiltrated with Agrobacterium carrying the empty vector as the control. Infiltration was performed using a needleless 1 mL syringe. After infiltration, the plants were kept in the dark for 12 h and then grown at 25 ℃ under a 16 h light/8 h dark photoperiod. After 5 days, the infiltrated leaf tissues were collected, frozen in liquid nitrogen, and used for qRT-PCR and LC-MS analysis. The primers used for qRT-PCR analysis are listed in Supplementary Table 21.

## DATA AVAILABILITY

The raw sequencing data generated in this study, including genome survey sequencing reads, PacBio HiFi sequencing reads, and Hi-C sequencing reads, have been deposited in the Genome Sequence Archive (GSA) of the National Genomics Data Center, China National Center for Bioinformation, under accession number CRA046475. The genome assembly sequences, genome annotation files, and associated datasets have been deposited in Figshare. The gene identifiers and corresponding source species information for the sequences analyzed in this study are provided in Supplementary Table 22. The amino acid sequences of ancestral enzymes reconstructed from internal nodes of the phylogenetic tree are provided in Supplementary Table 23.

## ACKNOWLEDGMENTS

The authors declare no conflict of interest.

## AUTHOR CONTRIBUTIONS

YL, SH, GZ and SY conceived the study. RY, CF, XZ and YZ contributed equally to this work. RY, CF, XZ, WH, ML, GX, JX, and YW performed the experiments. RY, YL, SH, GZ and SY the designed experiments. GX, GC and QL analyzed the data. RY, YL, SH, GZ and SY drafted the manuscript. YL, SH, GZ and SY reviewed and edited the manuscript. All authors contributed to the article and approved the submitted version.

