## Supplementary material for "Genome evolution and glycosyltransferase diversification shape asperosaponin VI biosynthesis in *Dipsacus asperoides*": Supplementary Information(1).docx

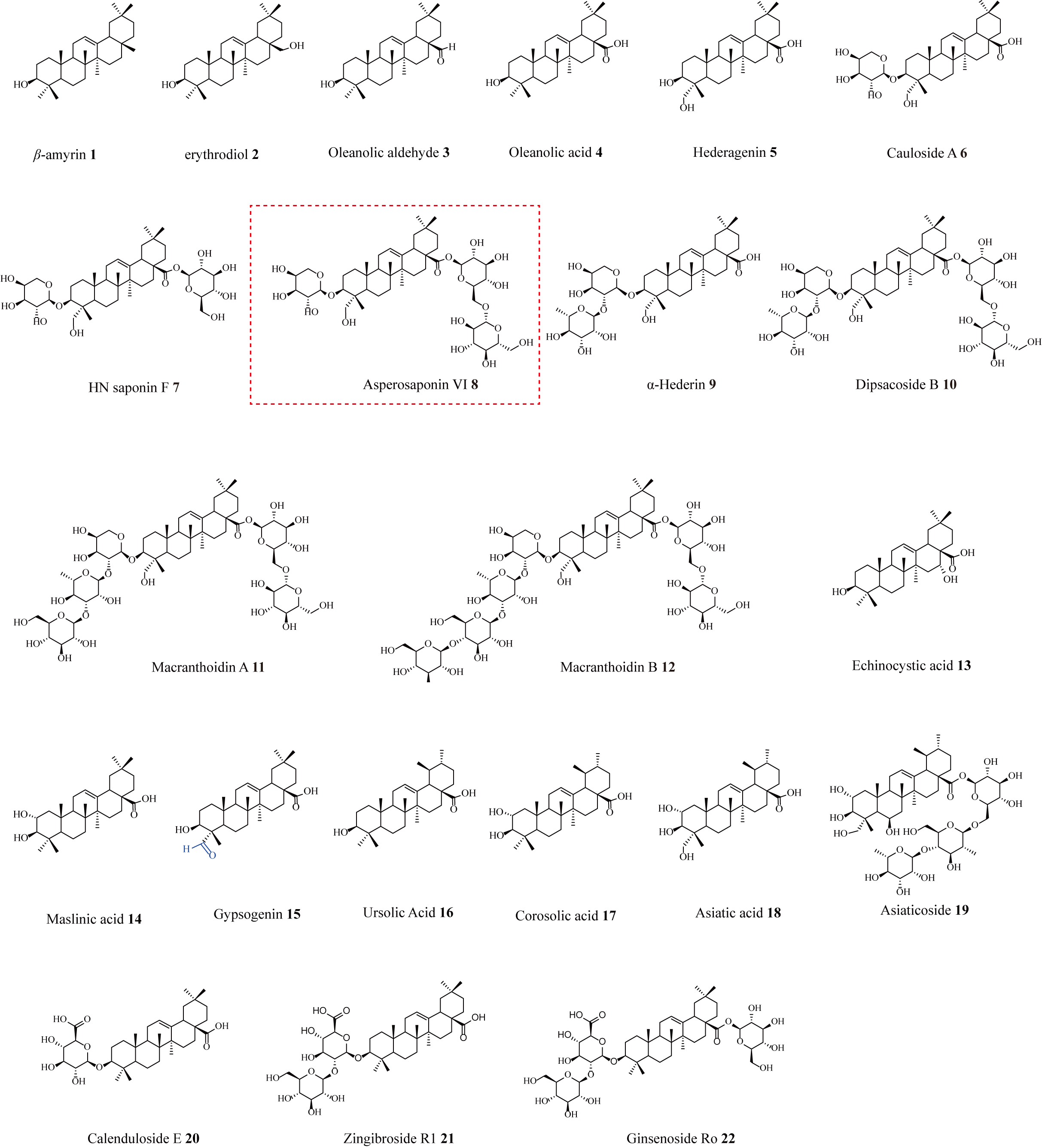

**Supplemental Figure 1.** **Structures of the analyzed triterpenoids**.

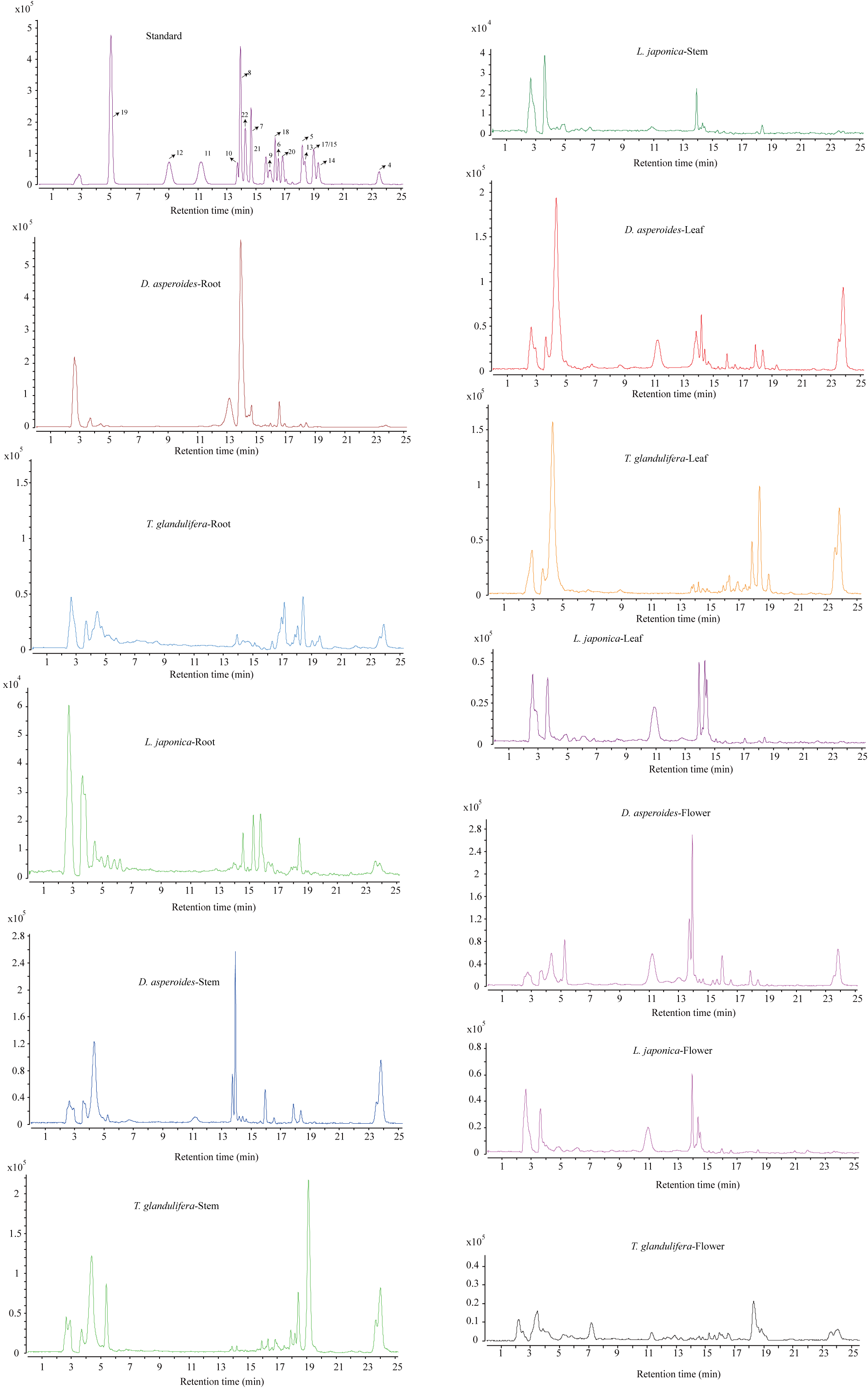
 **Supplemental Figure 2.** **LC-MS extracted ion chromatograms (EICs) of different tissues from three plant species (*D. asperoides*, *L. japonica* and *T. glandulifera*)**.

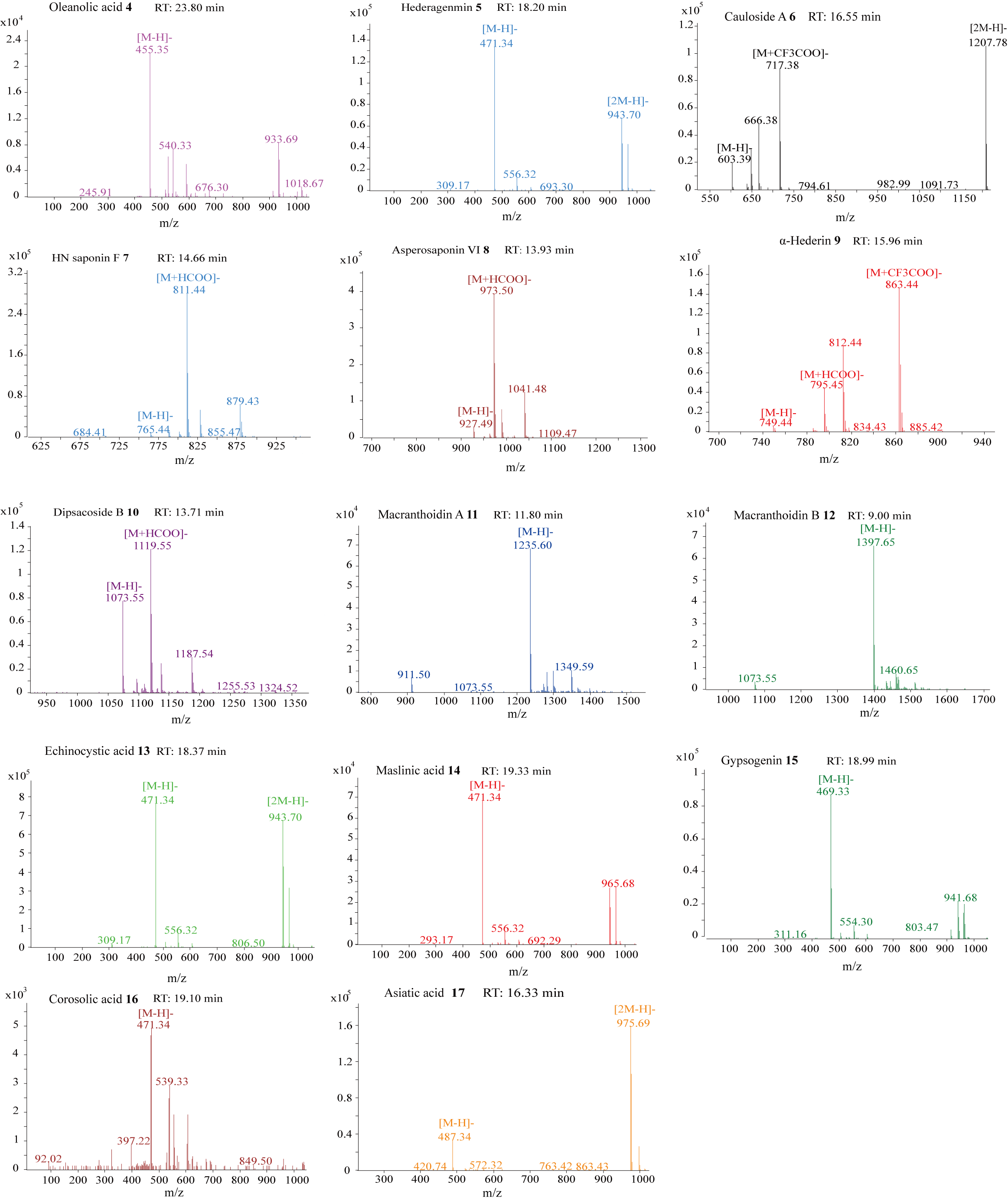

**Supplemental Figure 3.** **Mass profiles of triterpenoids identified in three plant (*D. asperoides*, *L. japonica* and *T. glandulifera*) species via LC-MS**.

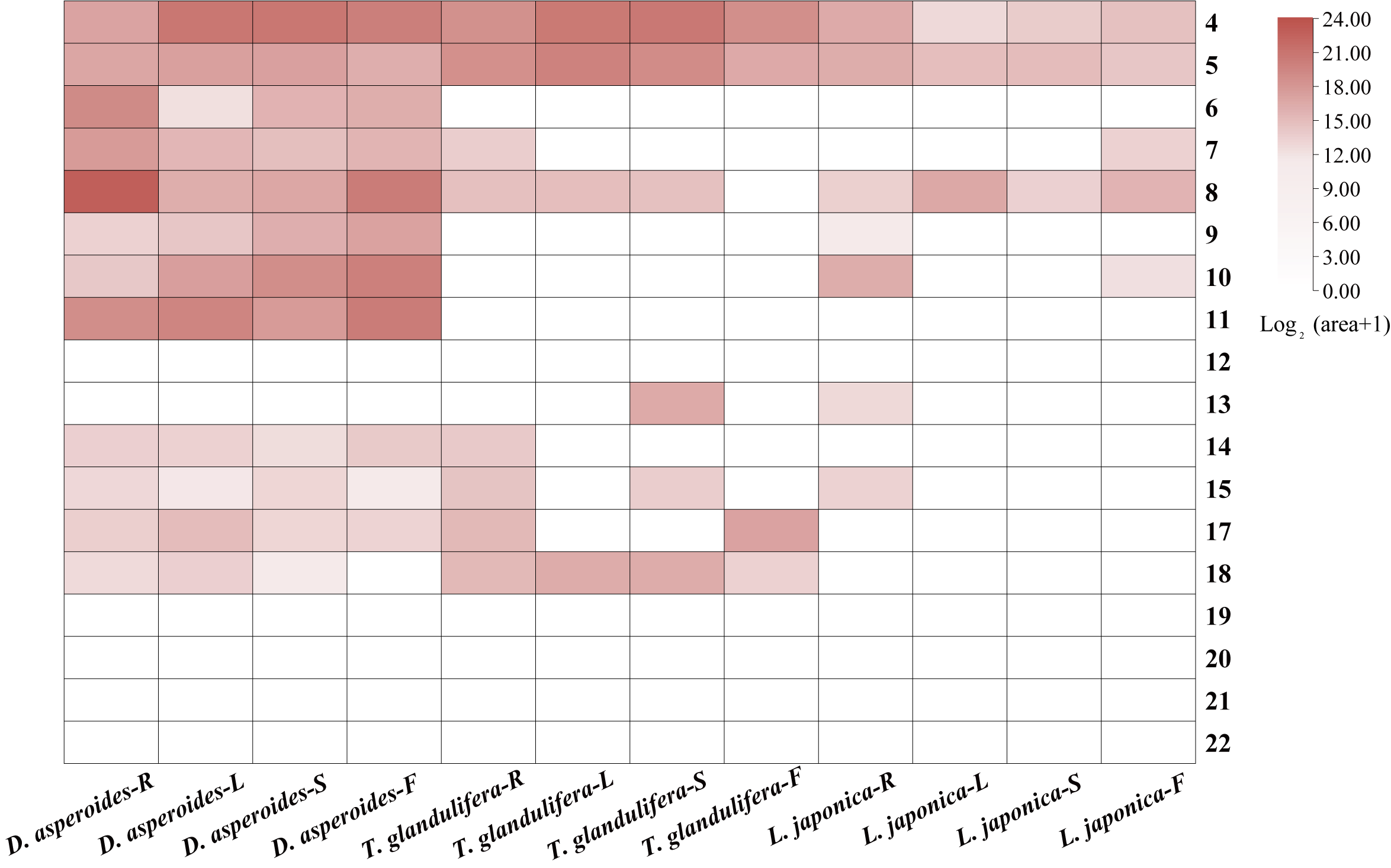

**Supplemental Figure 4.** **Metabolic profiling of 18 triterpenoids in three plant species.** The levels of hederagenin-based saponins, represented by values of log_2_ (peak area + 1), were plotted as a heatmap.

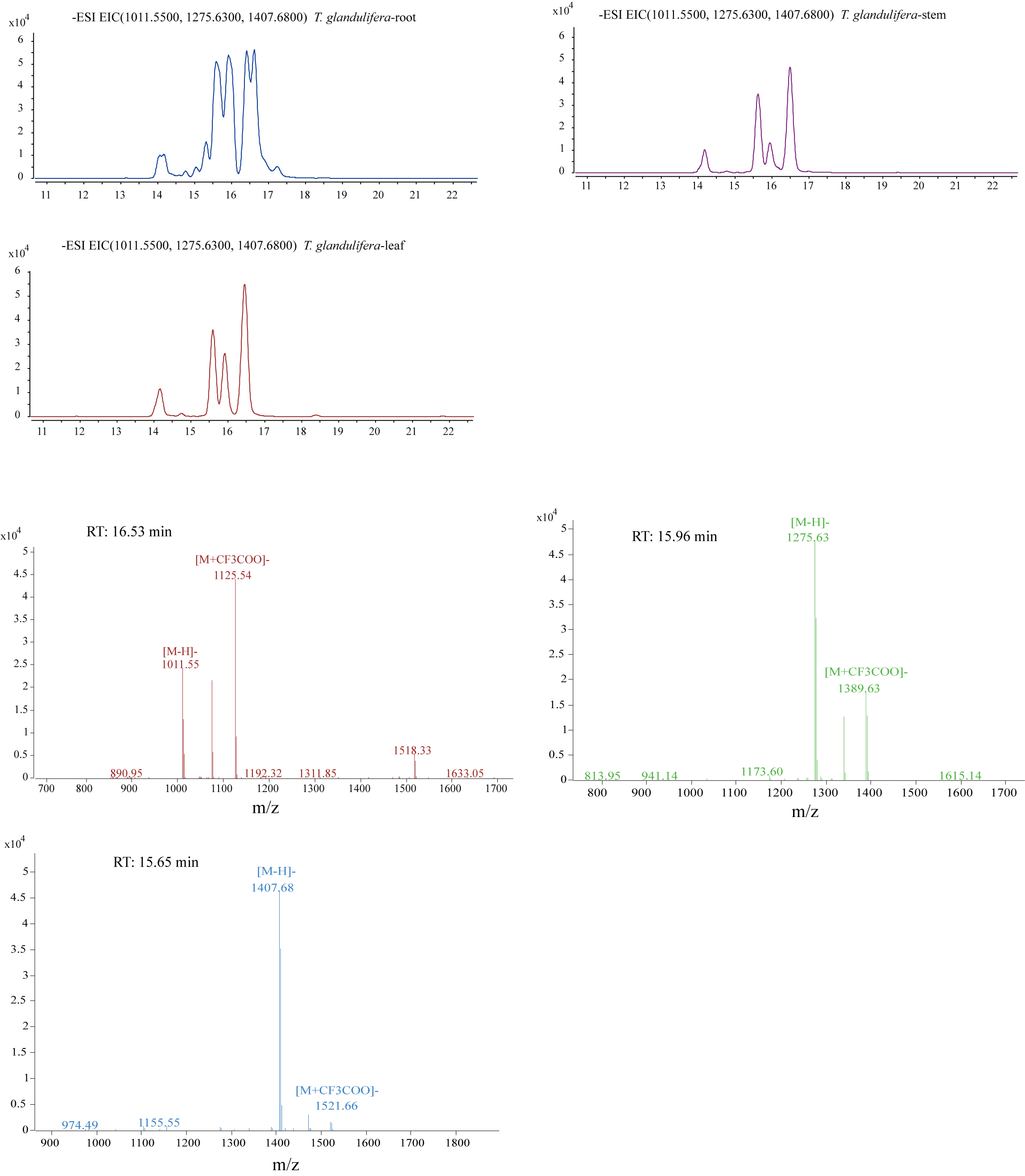

**Supplemental Figure 5.** **LC-MS EICs (1011.50，1275.63 and1407.68) of different tissues from *T. glandulifera***.

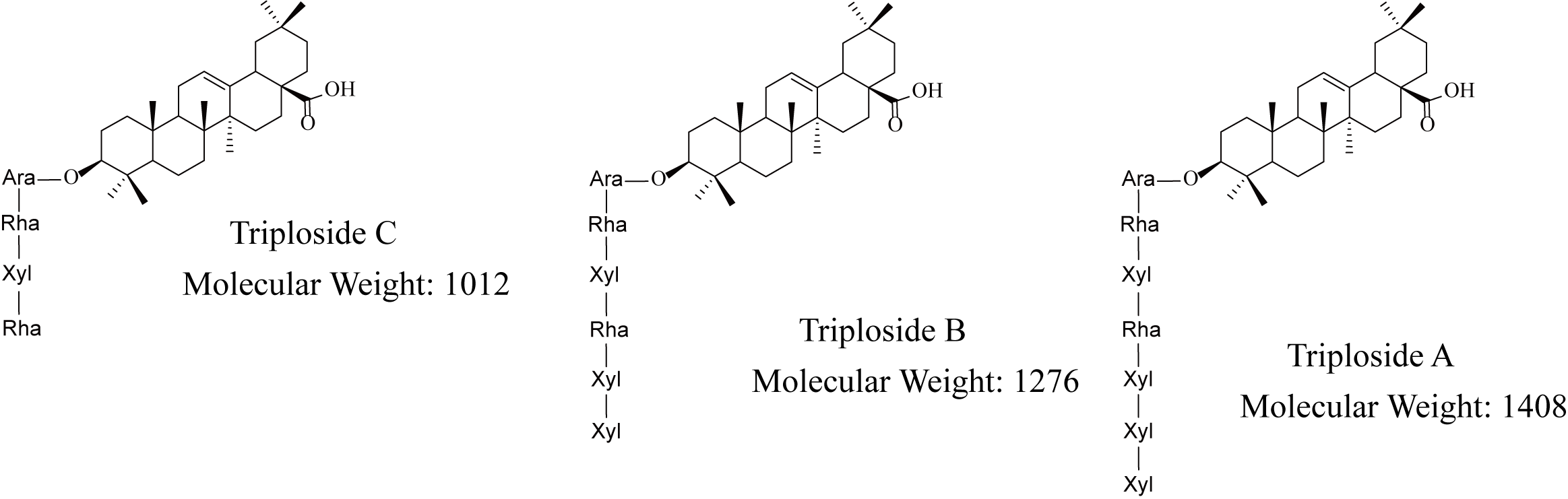

**Supplemental Figure 6. The structures of Triploside A, Triploside B, and Triploside C**.

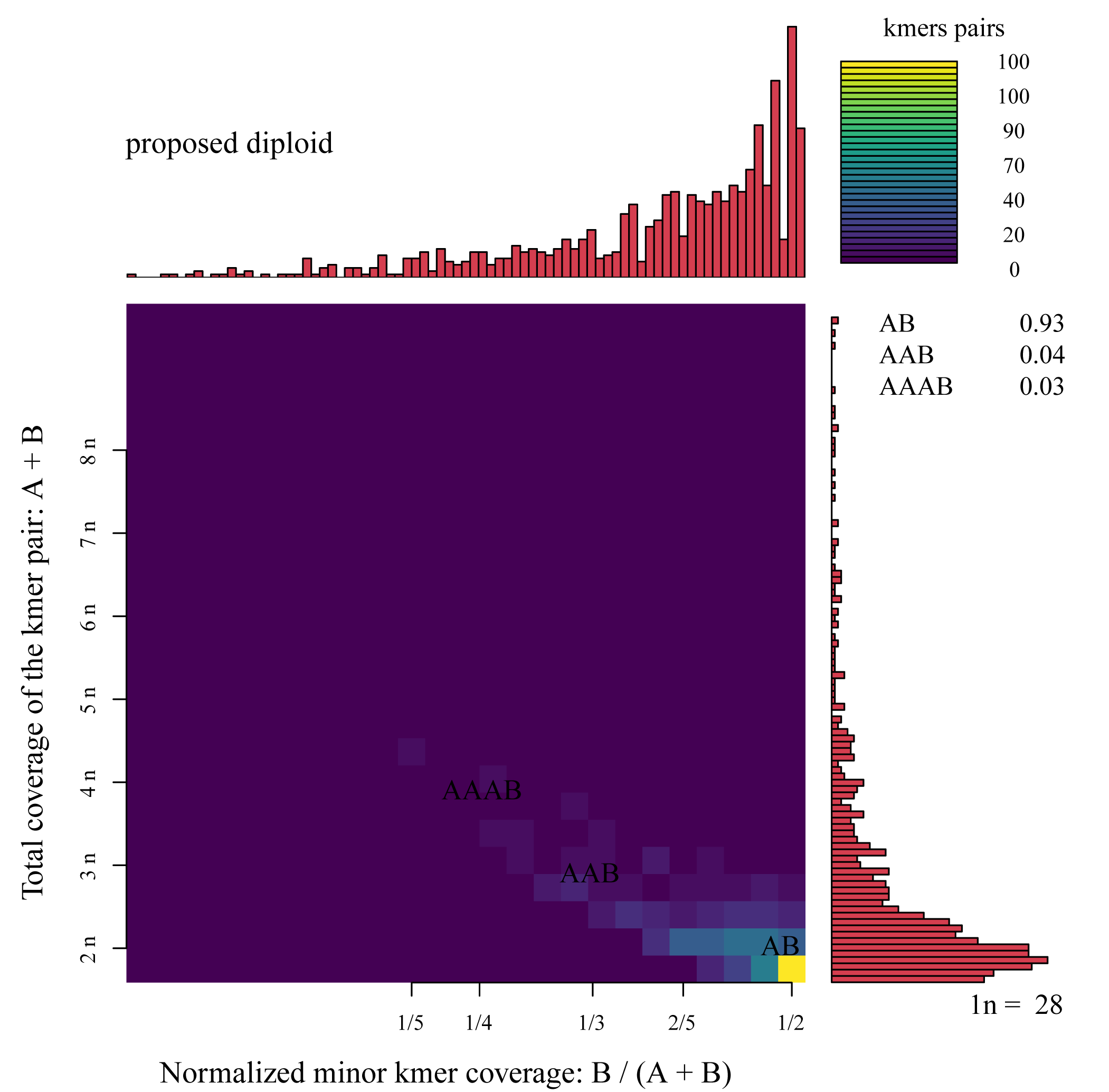

**Supplemental Figure 7.** **Ploidy analysis using Smudgeplot**.

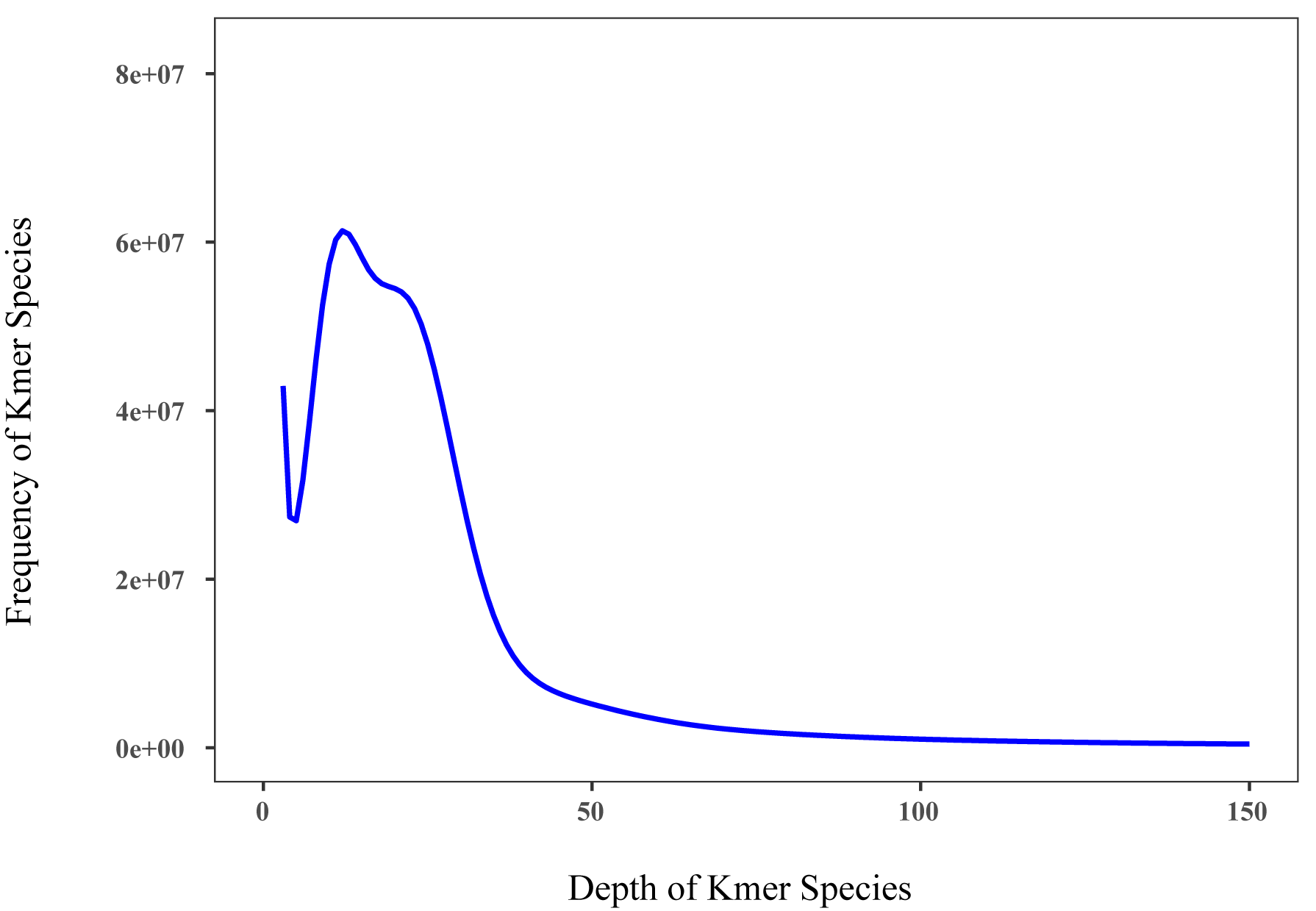

**Supplemental Figure 8.** **The *k-mer* (k=17) frequency-depth distribution map of** ***D. asperoides* genome shows the depth on the horizontal axis and the frequency of *k-mer* on the vertical axis**.

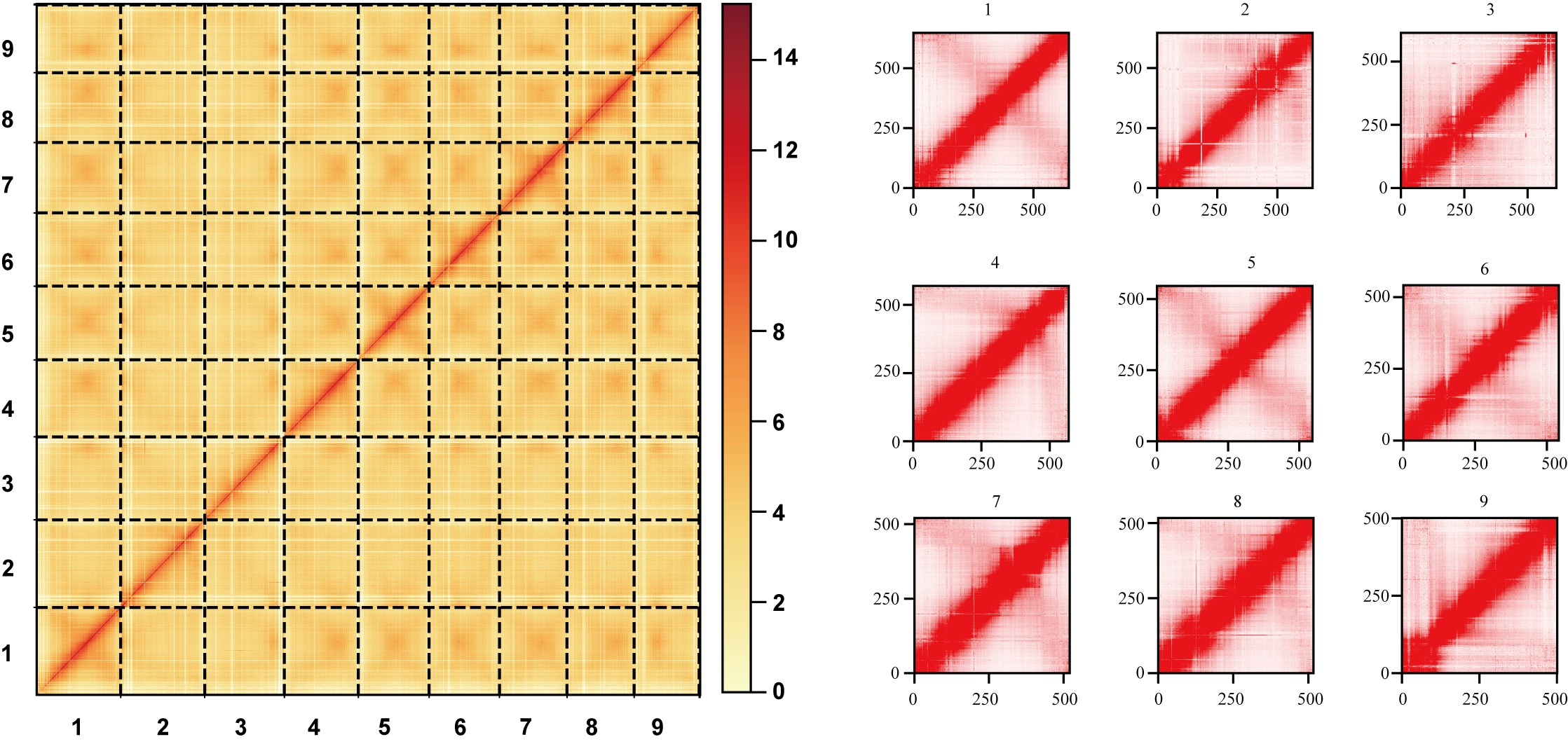

**Supplemental Figure 9. Hi-C interaction heatmap of 9 homoeologous chromosome groups in *D. asperoides****.*

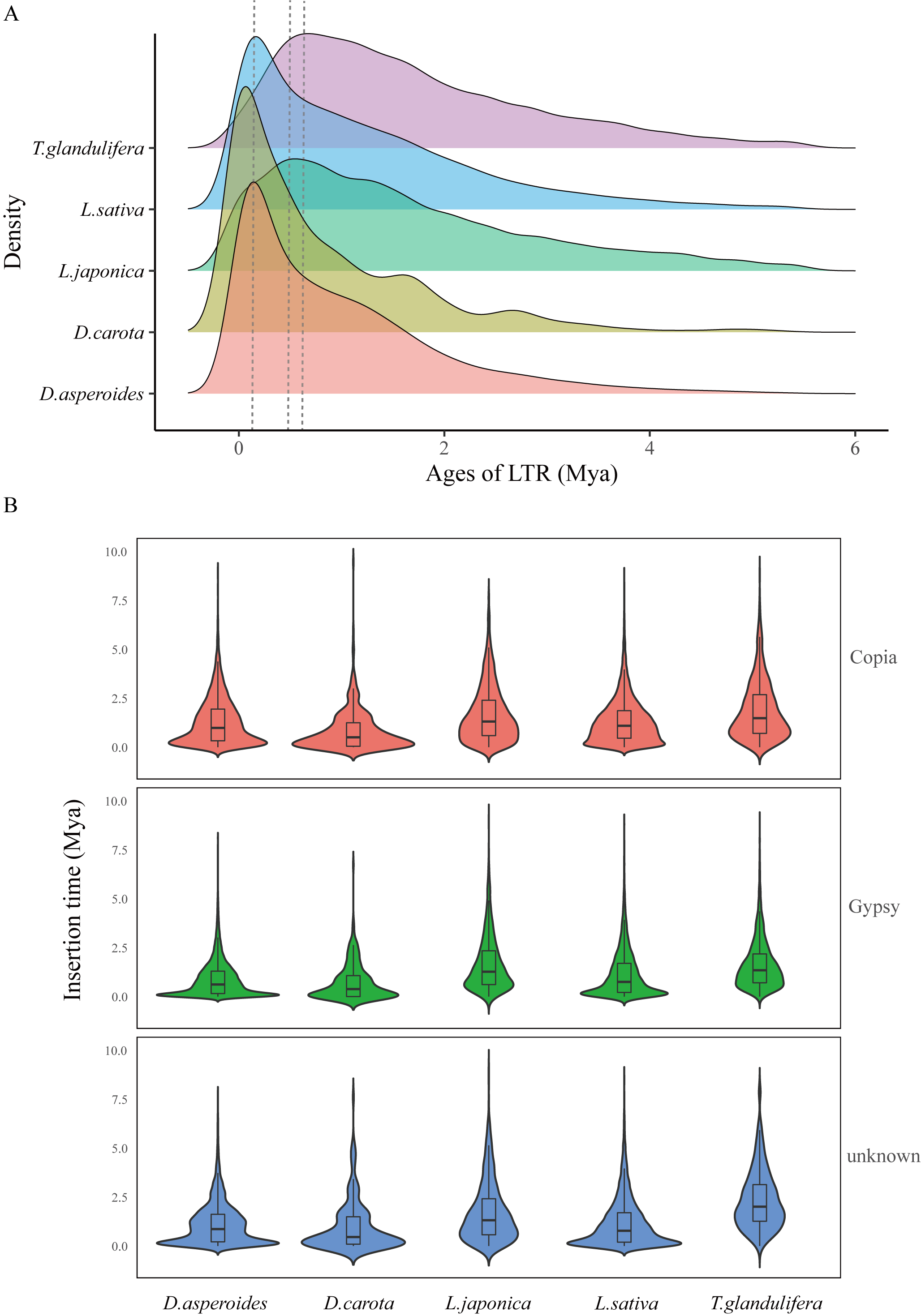

**Supplemental Figure 10. LTR insertion time comparison across** ***D. asperoides*, *D. carota*,** ***L. japonica*, *L. sativa*, and** ***T. glandulifera***. **(A)** Density distribution of estimated LTR insertion times (all types). **(B)** Estimated insertion times for Copia, Gypsy, and unknown.

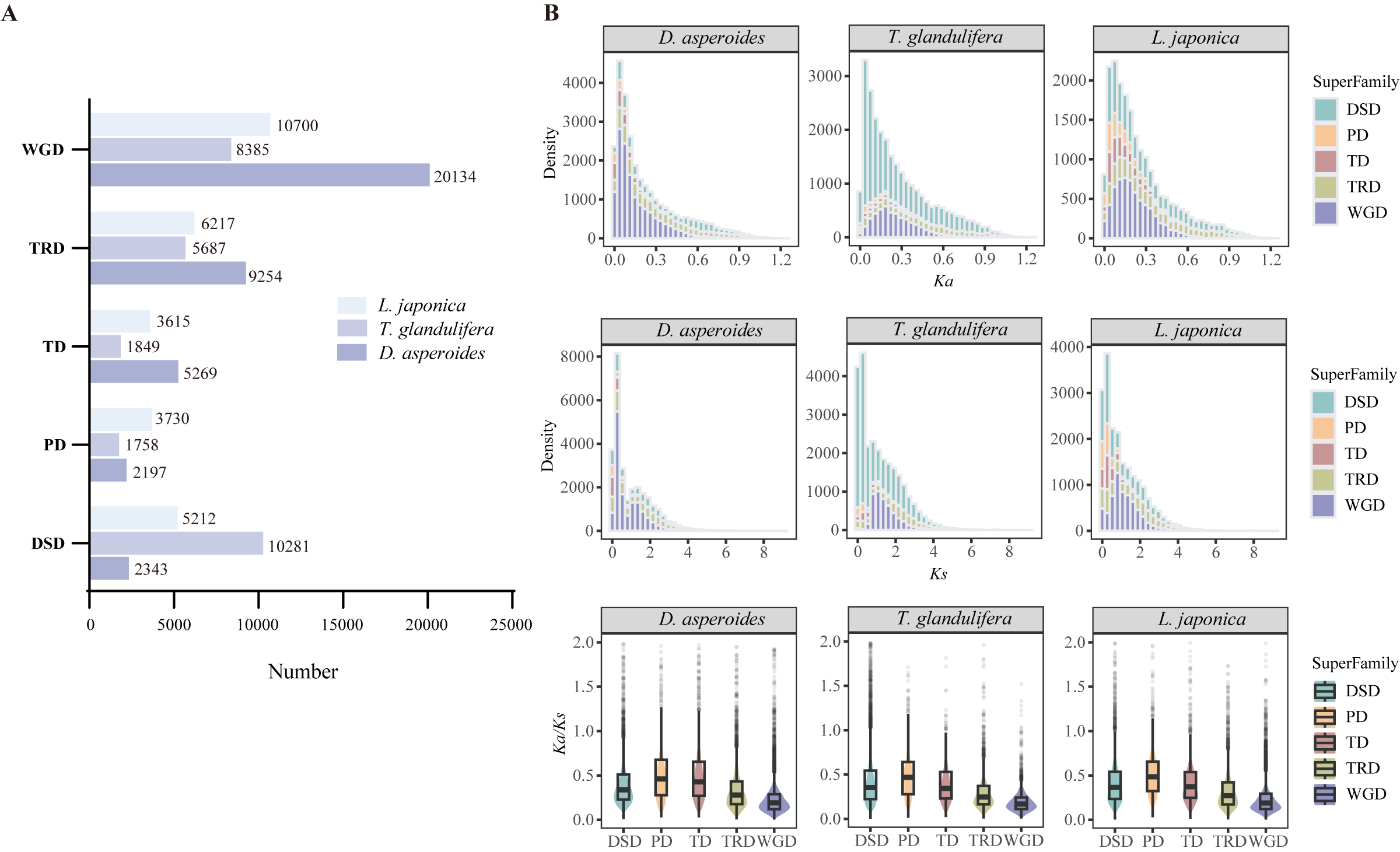

**Supplemental Figure 11**. **Identification and evolutionary analysis of gene duplication modes across three Caprifoliaceae species**. **(A)** Number of genes derived from five distinct duplication modes: whole-genome duplication (WGD), transposed duplication (TRD), tandem duplication (TD), proximal duplication (PD), and dispersed duplication (DSD). **(B)** Distribution of *Ka*, *Ks* values, and *Ka/Ks* ratios for duplicated gene pairs.

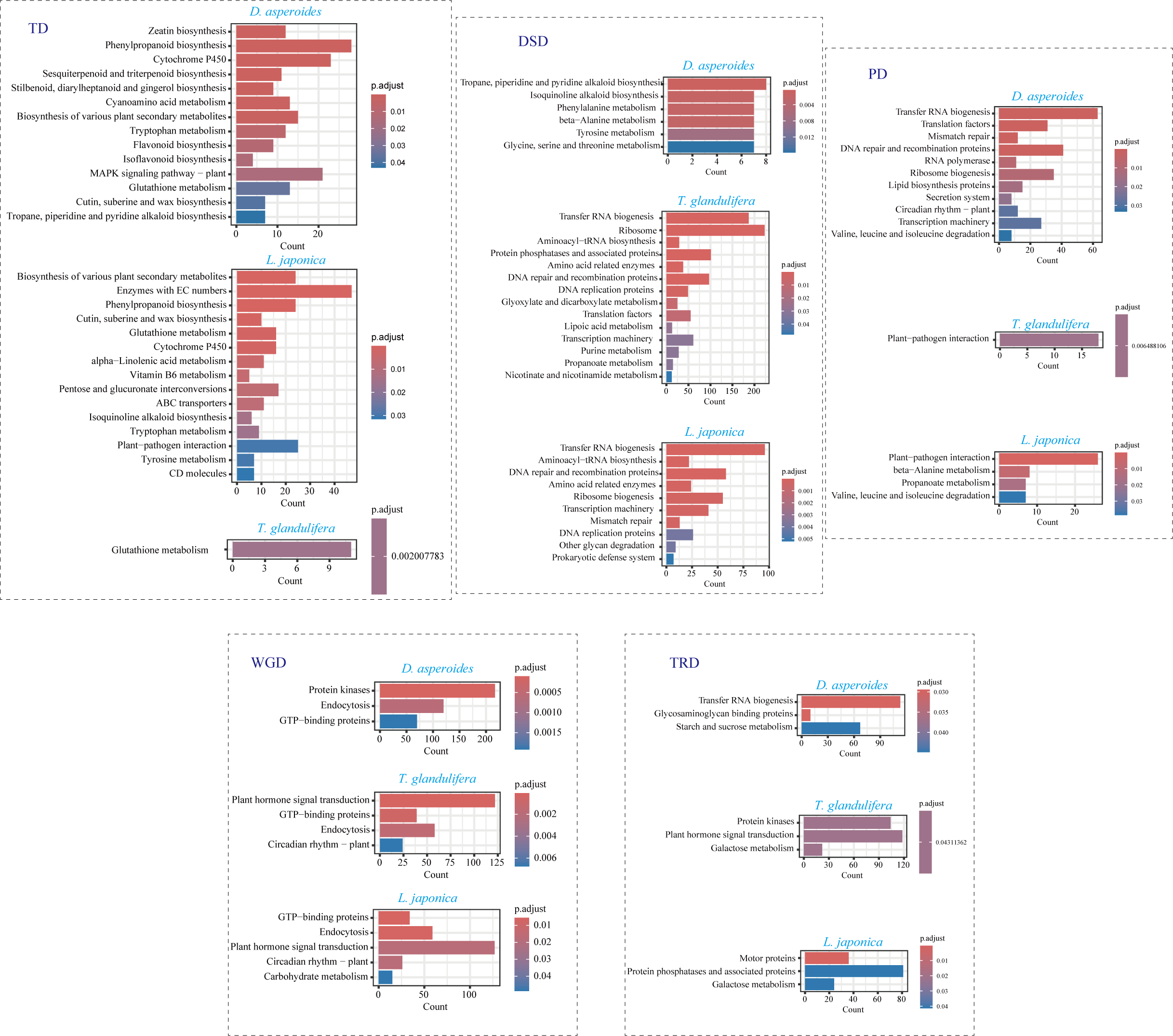

**Supplemental Figure 12.** **KEGG enrichment analyses of genes from different types of duplication**.

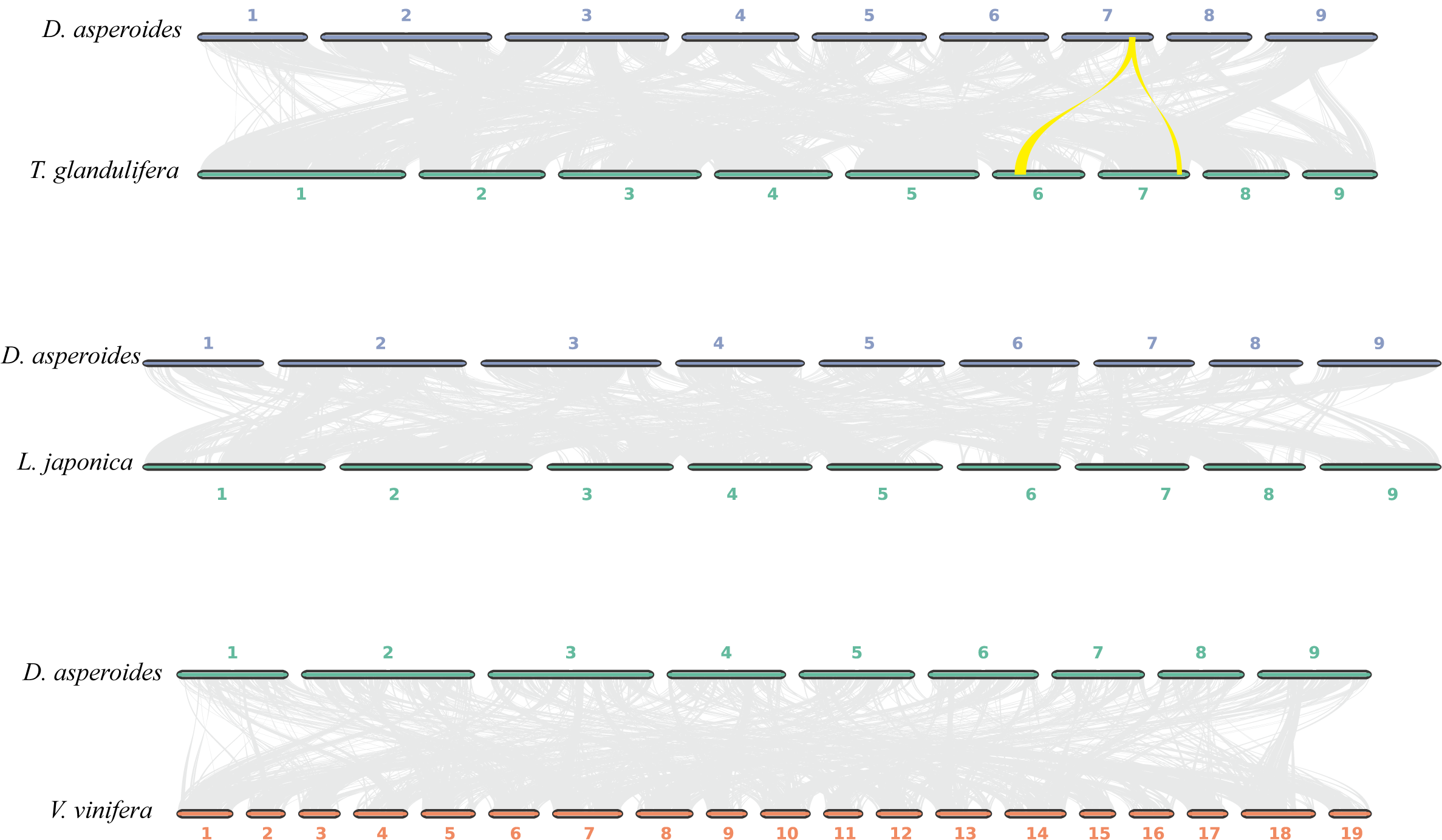

**Supplemental Figure 13. Comparative synteny analysis among *D. asperoides*, *T. glandulifera*, *L. japonica*, and *V. vinifera***.

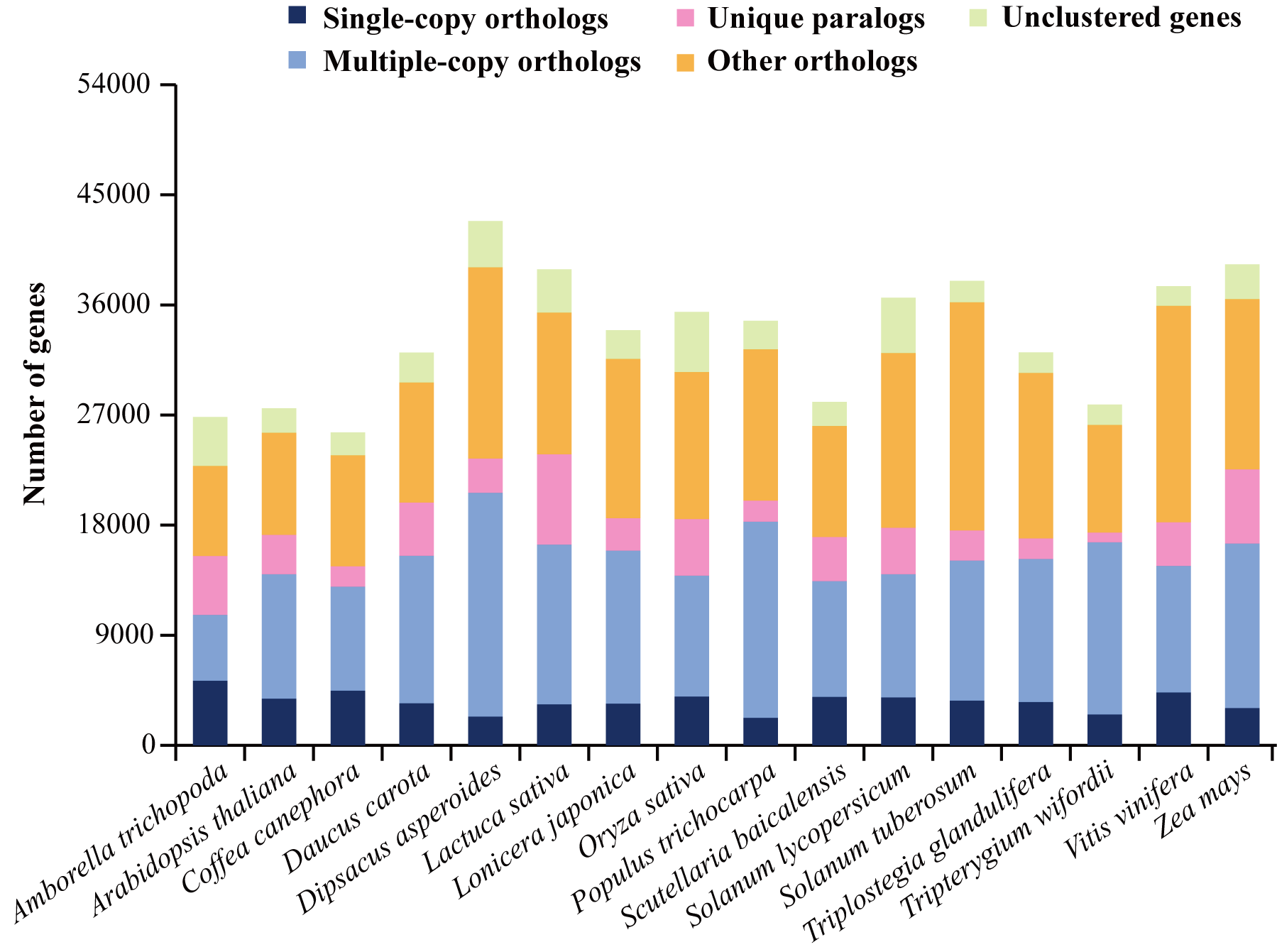

**Supplemental Figure 14. Homologous genes analysis results of 16 different species**.

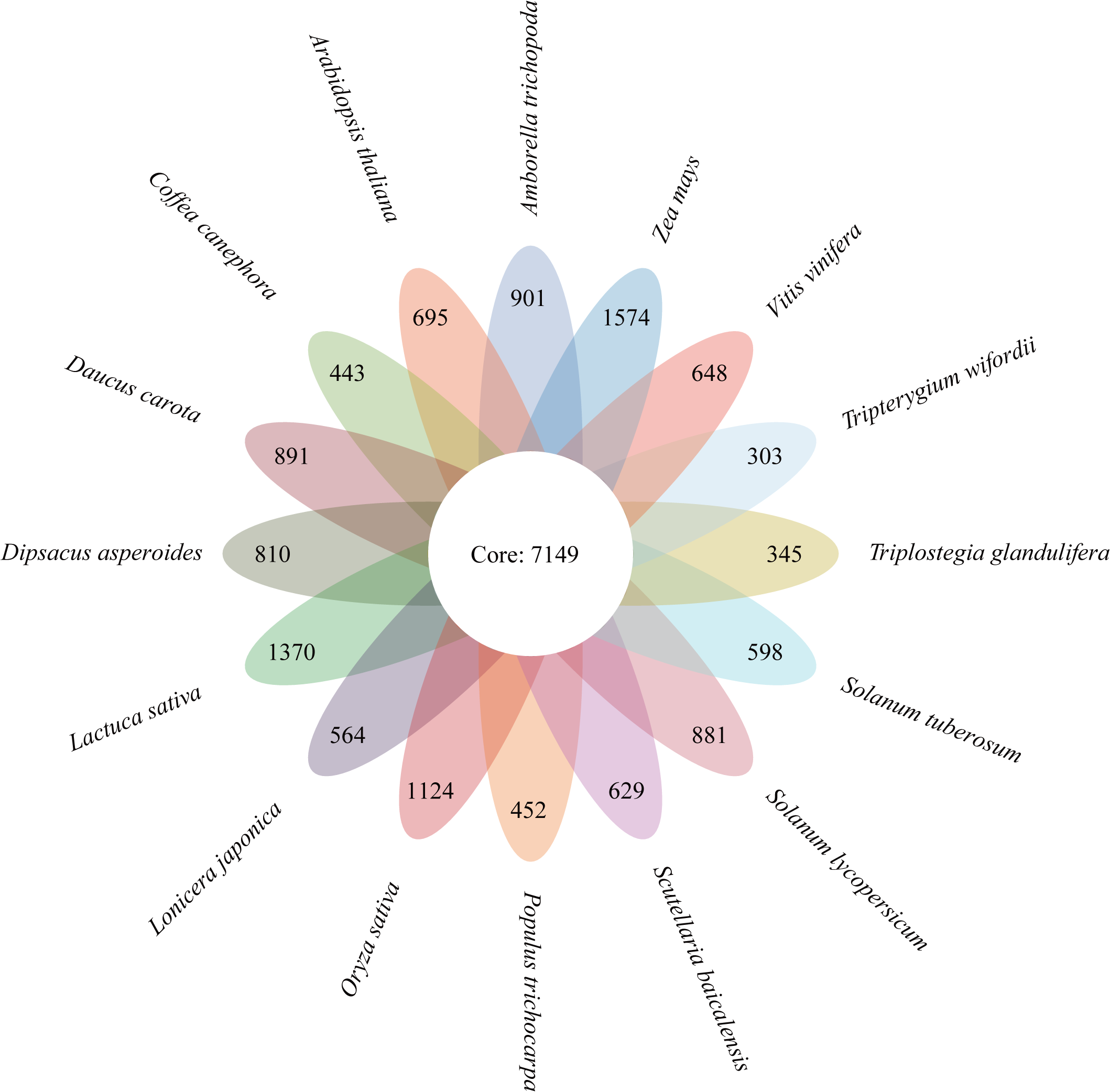

**Supplemental Figure 15. The flower plot shows the common orthogroups (center) of 16 species and their species-specific orthogroups**.

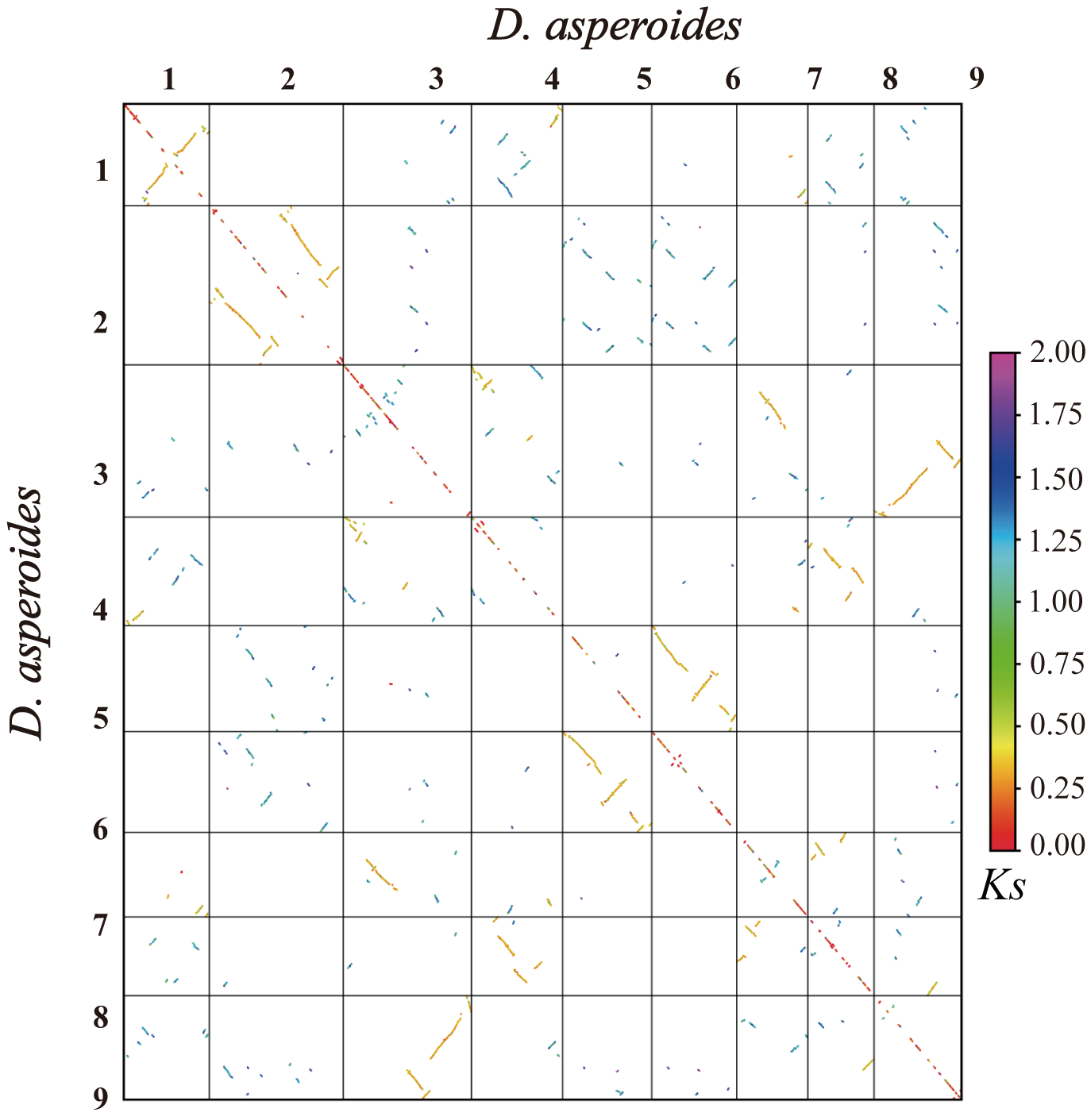

**Supplemental Figure 16. Syntenic blocks within the *D. asperoides* genome**.

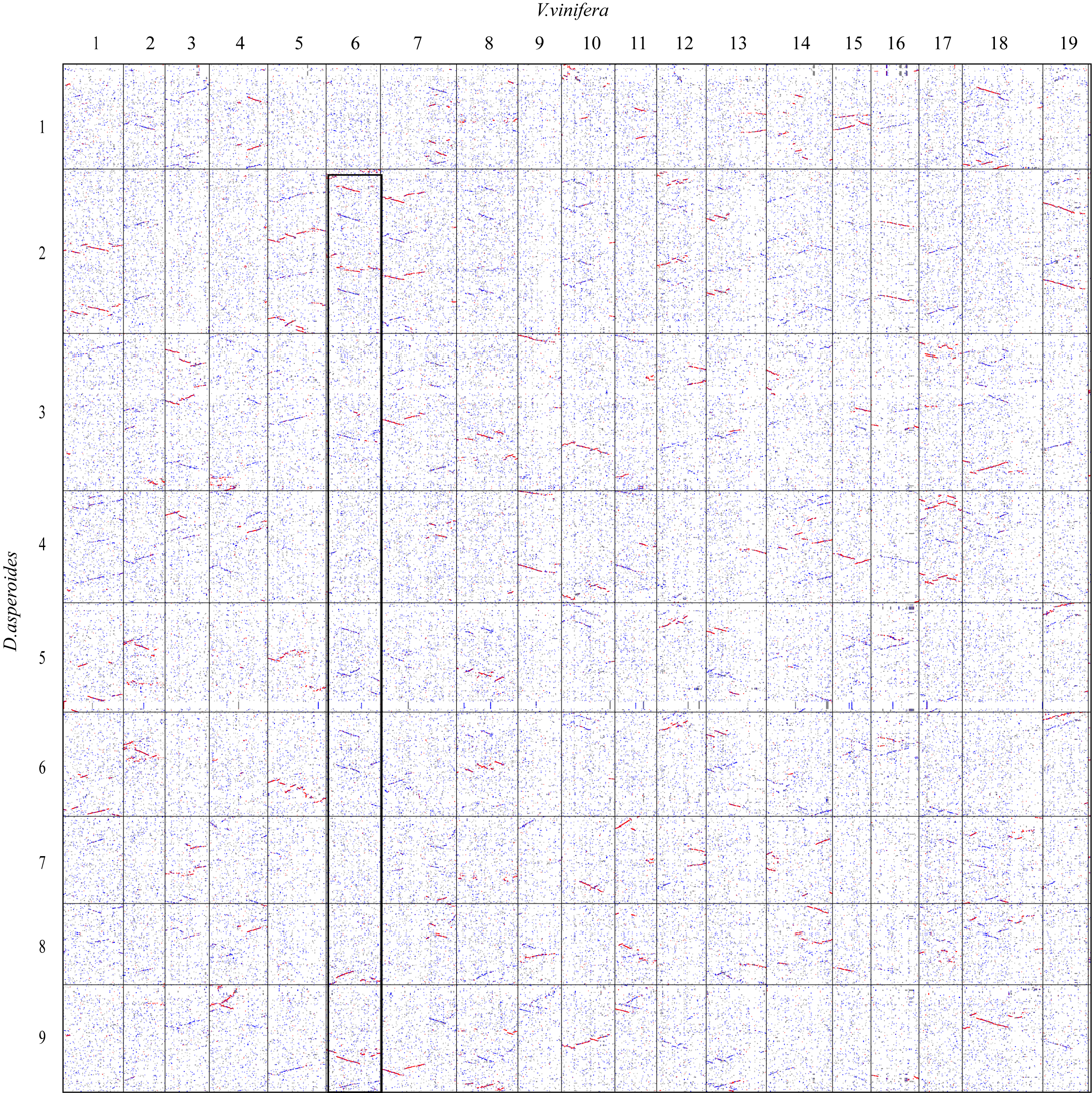

**Supplemental Figure 17. Genome-wide collinearity dot plot between *D. asperoides* and *V. vinifera***.

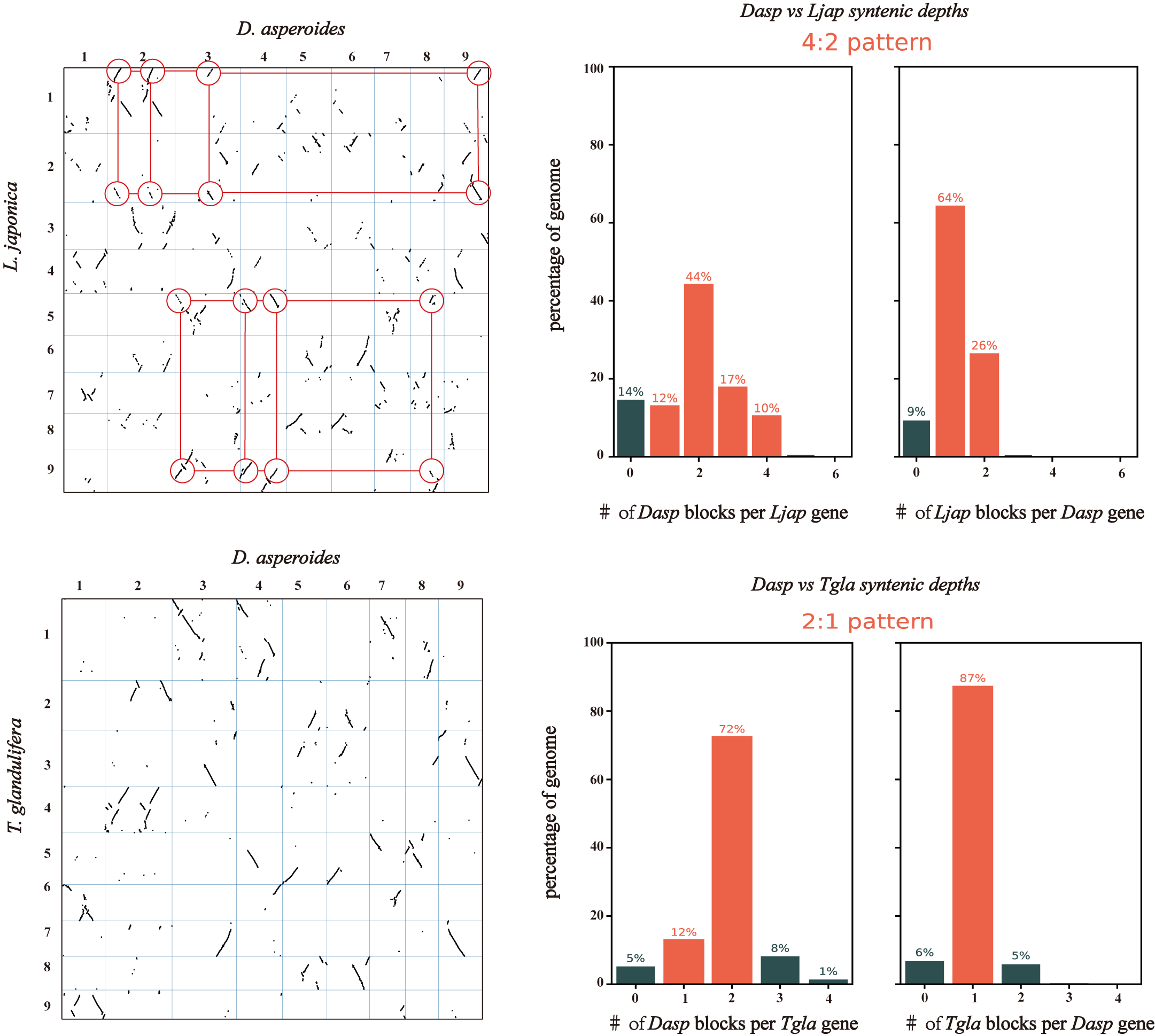

**Supplemental Figure 18. Dotplot and collinearity depth analysis of *D. asperoides* and the other two Caprifoliaceae species (*L. japonica* and *T. glandulifera*)**.

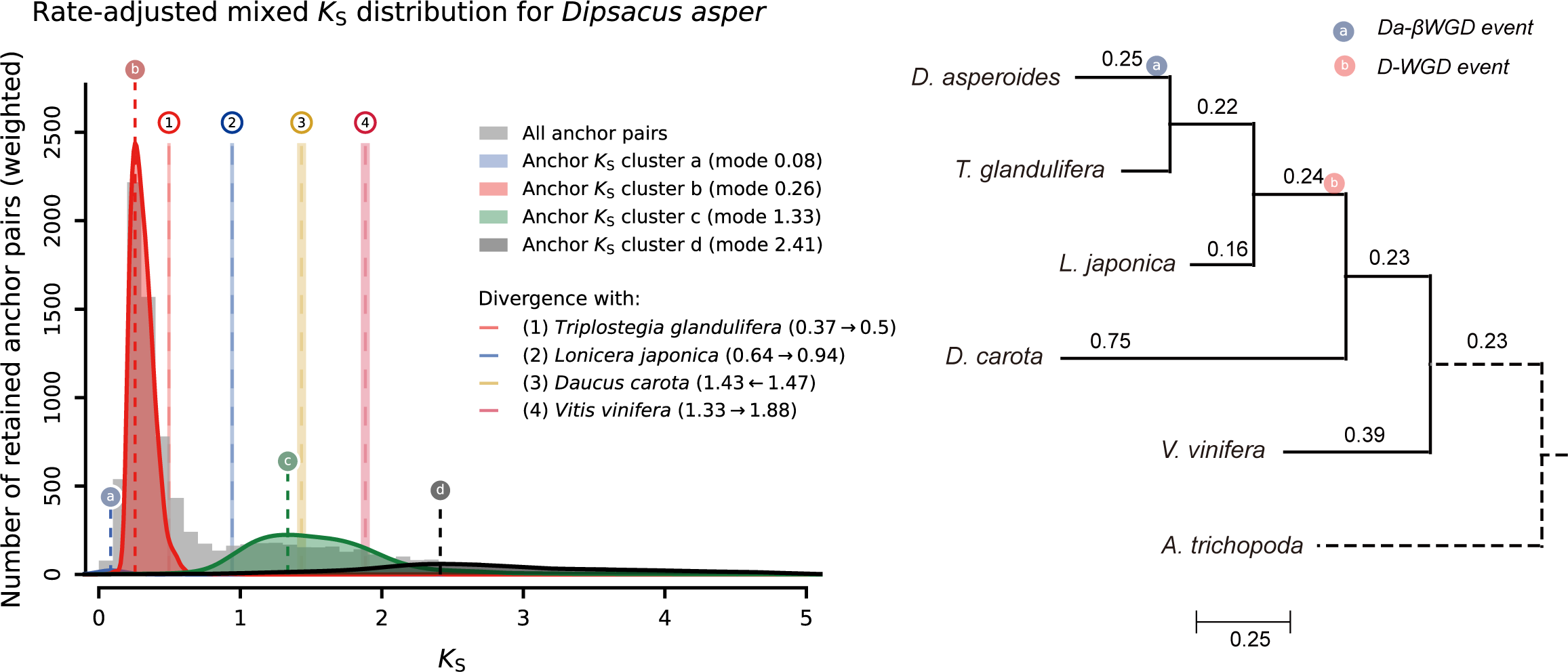

**Supplemental Figure 19. The mixed paralog-ortholog *Ks* plot after each median has been substituted by its original anchor-pair *Ks* list**.

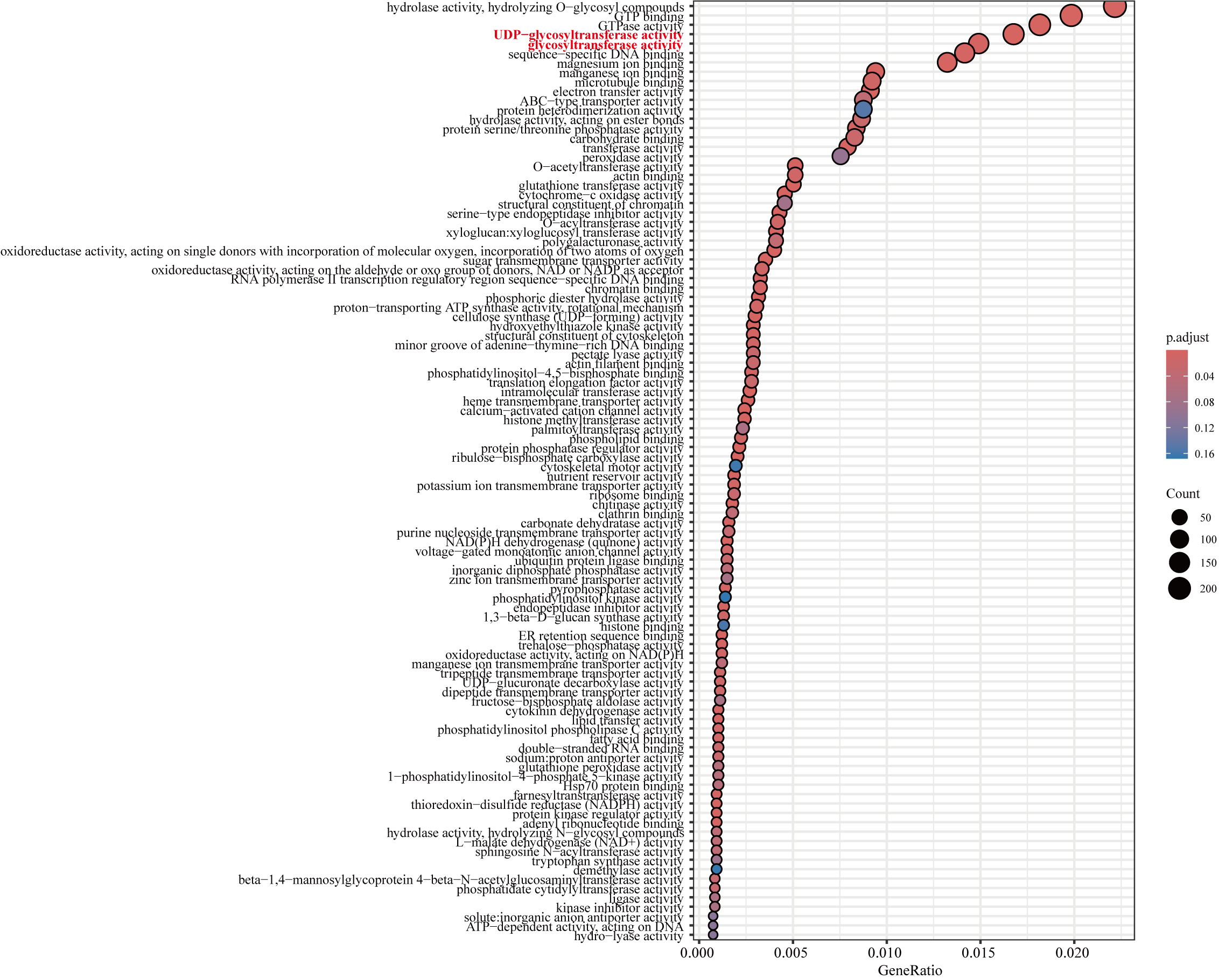

**Supplemental Figure 20. GO enrichment for the molecular function (MF) category of expanded gene families in *D. asperoides***.

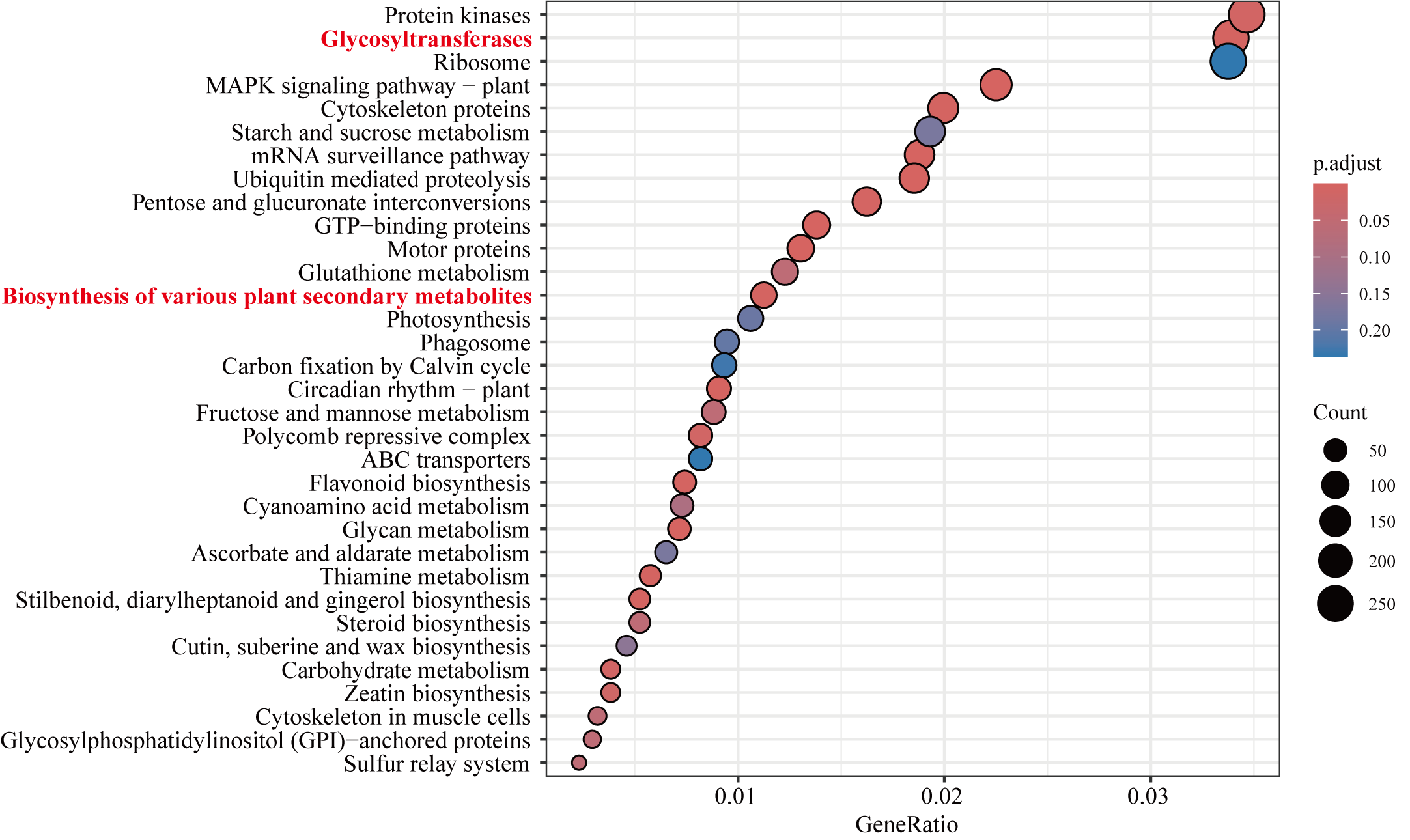

**Supplemental Figure 21. KEGG enrichment of expanded gene families in *D. asperoides***.

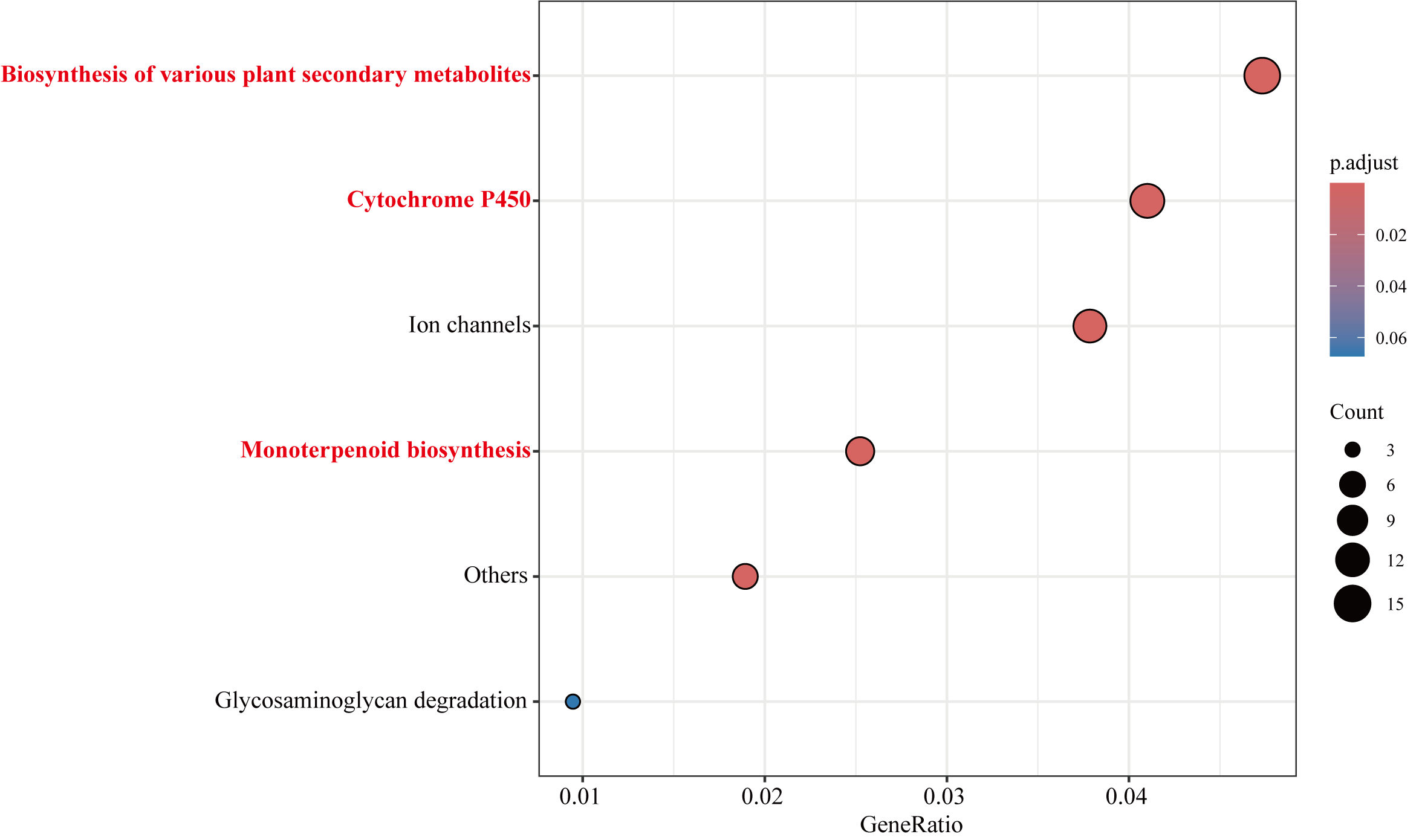

**Supplemental Figure 22. KEGG enrichment of contracted gene families in *D. asperoides***.

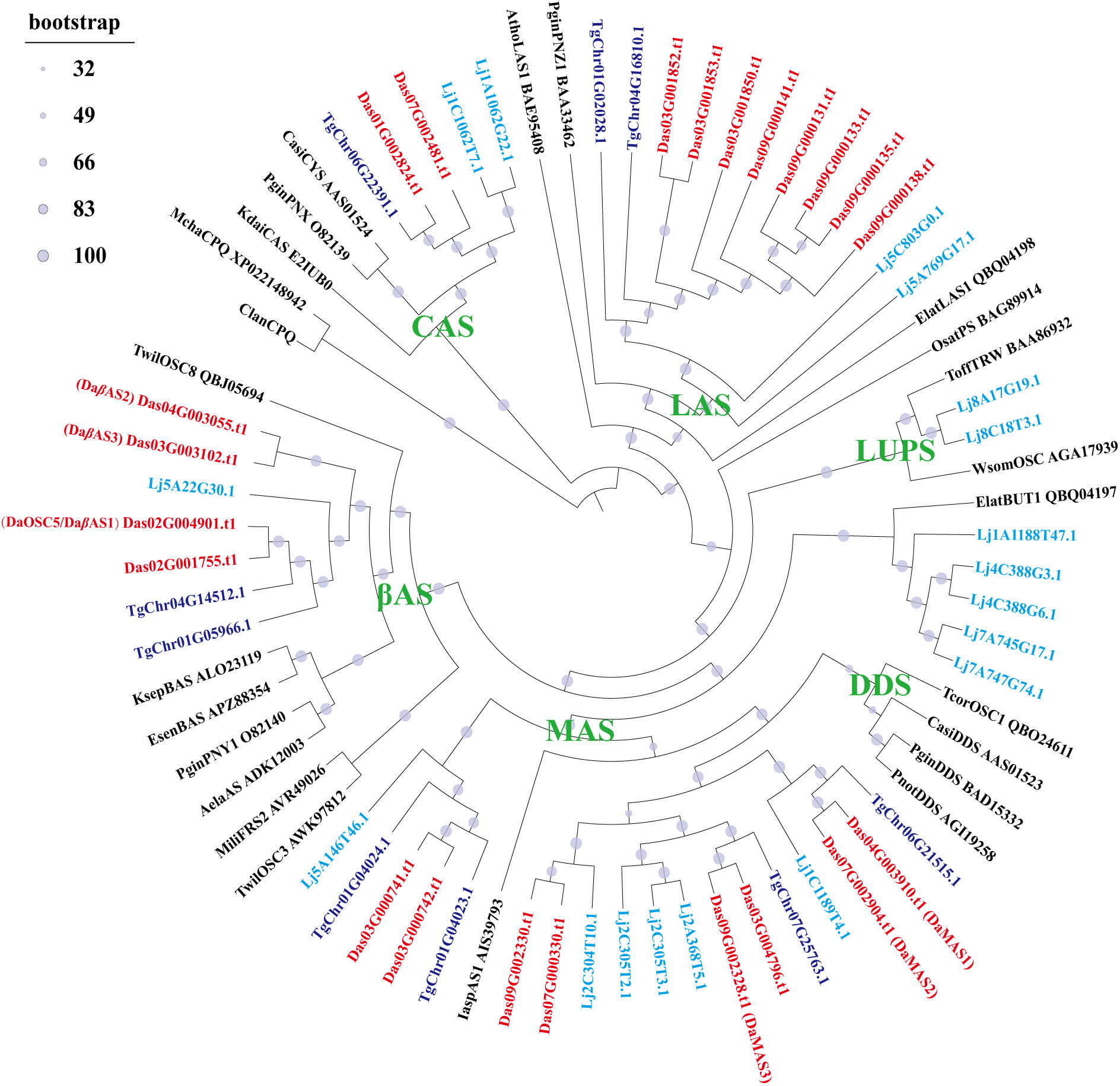

**Supplemental Figure 23. Phylogenetic tree of OSC family proteins in *D. asperoides*, *L. japonica*, and *T. glandulifera***. The phylogenetic tree was constructed using the maximum likelihood method, in which tree nodes (>50%) are indicated by purple dots on branches (1000 bootstrap replications).

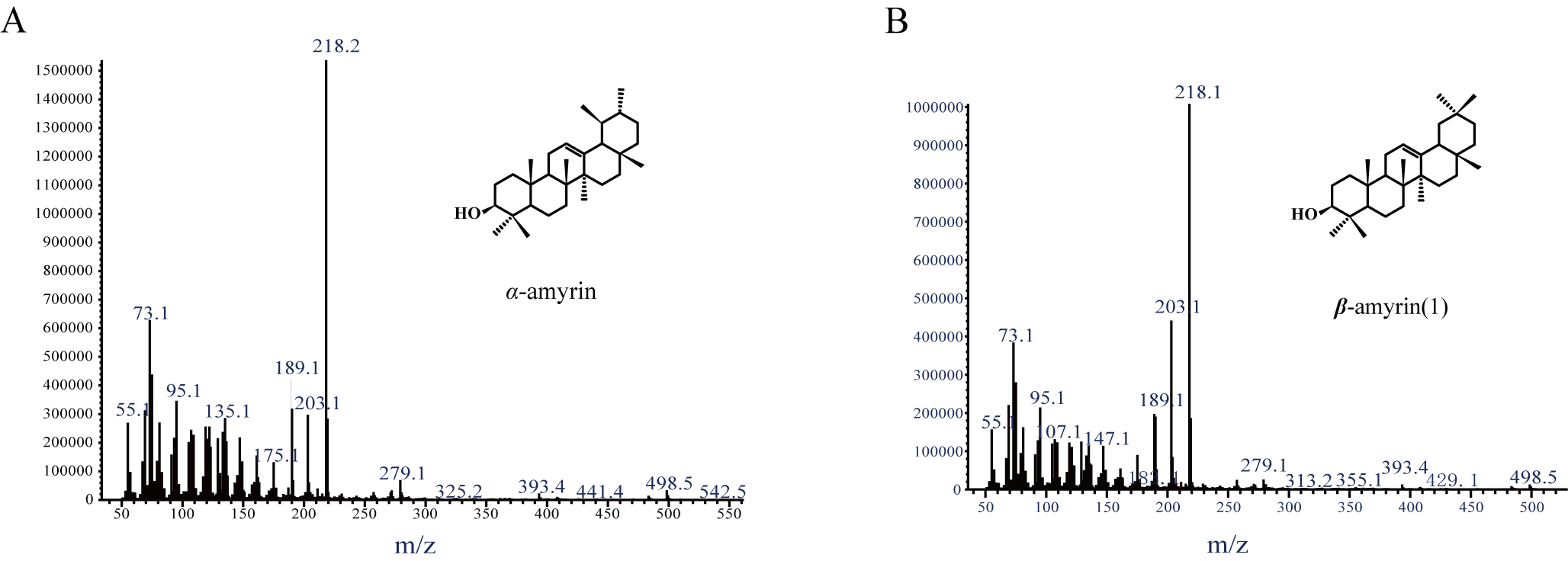

**Supplemental Figure 24. Mass spectra identification for *α*-amyrin and *β*-amyrin**.

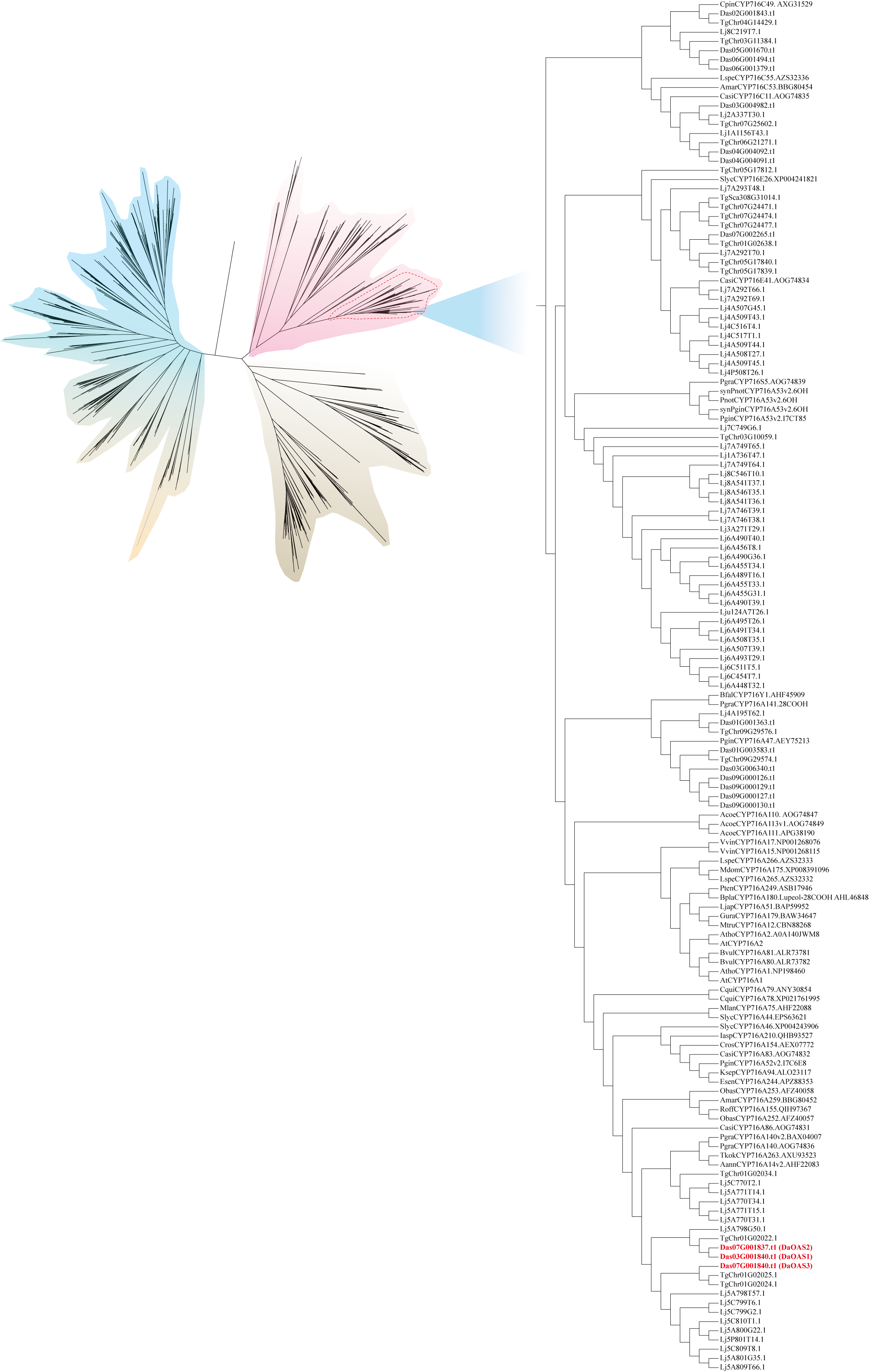

**Supplemental Figure 25. Phylogenetic tree of CYP716 family proteins in *D. asperoides*, *L. japonica*, and *T. glandulifera***. The phylogenetic tree was constructed using the maximum likelihood method, in which tree nodes (>50%) are indicated by purple dots on branches (1000 bootstrap replications).

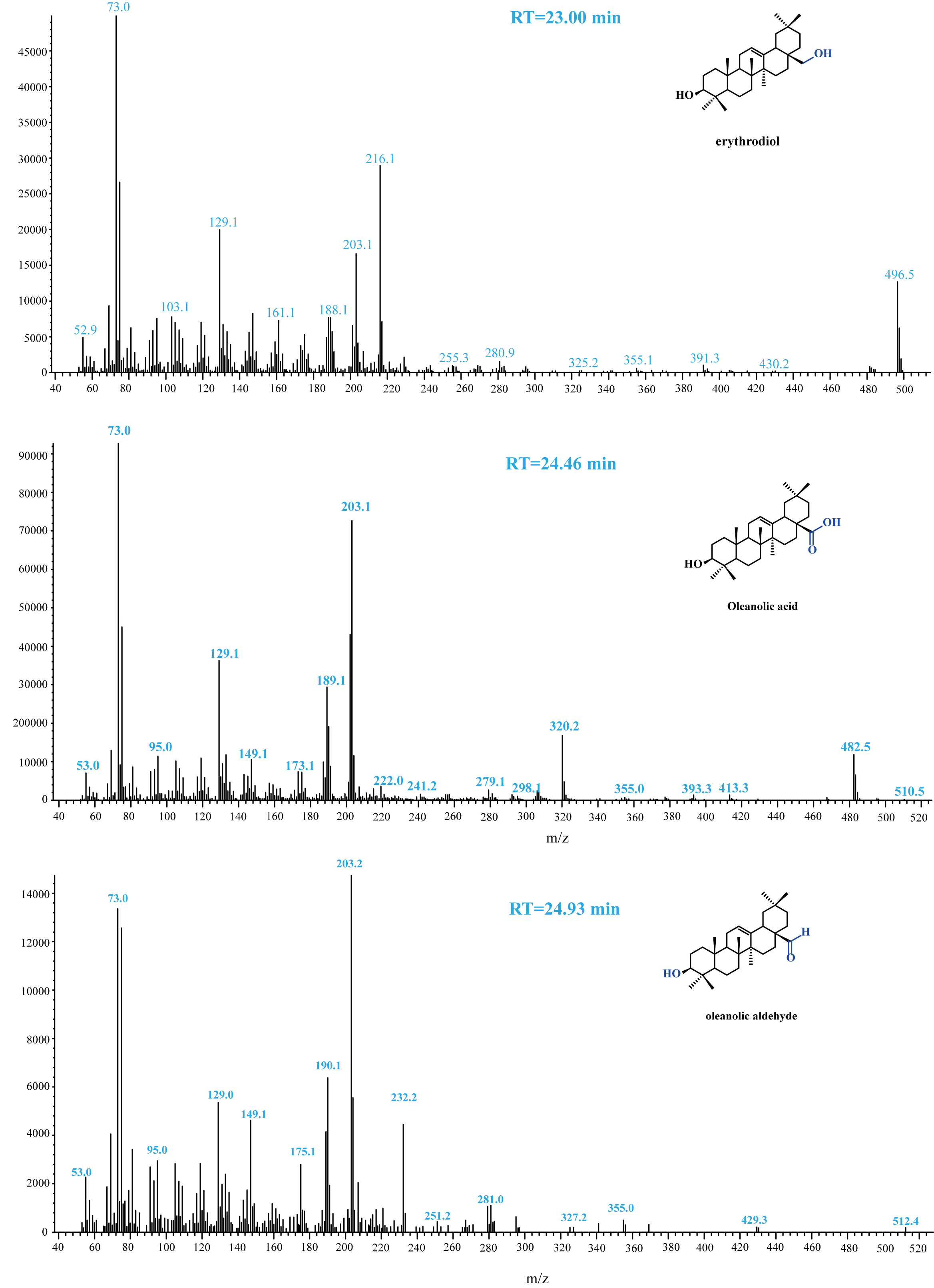

**Supplemental Figure 26. Mass spectra identification for erythrodiol, oleanolic acid and oleanolic aldehyde**.

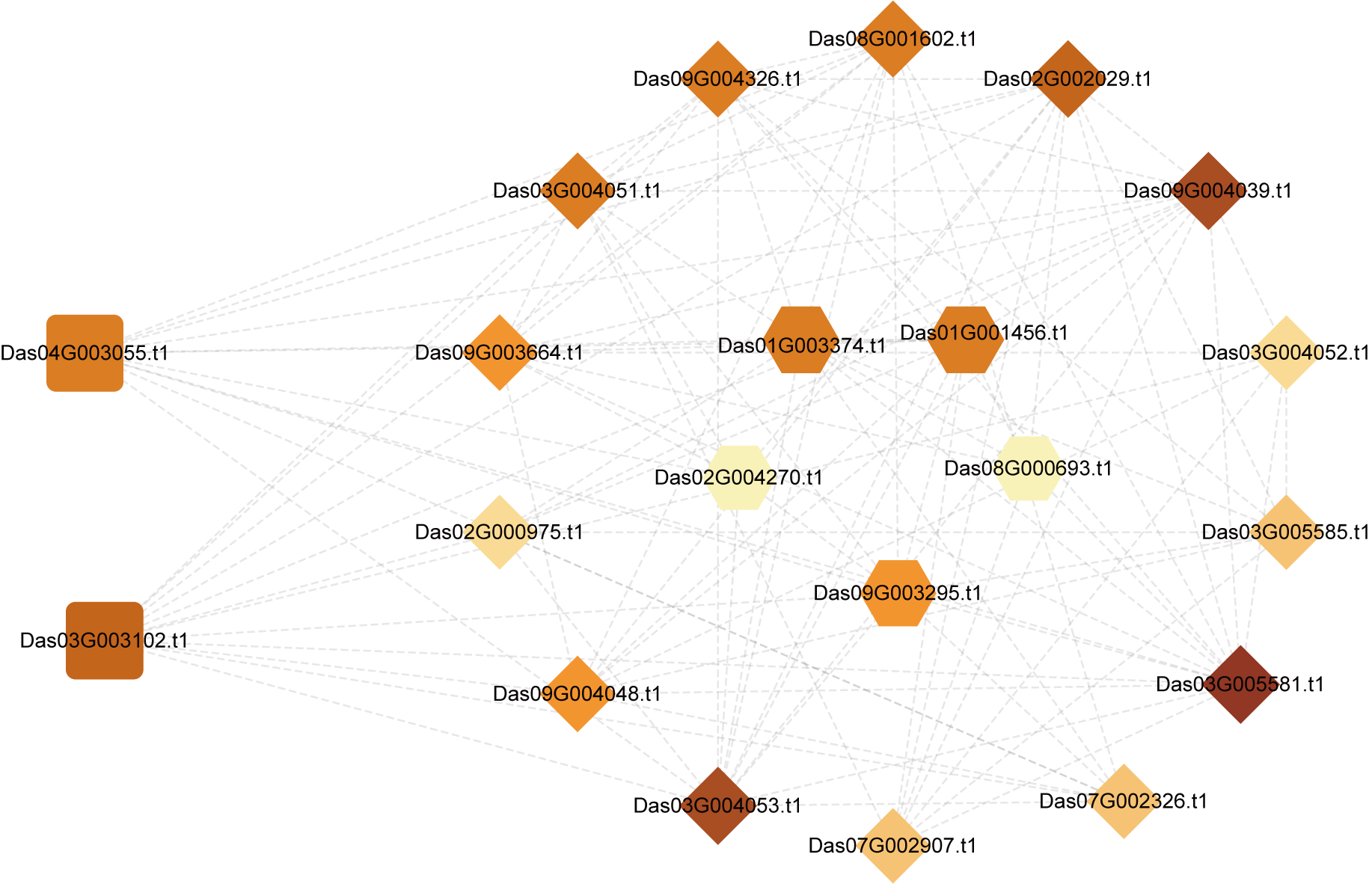

**Supplemental Figure 27.** **Co-expression analysis of *D. asperoides* transcriptome using Das04G003055.t1, Das09G003295.t1 and Das07G002907.t1 as bait genes**. The co-expression result was visualized by Cytoscape.

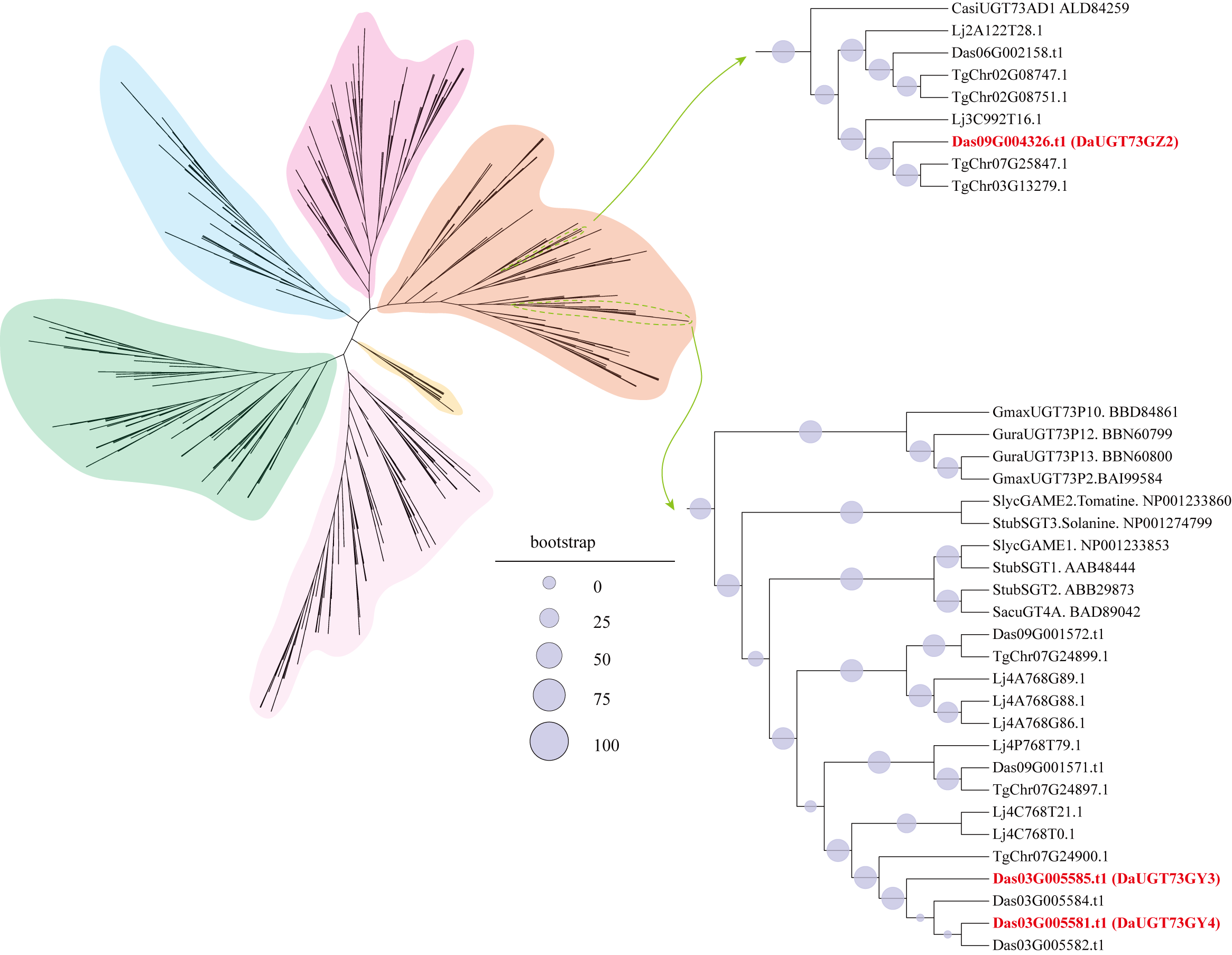

**Supplemental Figure 28.** **Phylogenetic tree of UGT73 family proteins in *D. asperoides*, *L. japonica*, and *T. glandulifera***. The phylogenetic tree was constructed using the maximum likelihood method, in which tree nodes (>50%) are indicated by purple dots on branches (1000 bootstrap replications).

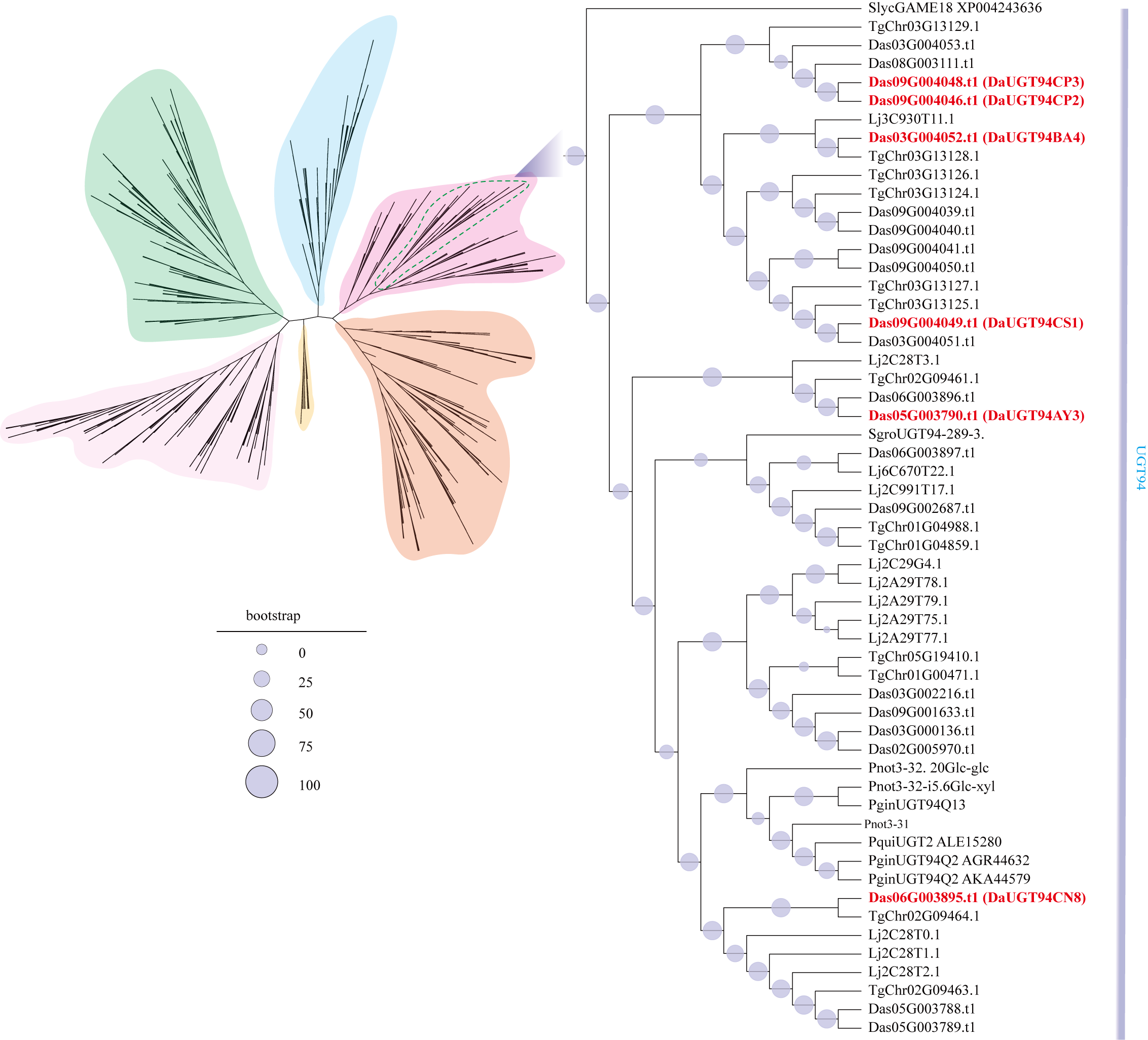

**Supplemental Figure 29.** **Phylogenetic tree of UGT94 family proteins in *D. asperoides*, *L. japonica*, and *T. glandulifera***. The phylogenetic tree was constructed using the maximum likelihood method, in which tree nodes (>50%) are indicated by purple dots on branches (1000 bootstrap replications).

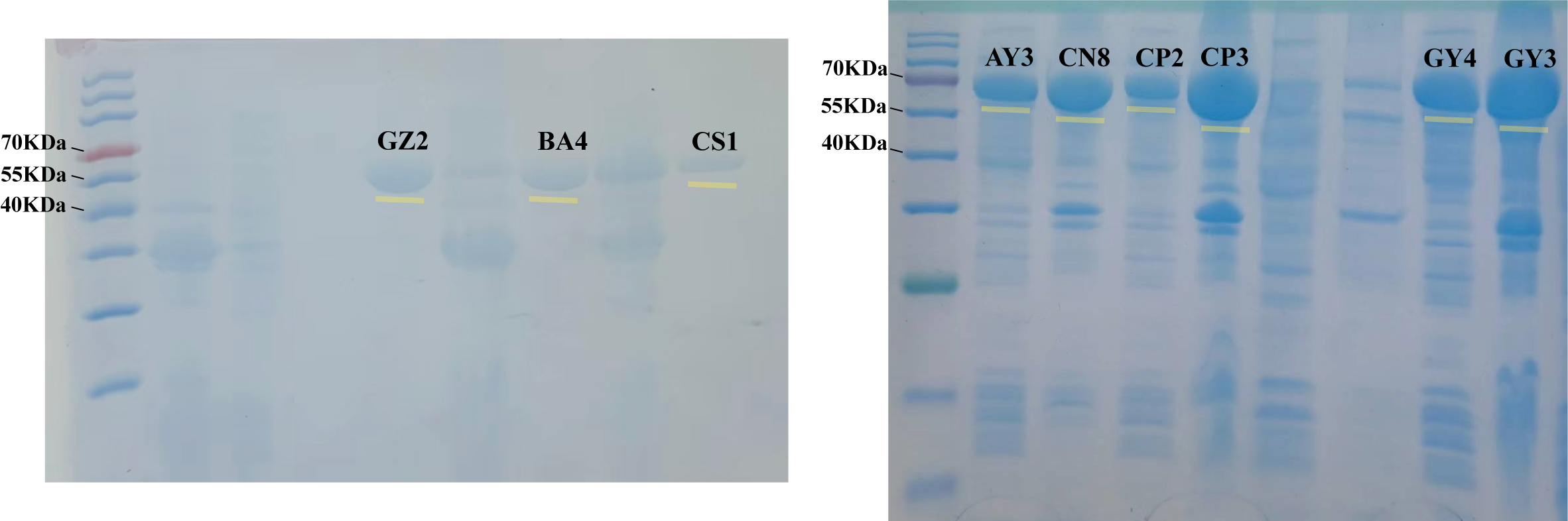

**Supplemental Figure 30. SDS-PAGE analysis of recombinant UGT proteins heterologously expressed in *E. coli*. Lane M: protein molecular weight marker**. The numbered lanes correspond to the purified DaUGT candidates, including DaUGT73GZ2, DaUGT94BA4, DaUGT94CS1, DaUGT94AY3, DaUGT94CN8, DaUGT94CP2, DaUGT94CP3, DaUGT73GY4, and DaUGT73GY3. Yellow bars indicate the specific bands of the target proteins.

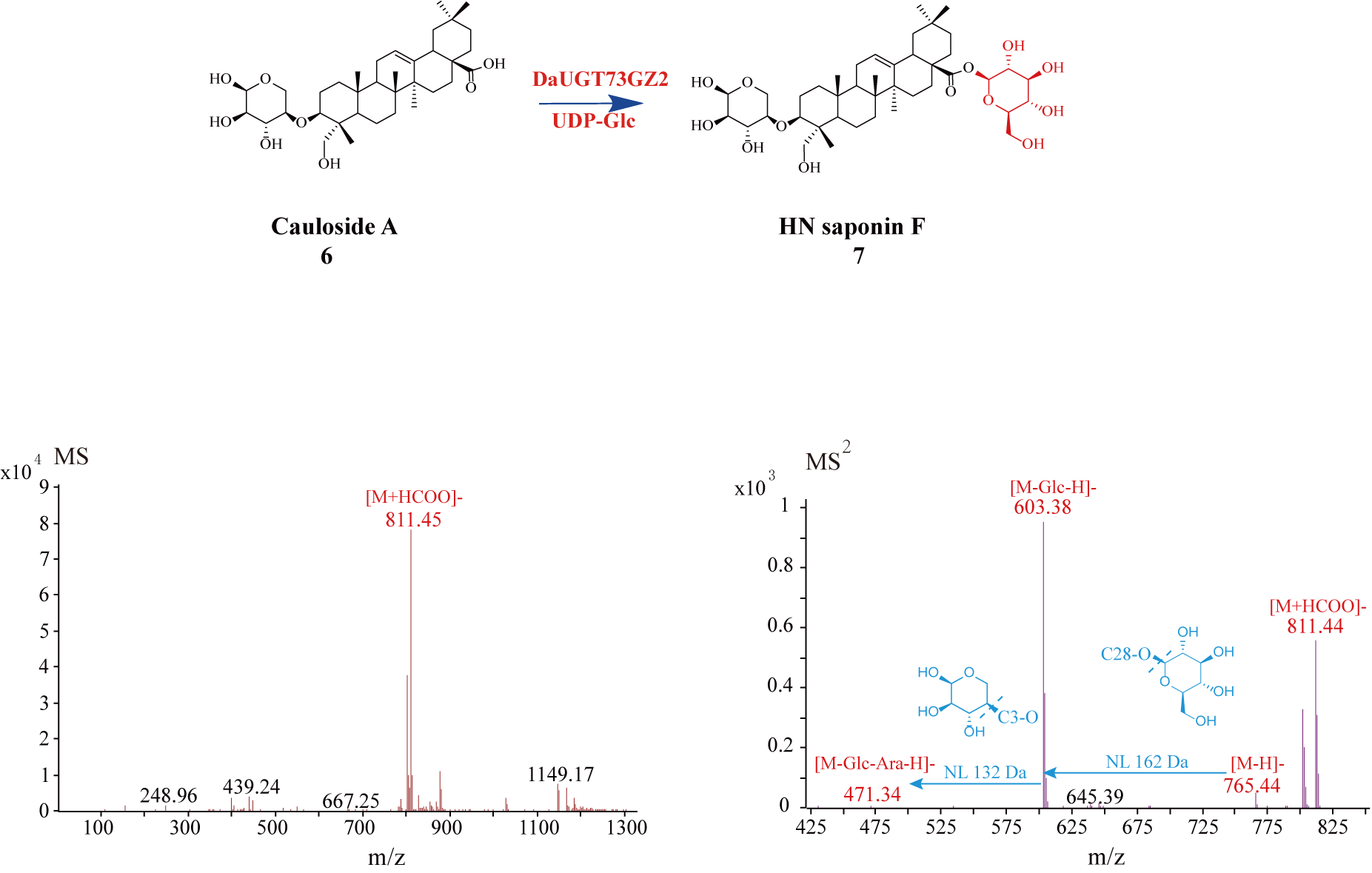
 **Supplemental Figure 31. The (-)-ESI-MS and MS/MS spectra of HN saponin F**.

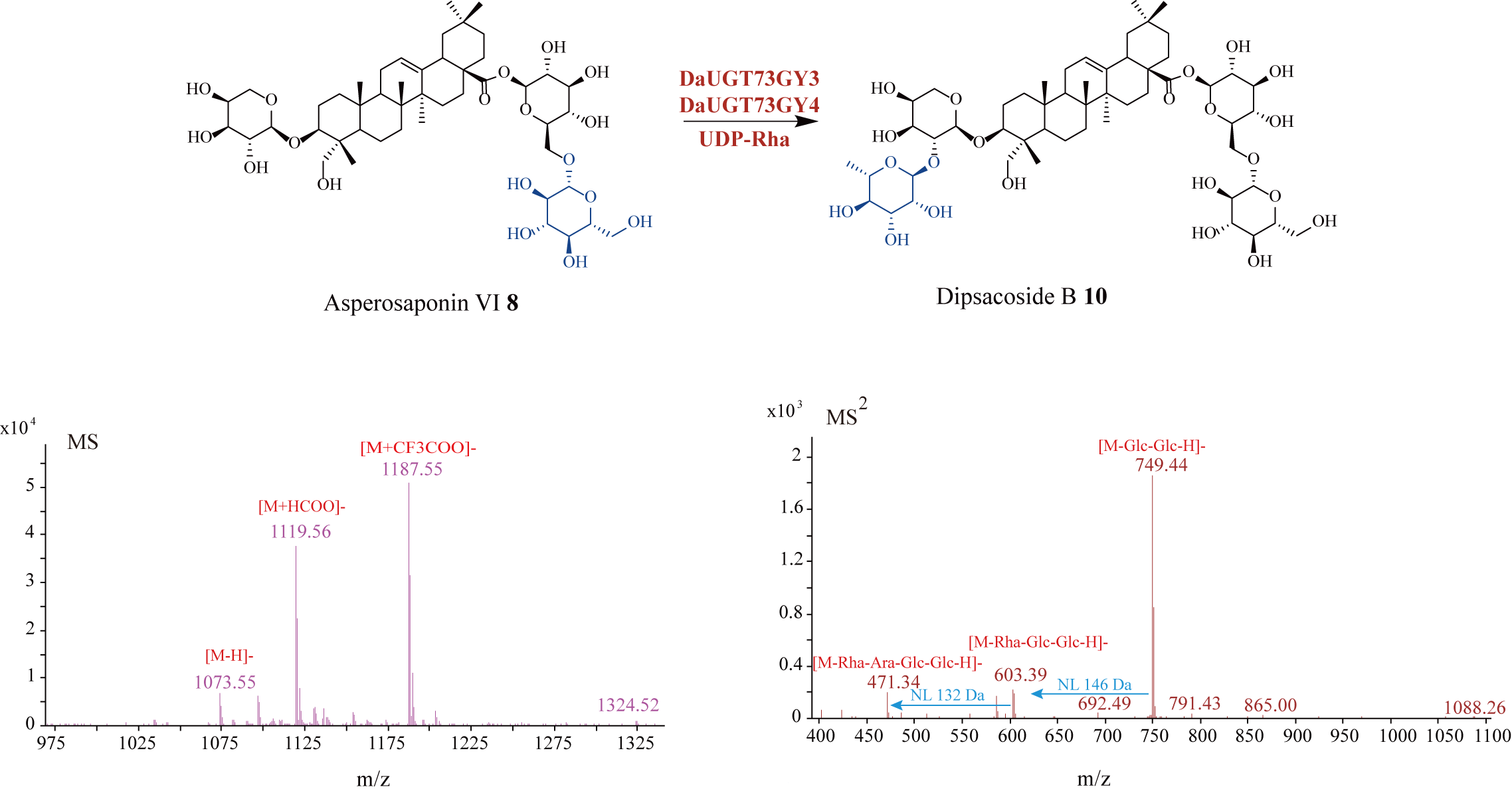

**Supplemental Figure 32. The (-)-ESI-MS and MS/MS spectra of dipsacoside B**.

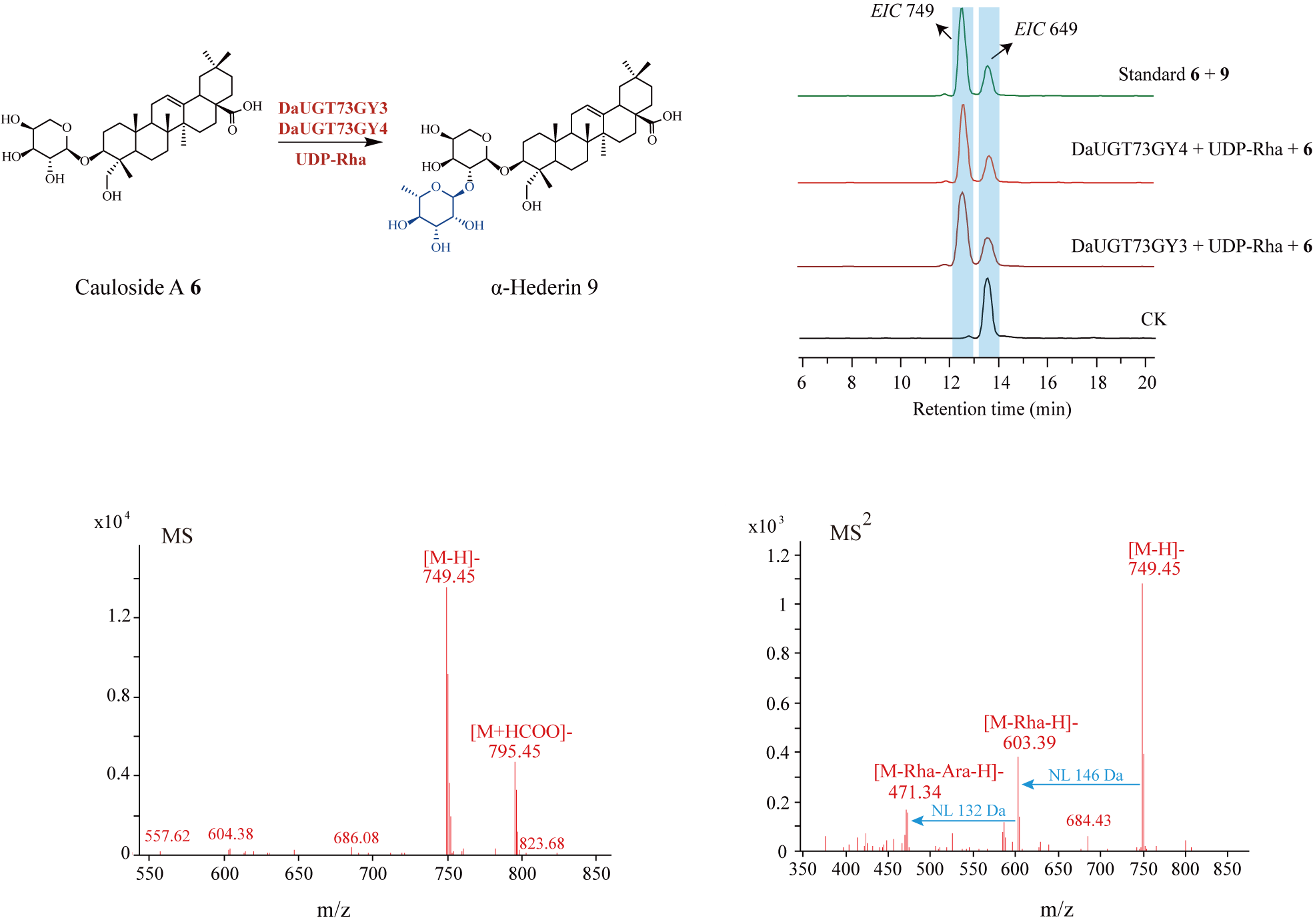

**Supplemental Figure 33. Functional characterization of DaUGT73GY4 and DaUGT73GY3 catalyzing the conversion of compound 6 to compound 9**.

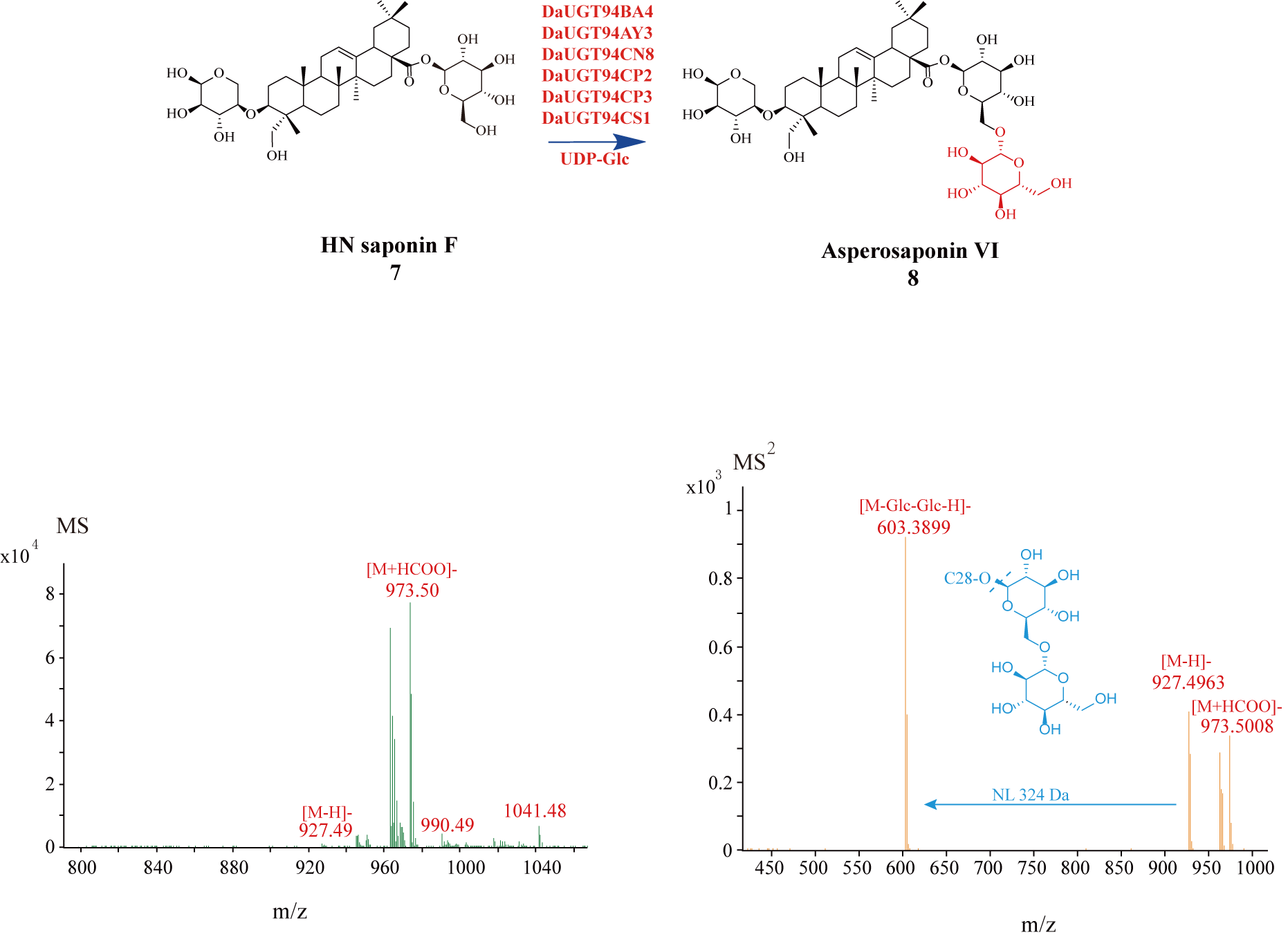
 **Supplemental Figure 34. The (-)-ESI-MS and MS/MS spectra of asperosaponin Ⅵ**.

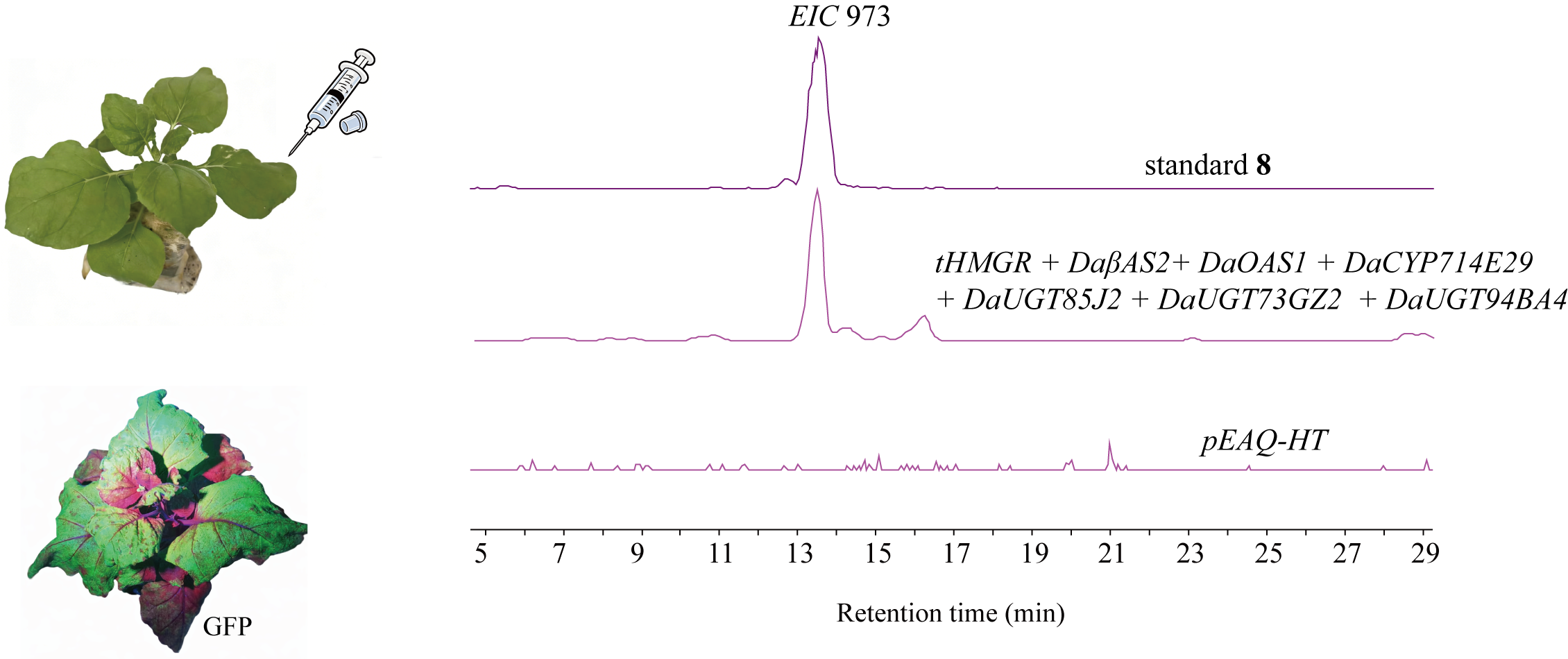

**Supplemental Figure 35. De novo biosynthesis of asperosaponin VI (8) in *N. benthamiana* leaves**.

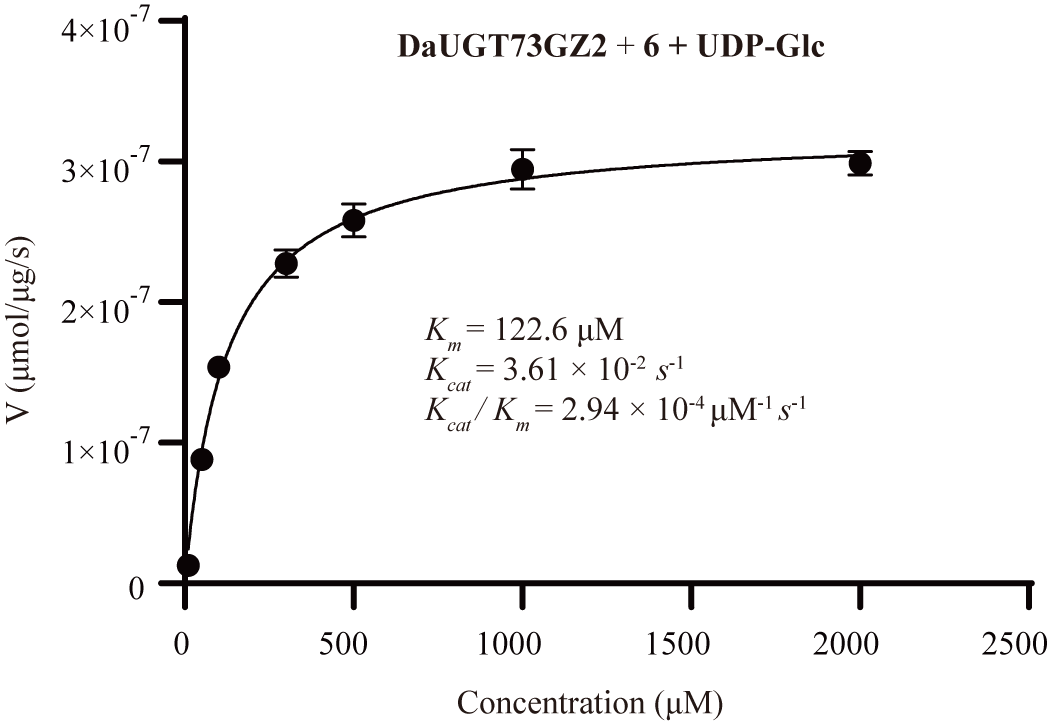

**Supplemental Figure 36. Enzyme kinetics of DaUGT73GZ2**.

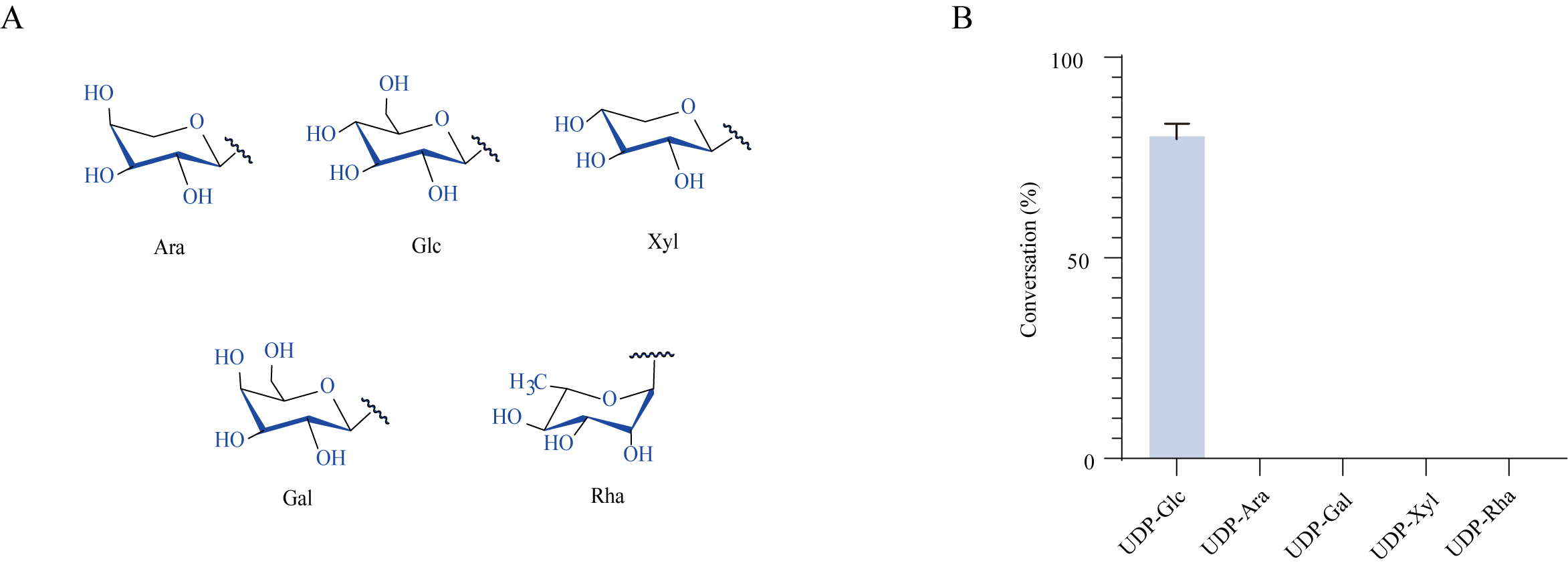

**Supplemental Figure 37. Sugar donor specificity of DaUGT73GZ2**. **(A)** Chemical structures of sugar donors tested in this study. **(B)** Conversion rates (%) of **6** catalyzed by DaUGT73GZ2 using different sugar donors.

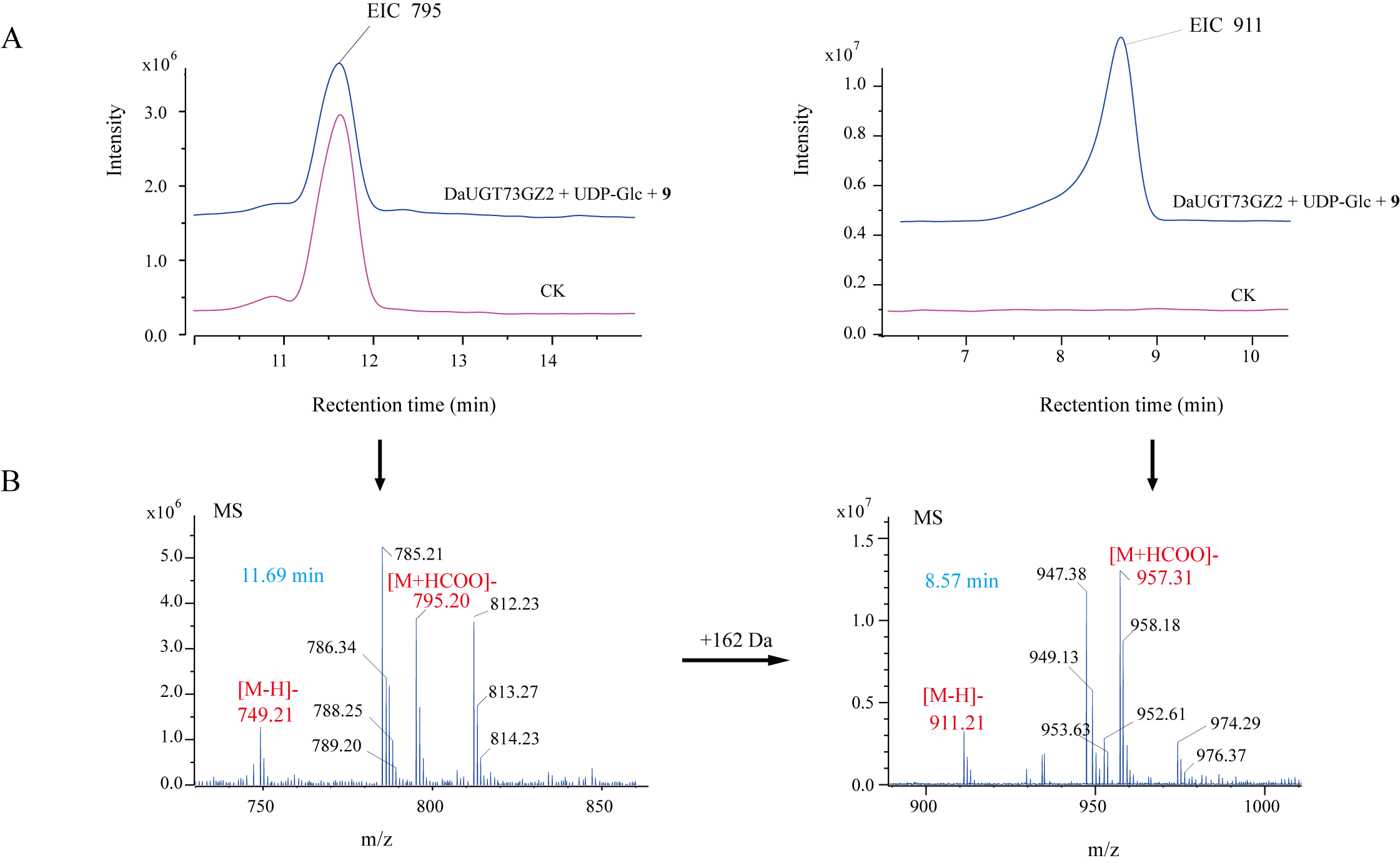

**Supplemental Figure 38. LC-MS analysis of the reaction products of 9 and UDP-Glc catalyzed by DaUGT73GZ2**. **(A)** Extracted ion chromatograms (EICs) from UPLC-MS analysis. **(B)** LC-MS spectra for glycoside products.

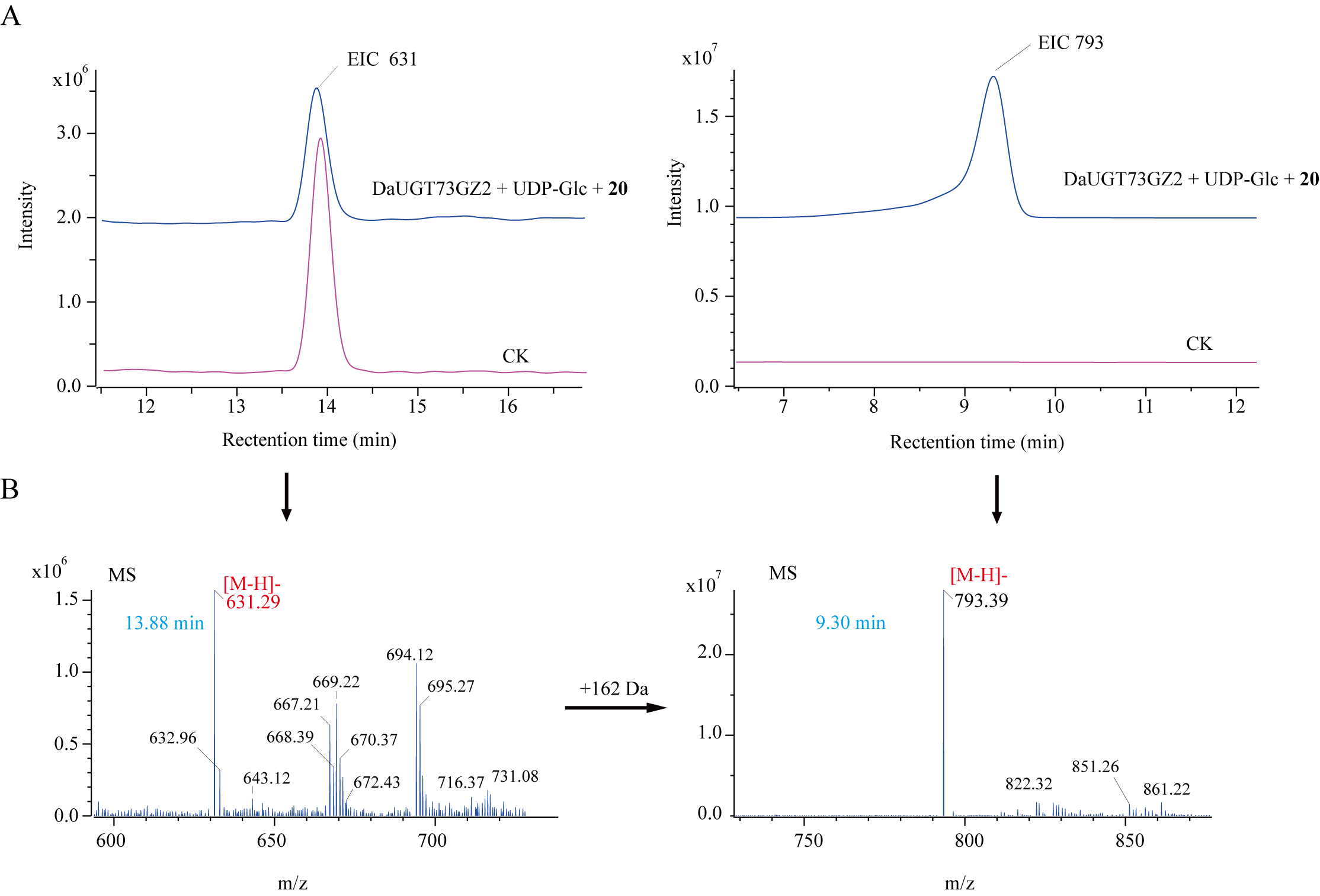

**Supplemental Figure 39. LC-MS analysis of the reaction products of 20 and UDP-Glc catalyzed by DaUGT73GZ2**. **(A)** Extracted ion chromatograms (EICs) from UPLC-MS analysis. **(B)** LC-MS spectra for glycoside products.

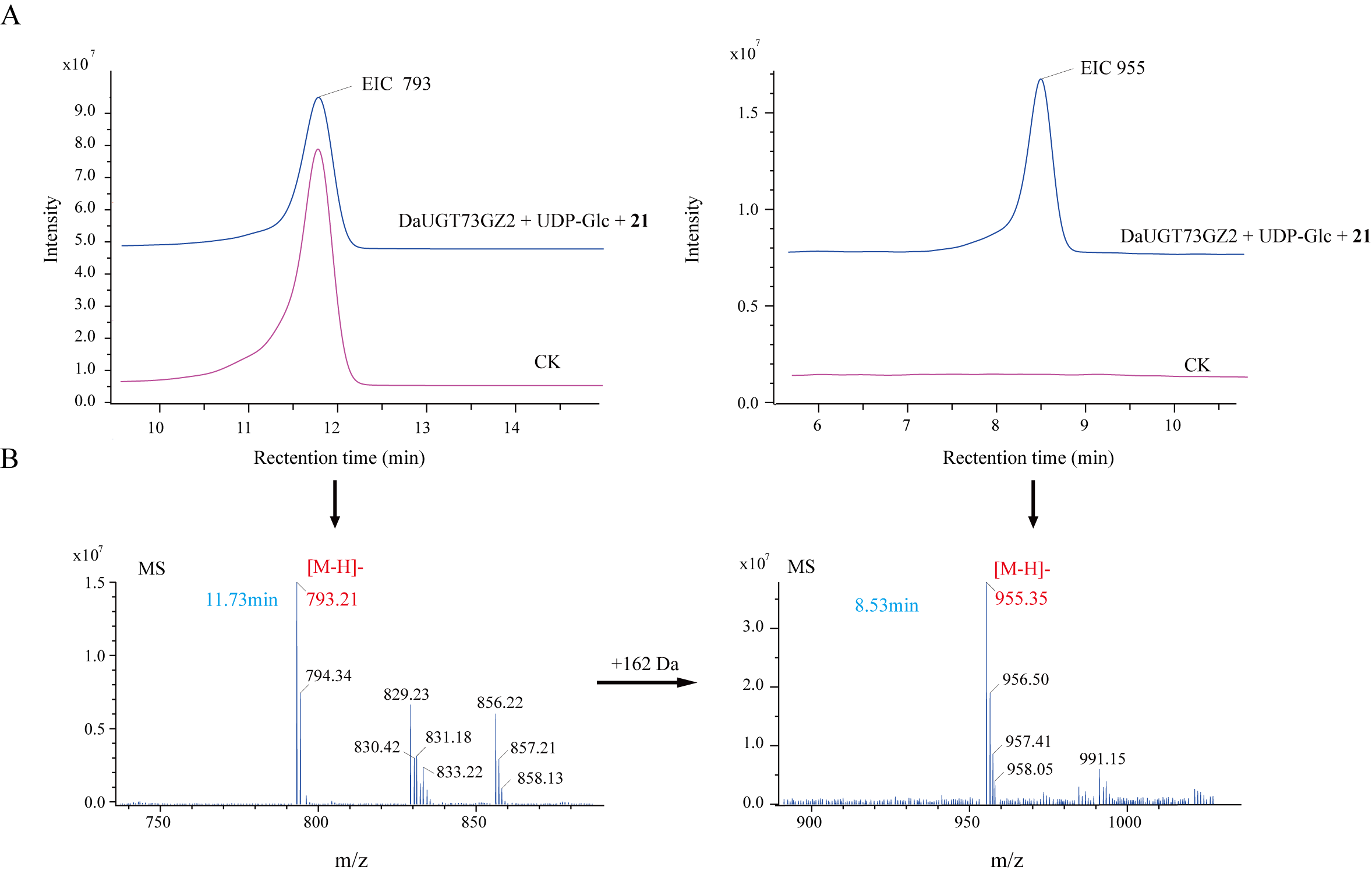

**Supplemental Figure 40. LC-MS analysis of the reaction products of 21 and UDP-Glc catalyzed by DaUGT73GZ2**. **(A)** Extracted ion chromatograms (EICs) from UPLC-MS analysis. **(B)** LC-MS spectra for glycoside products.

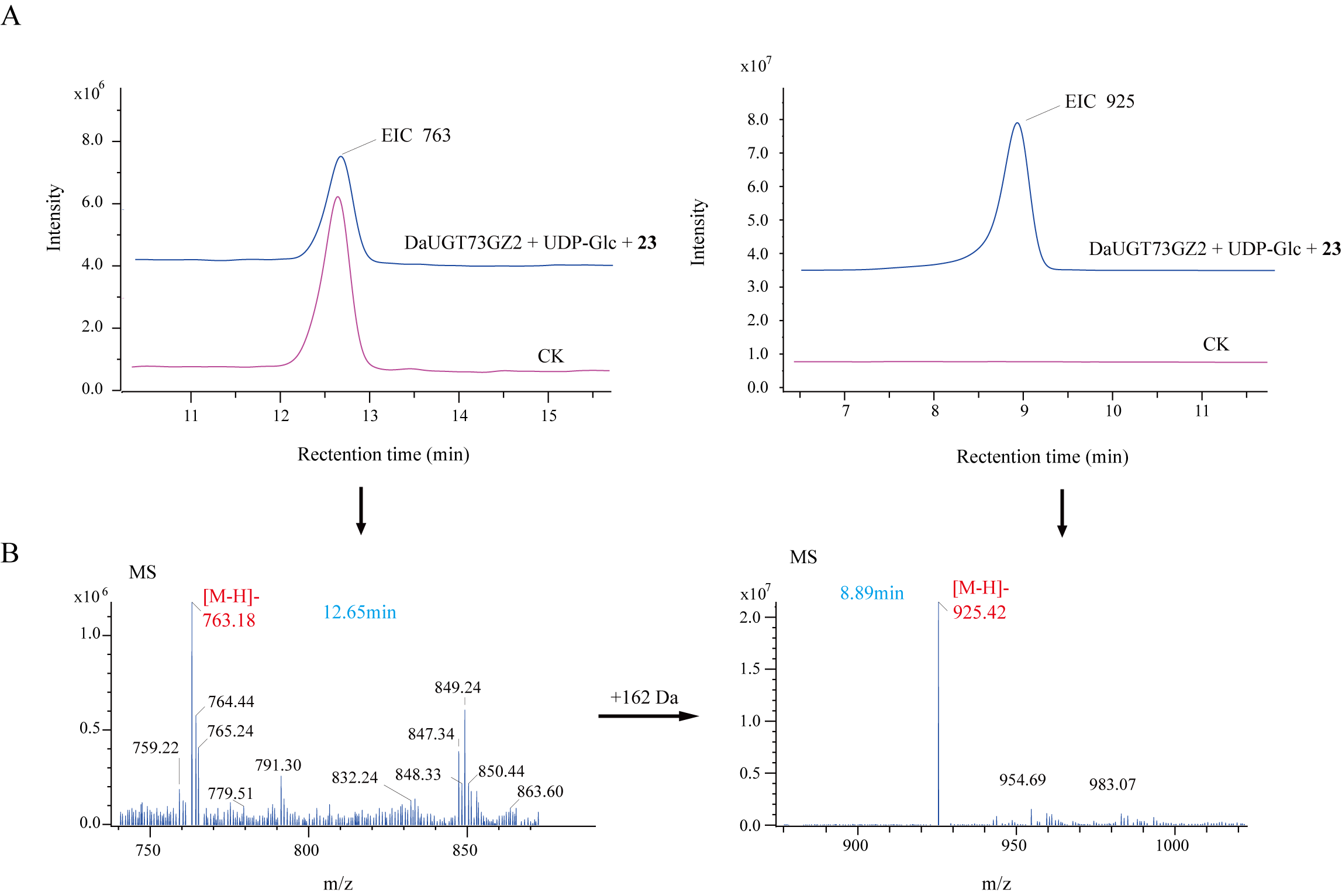

**Supplemental Figure 41. LC-MS analysis of the reaction products of 23 and UDP-Glc catalyzed by DaUGT73GZ2**. **(A)** Extracted ion chromatograms (EICs) from UPLC-MS analysis. **(B)** LC-MS spectra for glycoside products.

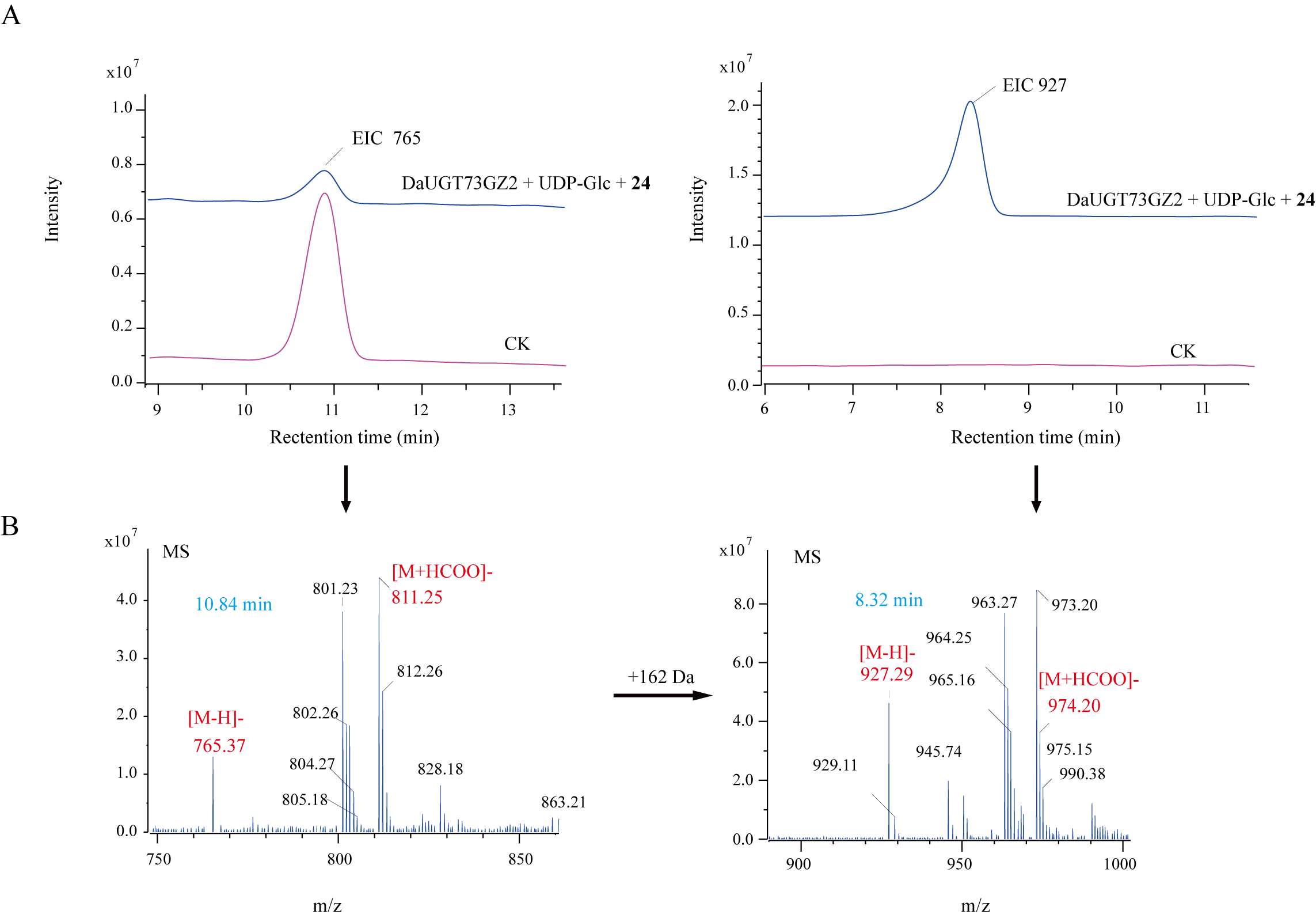

**Supplemental Figure 42. LC-MS analysis of the reaction products of 24 and UDP-Glc catalyzed by DaUGT73GZ2**. **(A)** Extracted ion chromatograms (EICs) from UPLC-MS analysis. **(B)** LC-MS spectra for glycoside products.

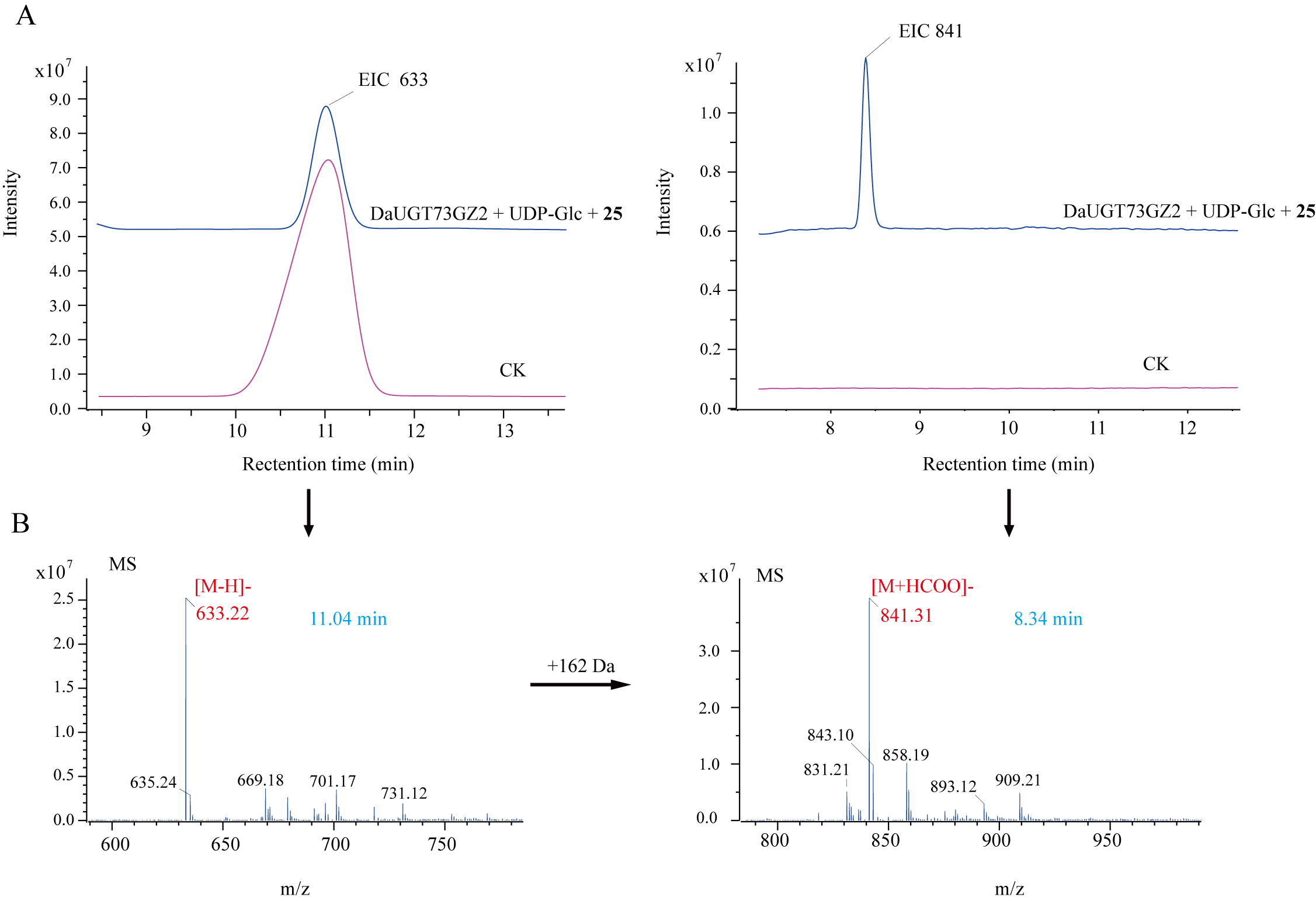

**Supplemental Figure 43. LC-MS analysis of the reaction products of 25 and UDP-Glc catalyzed by DaUGT73GZ2**. **(A)** Extracted ion chromatograms (EICs) from UPLC-MS analysis. **(B)** LC-MS spectra for glycoside products.

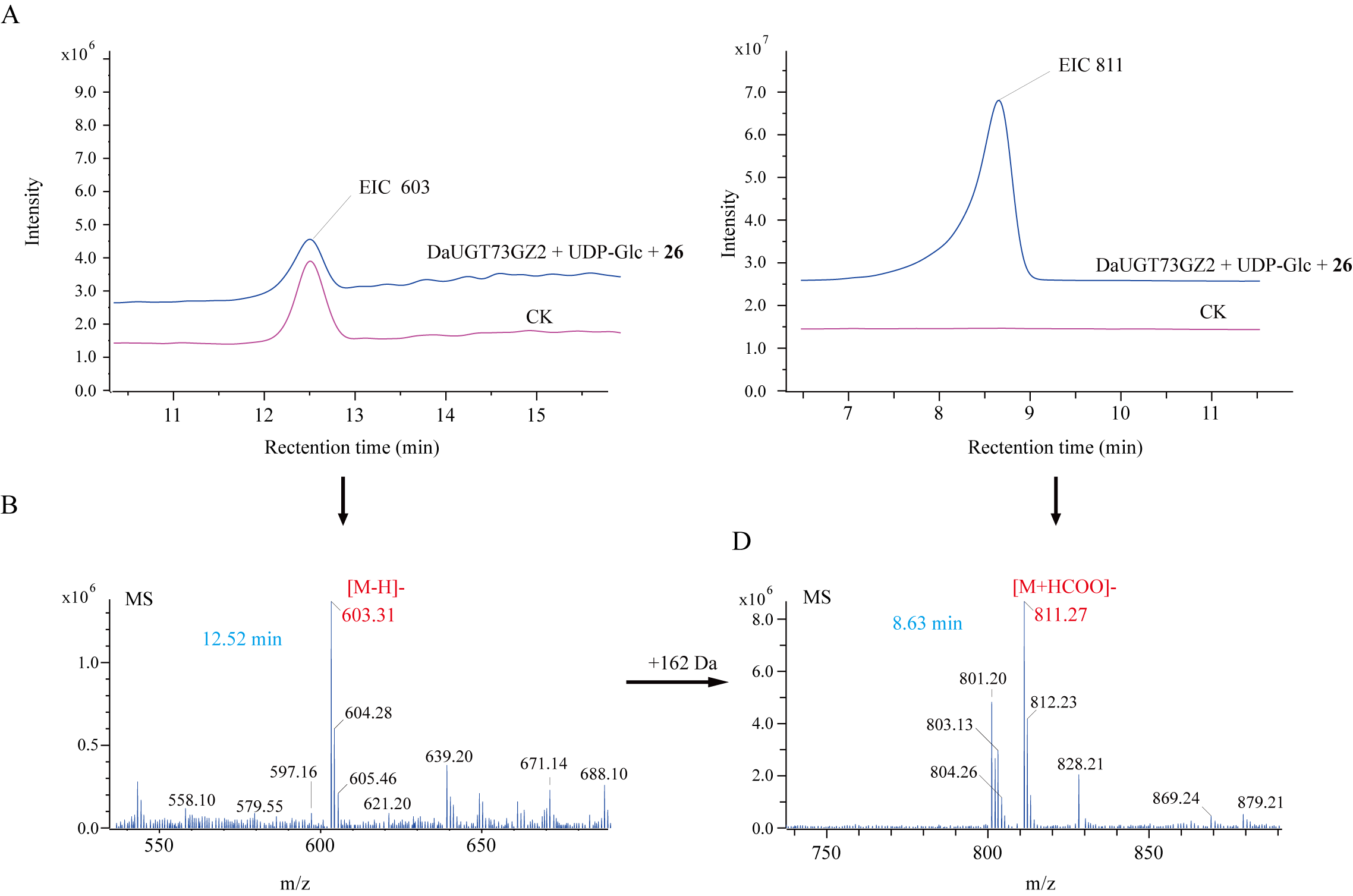

**Supplemental Figure 44. LC-MS analysis of the reaction products of 26 and UDP-Glc catalyzed by DaUGT73GZ2**. **(A)** Extracted ion chromatograms (EICs) from UPLC-MS analysis. **(B)** LC-MS spectra for glycoside products.

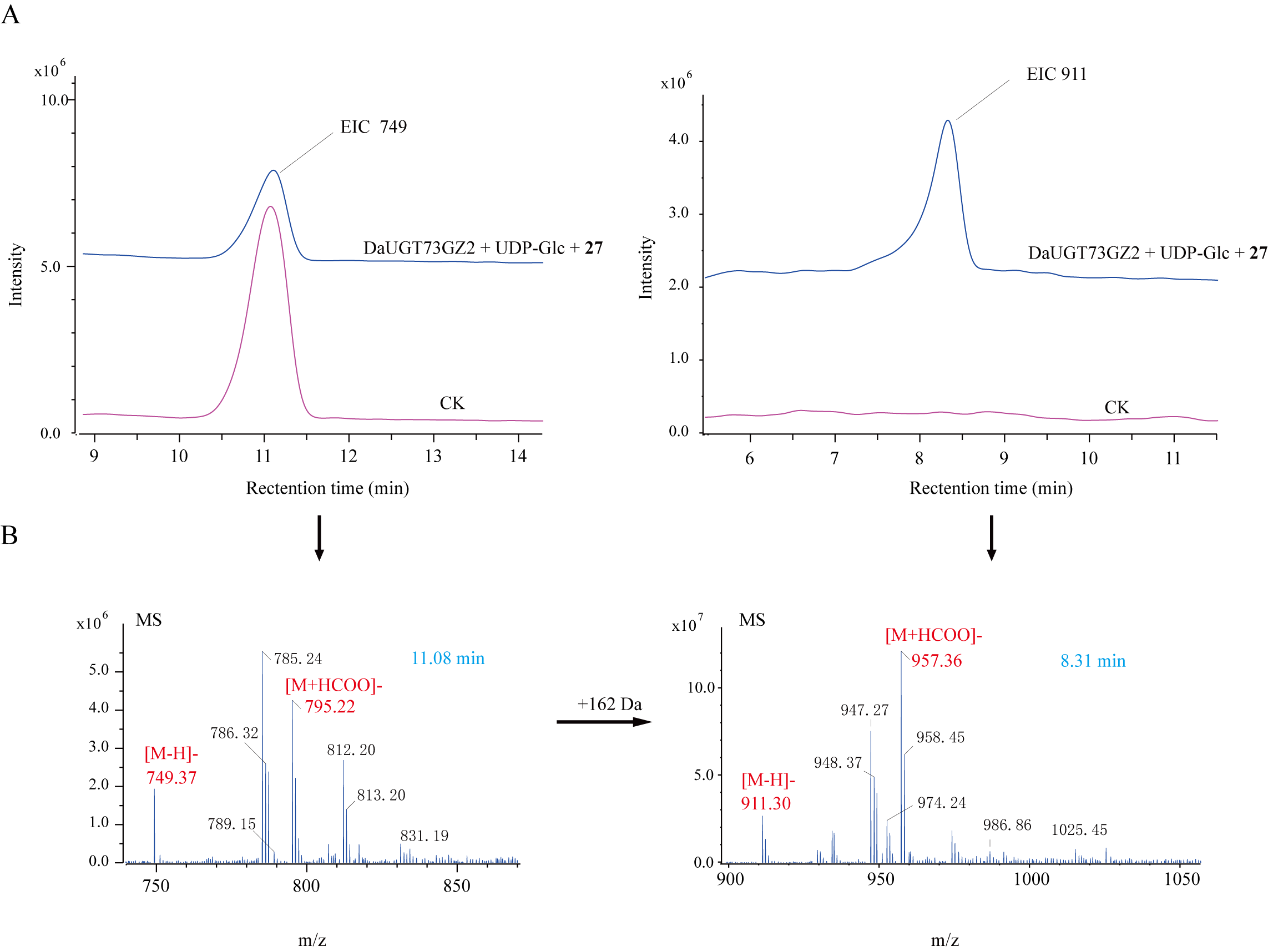

**Supplemental Figure 45. LC-MS analysis of the reaction products of 27 and UDP-Glc catalyzed by DaUGT73GZ2**. **(A)** Extracted ion chromatograms (EICs) from UPLC-MS analysis. **(B)** LC-MS spectra for glycoside products.

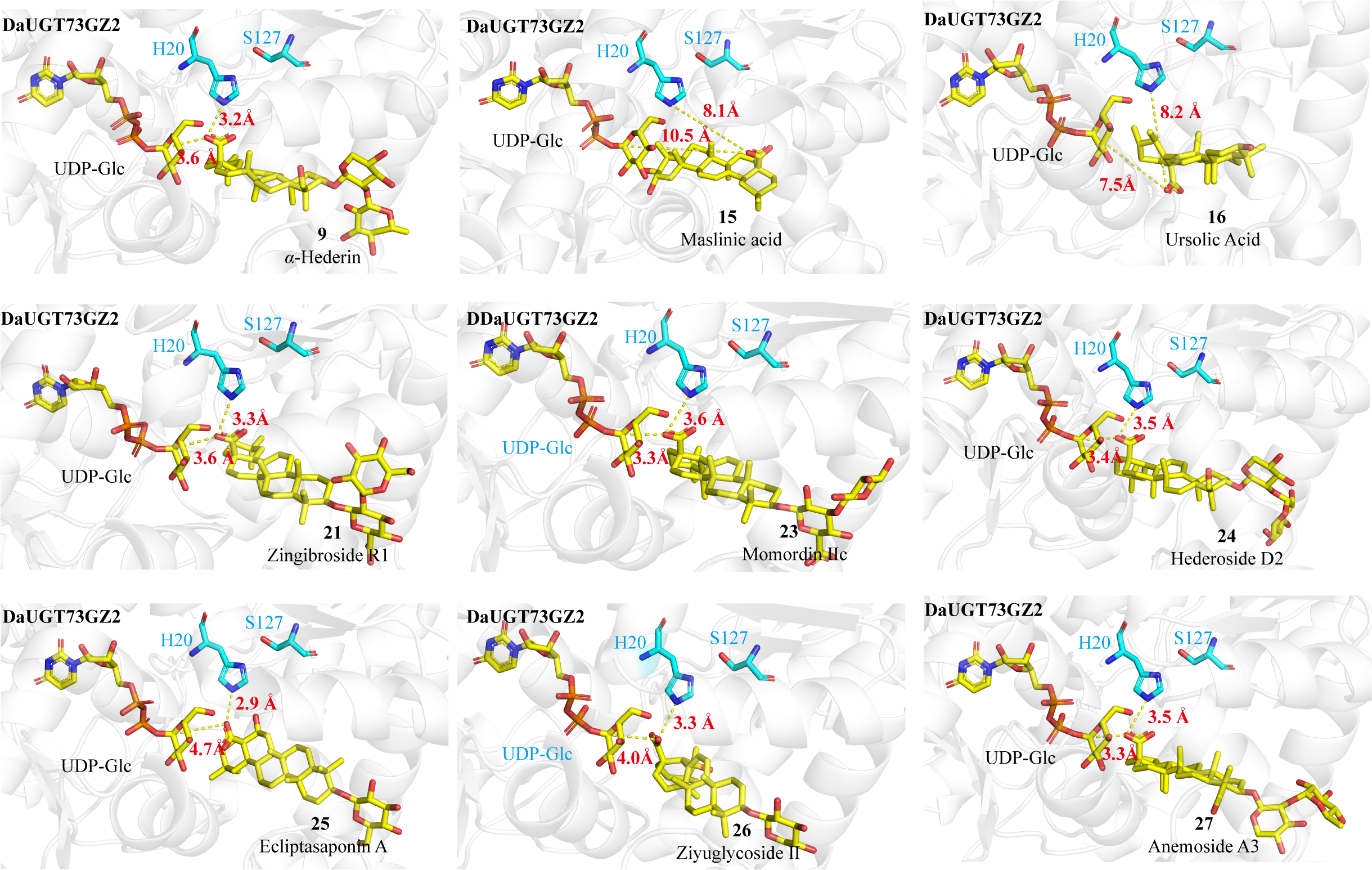

**Supplemental Figure 46. Predicted structure of protein DaUGT73GZ2 and its molecular docking models with ligands 9, 15, 16, 21, and 23-27**. Yellow dashed lines represent the distances between the *C*-28 carboxyl oxygen atom and the anomeric carbon of UDP-Glc, and between the *C*-28 carboxyl oxygen atom and the nitrogen atom of catalytic histidine H20, respectively.

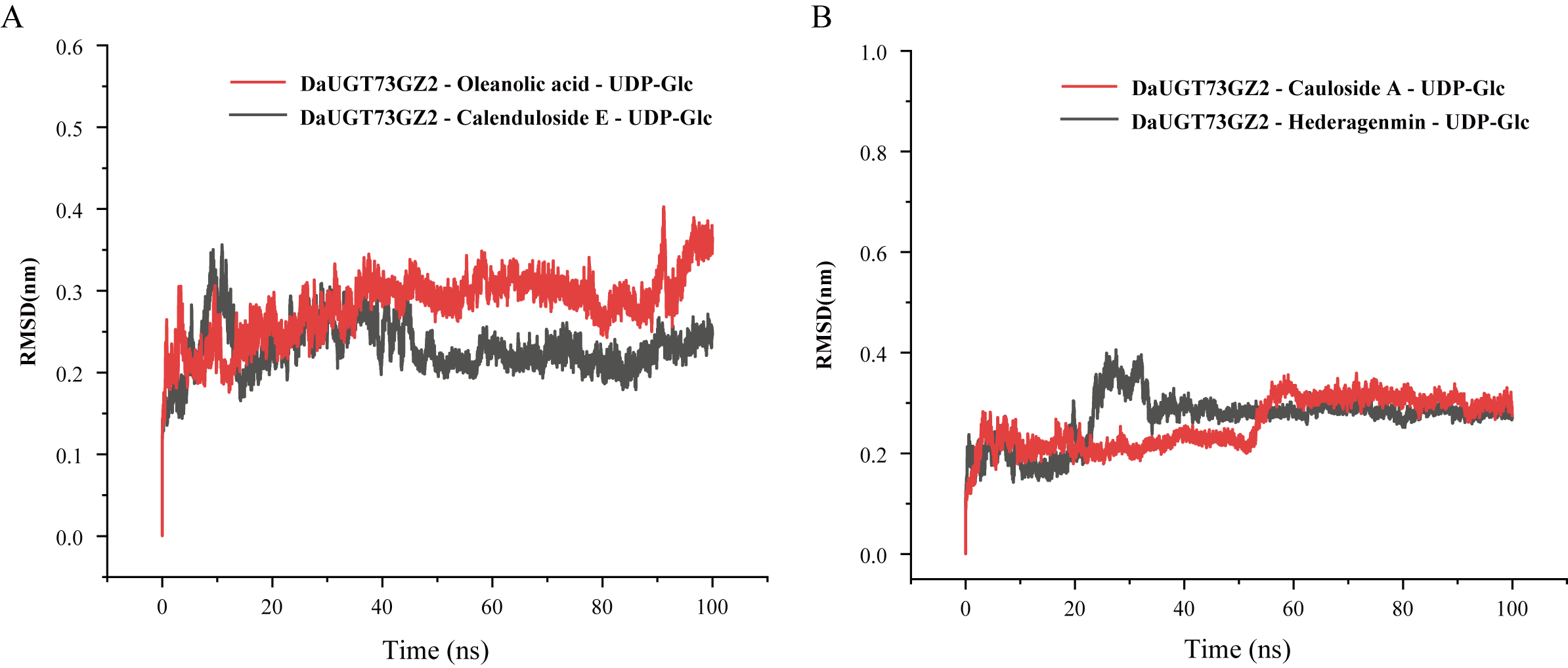

**Supplemental Figure 47. RMSD profiles of DaUGT73GZ2–substrate–UDP-Glc complexes during 100 ns molecular dynamics simulations. (A)** RMSD profiles of DaUGT73GZ2 in complex with oleanolic acid or calenduloside E and UDP-Glc. **(B)** RMSD profiles of DaUGT73GZ2 in complex with cauloside A or hederagenin and UDP-Glc.

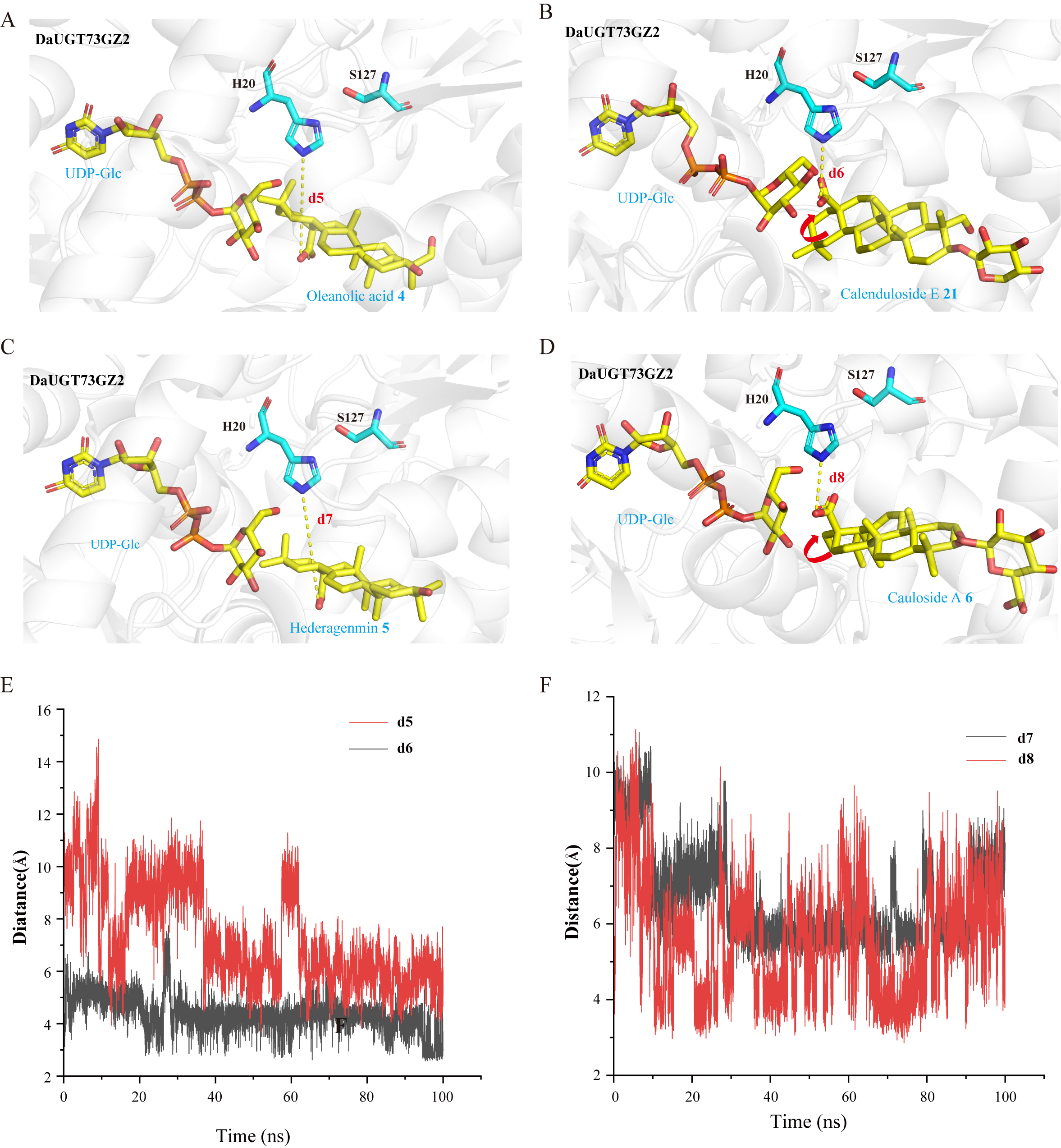

**Supplemental Figure 48. Distance analysis between the *C*-28 carboxyl oxygen atom of acceptor substrates and catalytic histidine H20 in DaUGT73GZ2–substrate–UDP-Glc complexes.**

**(A–D)** representative binding conformations of DaUGT73GZ2 with oleanolic acid, calenduloside E, hederagenin, or cauloside A in the presence of UDP-Glc. In each panel, the yellow dashed line indicates the distance between the *C*-28 carboxyl oxygen atom of the acceptor substrate and the nitrogen atom of catalytic histidine H20. **(E–F)** distance changes of d5–d8 during 100 ns molecular dynamics simulations.

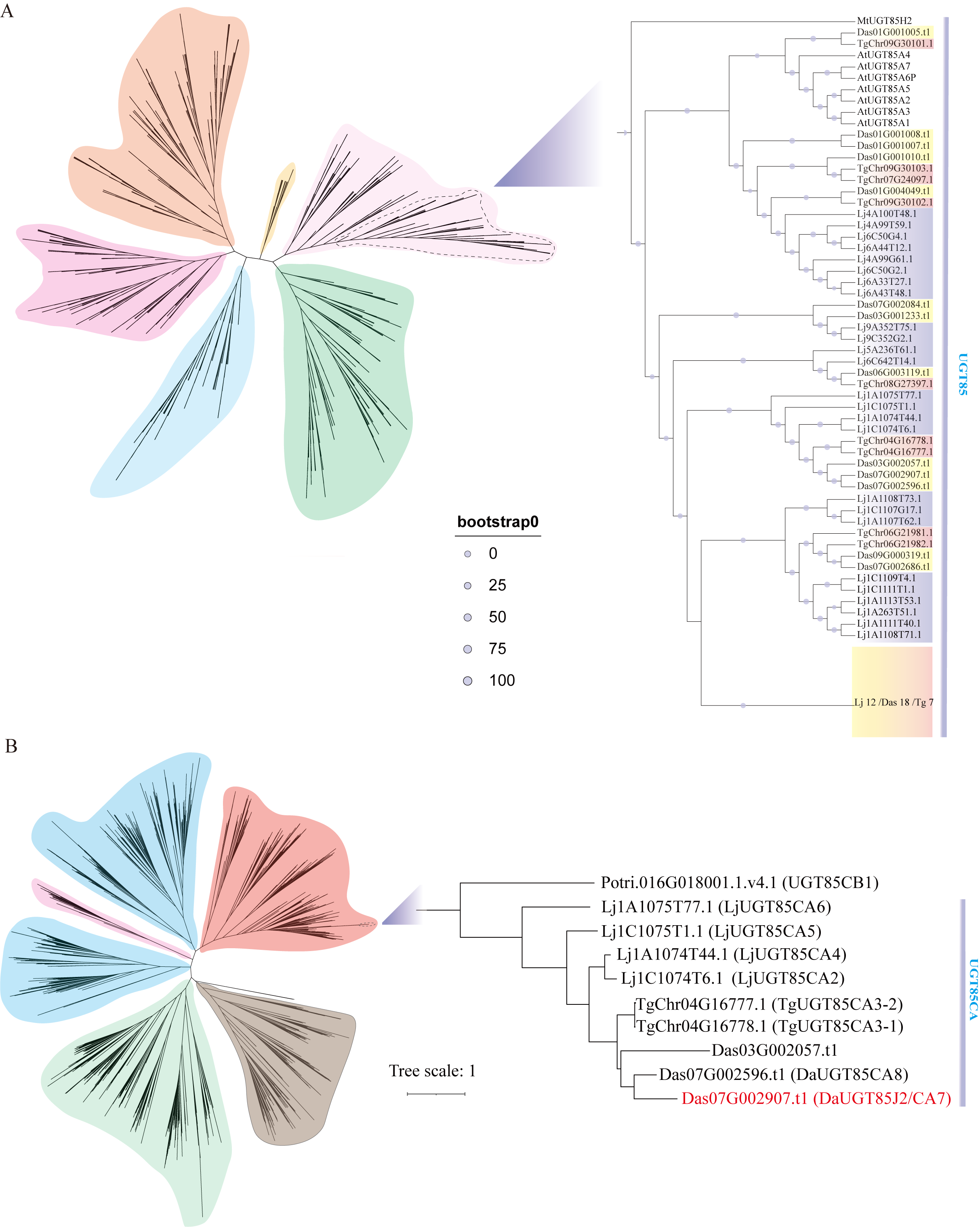

**Supplemental Figure 49.** **Phylogenetic analysis of the UGT gene family.** **(A)** Phylogenetic tree of UGTs from three species. The detailed topology of the UGT85 family is shown in the right panel. **(B)** Phylogenetic tree of UGTs constructed from 16 species, highlighting the evolutionary relationships within the UGT85CA clade including DaUGT85J2. The phylogenetic tree was constructed using the maximum likelihood method, in which tree nodes (>50%) are indicated by purple dots on branches (1000 bootstrap replications).

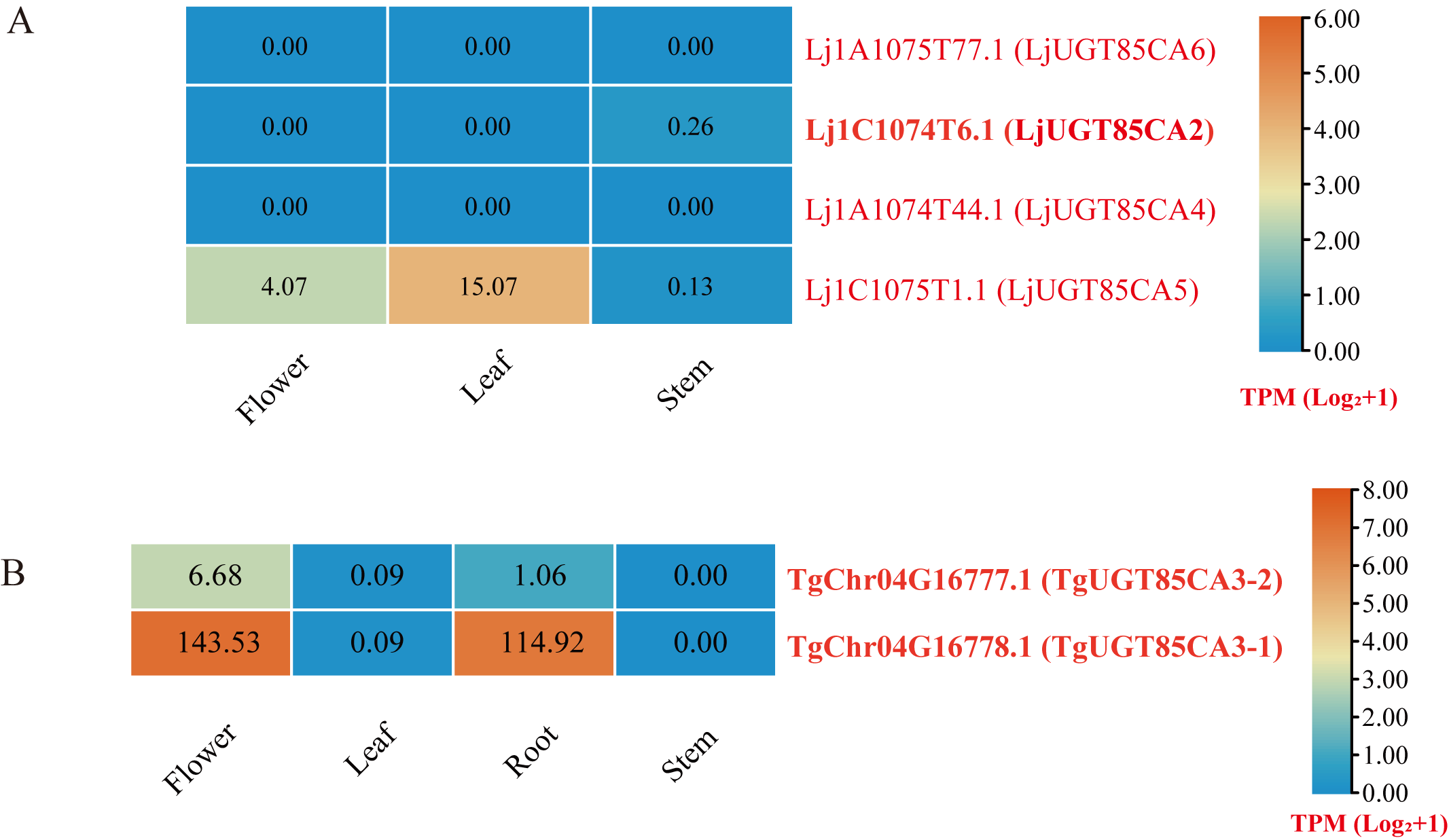

**Supplemental Figure 50. Expression levels of UGT85CA subfamily genes in *T. glandulifera*, and *L. japonica*.** **(A)** Expression levels of UGT85CA subfamily genes in *L. japonica*. **(B)** Expression levels of UGT85CA subfamily genes in *T. glandulifera*.

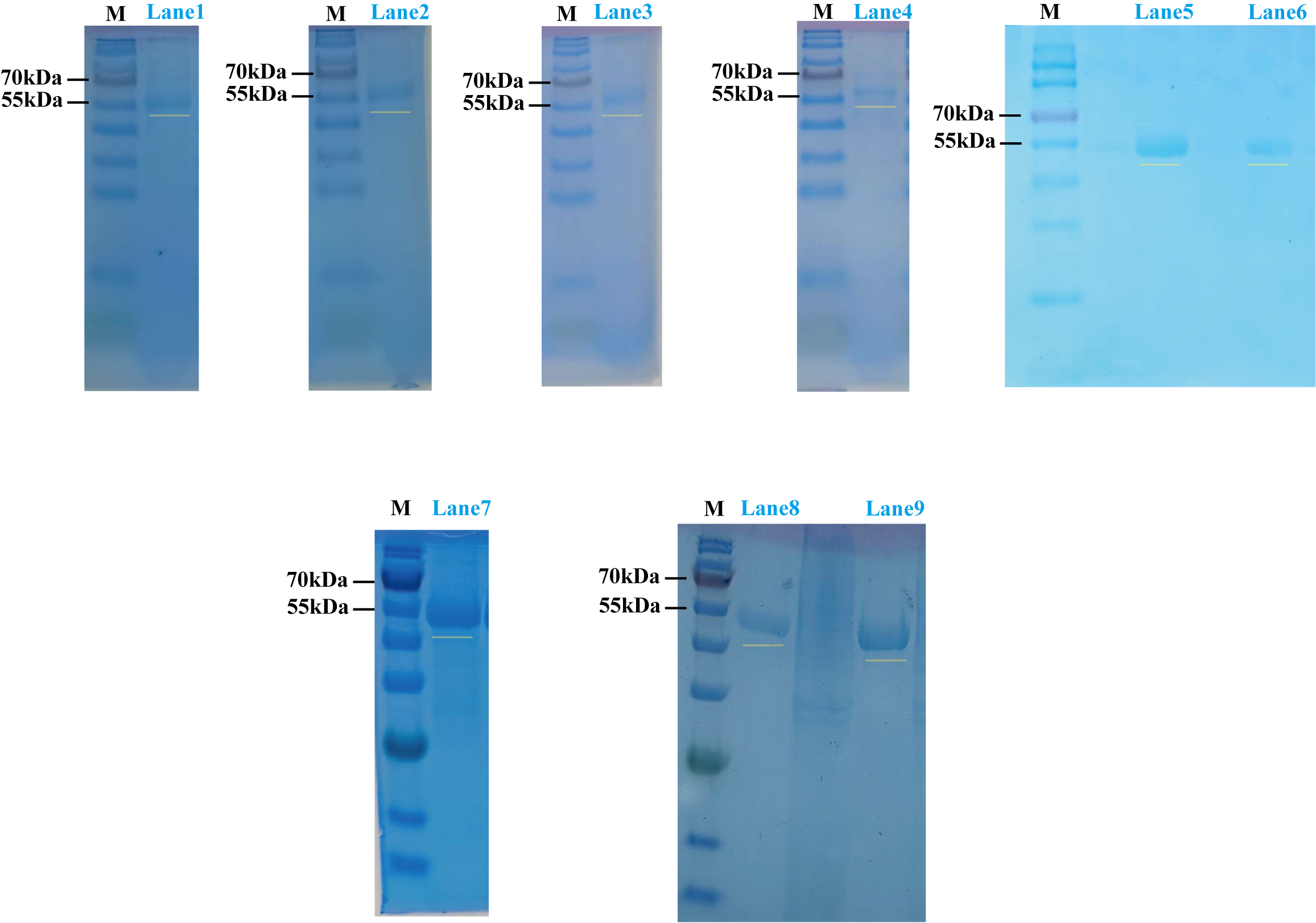

**Supplemental Figure 51. SDS-PAGE analysis of recombinant UGT proteins heterologously expressed in *E. coli*.** Lane M: protein molecular weight marker. Lanes 1–9 correspond to the purified candidate UGT proteins: TgUGT85CA3-1, TgUGT85CA3-2, LjUGT85CA2, DaUGT85A245, DaUGT85J2, DaUGT2596, LjUGT85CA5, anc156, and anc160. Yellow bars indicate the target protein bands.

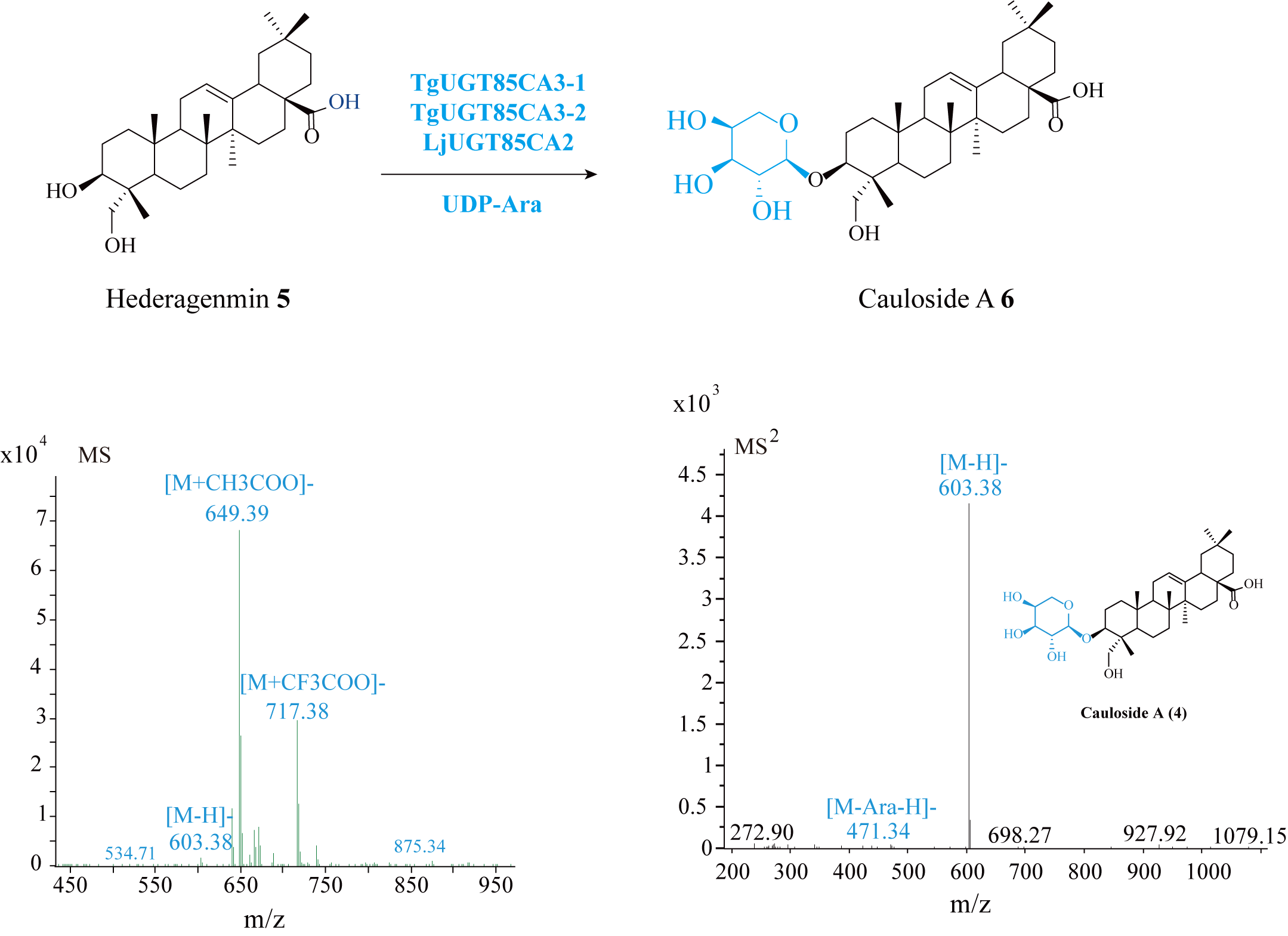

**Supplemental Figure 52. The (-)-ESI-MS and MS/MS spectra of 6.**

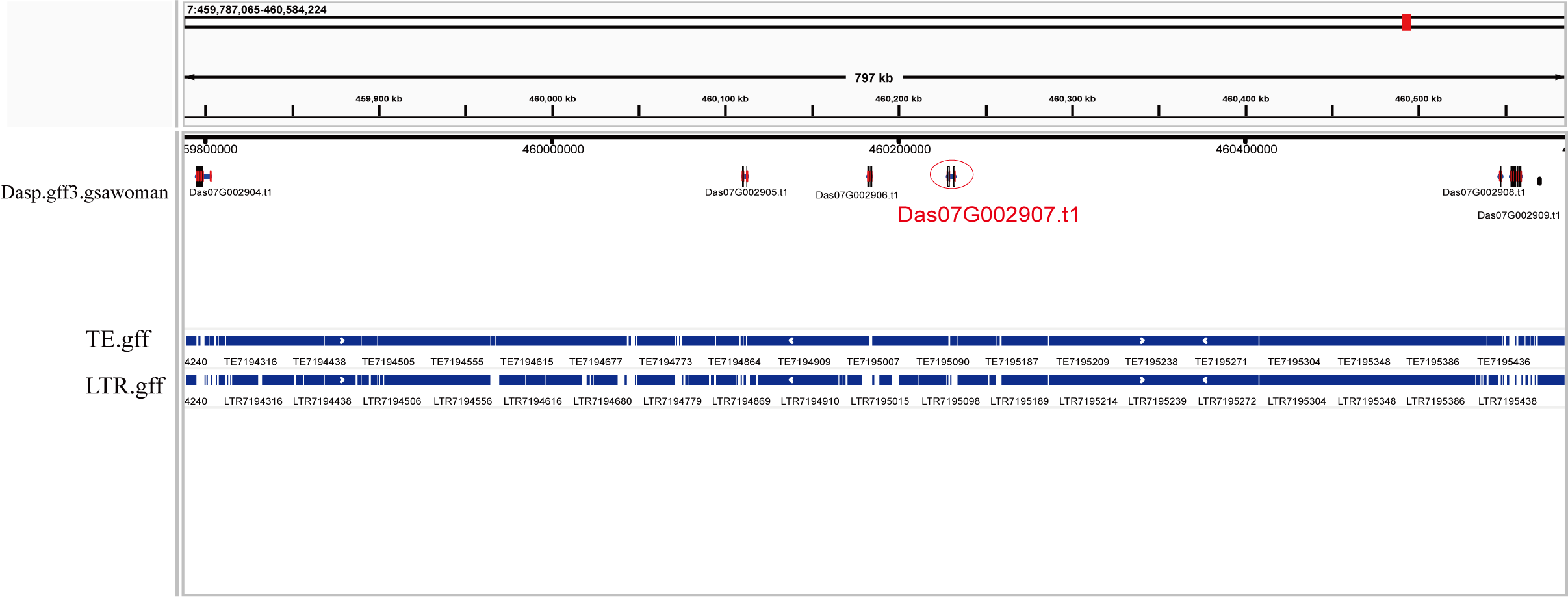

**Supplemental Figure 53. Quantification of TEs and LTRs within various genomic windows flanking *DaUGT85J2***.

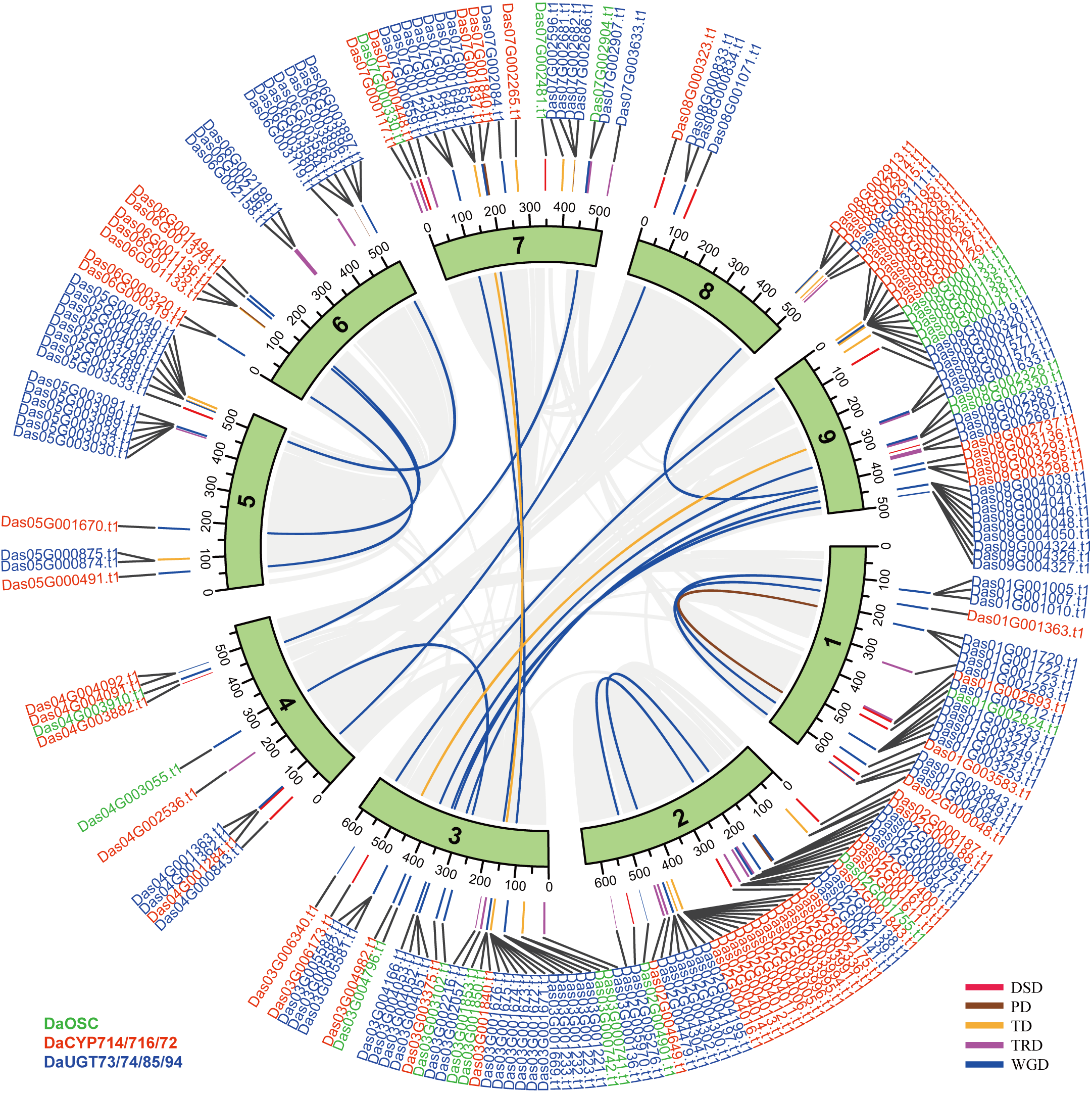

**Supplemental Figure 54.** **Gene positions and duplication sources of related genes in *D. asperoides***.

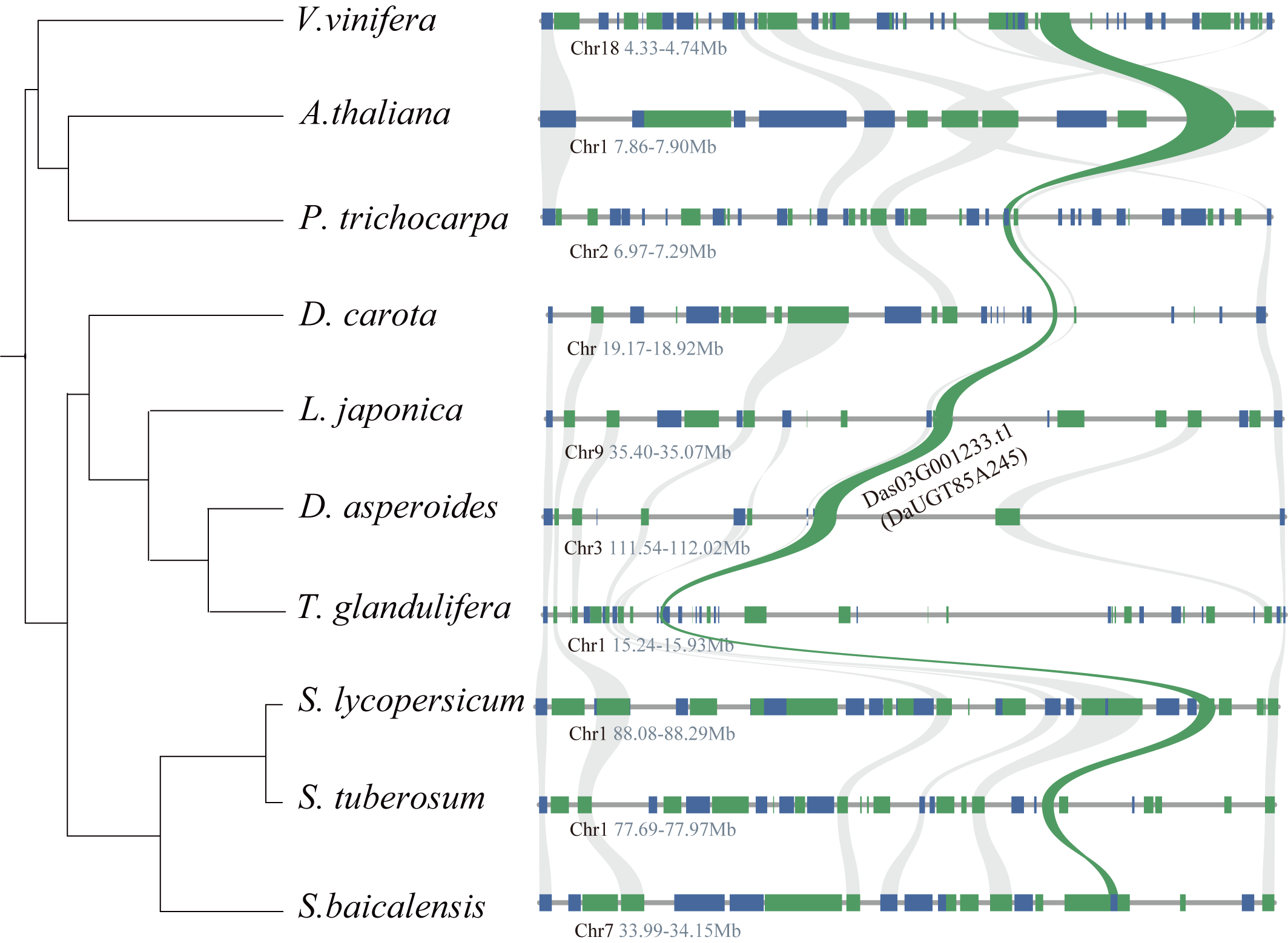

**Supplemental Figure 55.** **Collinearity analysis of Da03G001233.t1 (DaUGT85A245)**.

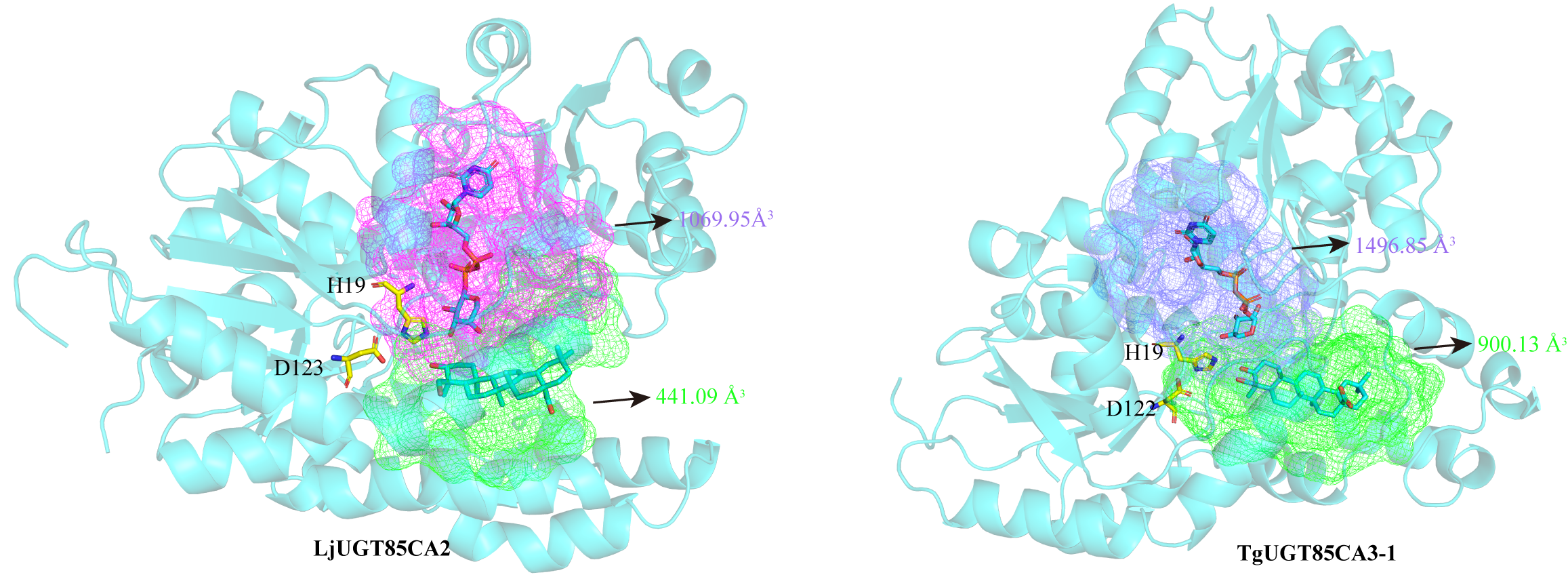

**Supplemental Figure 56. Predicted donor- and acceptor-binding pockets of LjUGT85CA2 and TgUGT85CA3-1**.

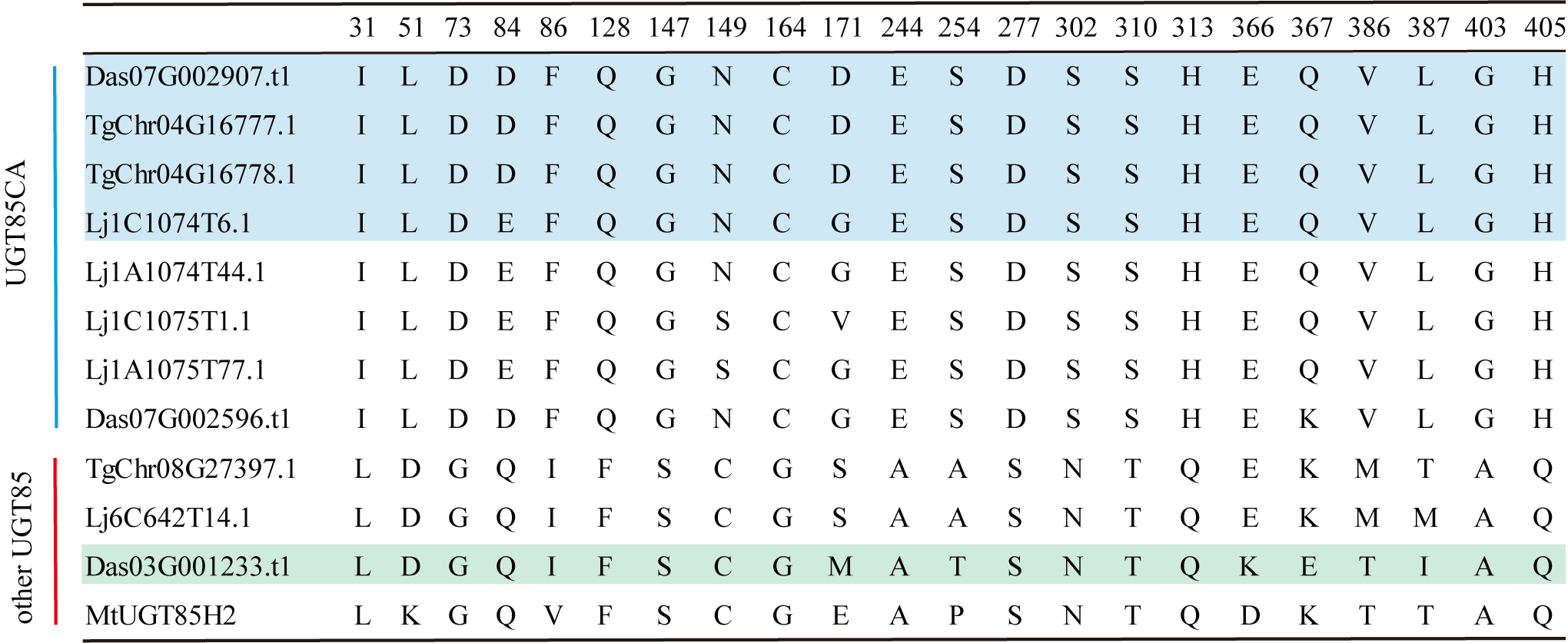

**Supplemental Figure 57. Sequence alignment of UGT85CA arabinosyltransferases and related UGT85 homologs**.

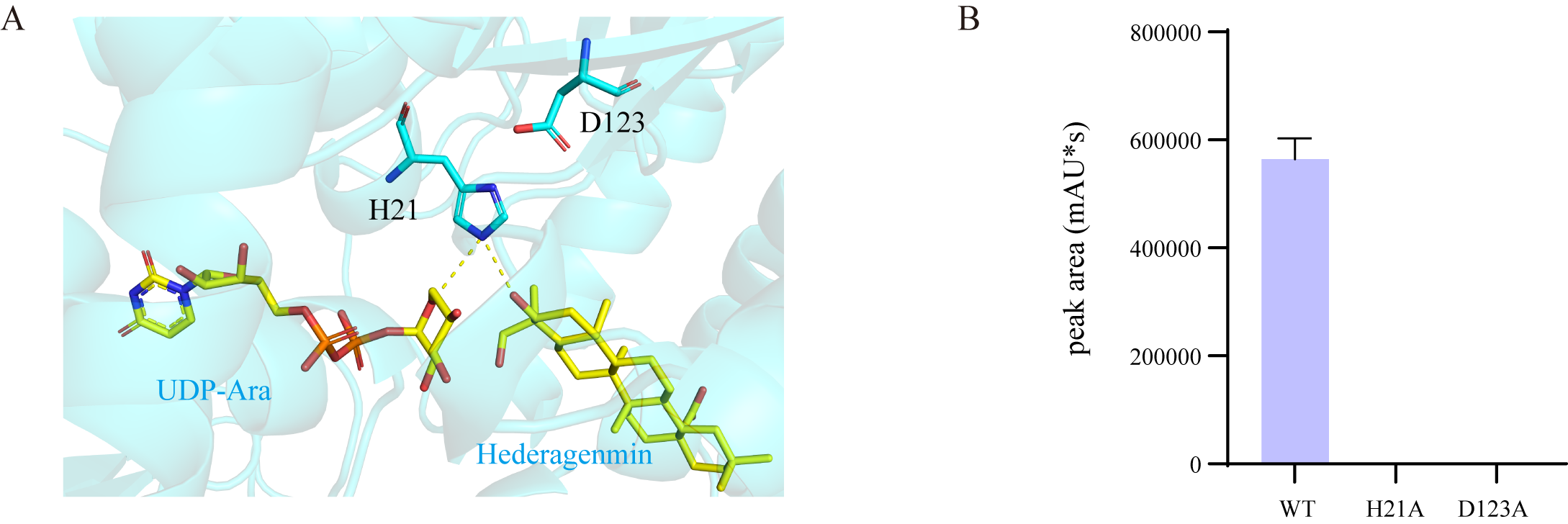

**Supplemental Figure 58. Analysis of catalytic mechanism of DaUGT85J2**. **(A)** Schematic diagram of catalytic mechanism of DaUGT85J2**.** Hydrogen bonds are indicated by yellow dashed lines. **(B)** Catalytic activities of the wild-type enzyme (WT) and the H21A and D123A mutants, as determined by the chromatographic peak area of the reaction product. Data are presented as the mean ± SD of three independent experiments.

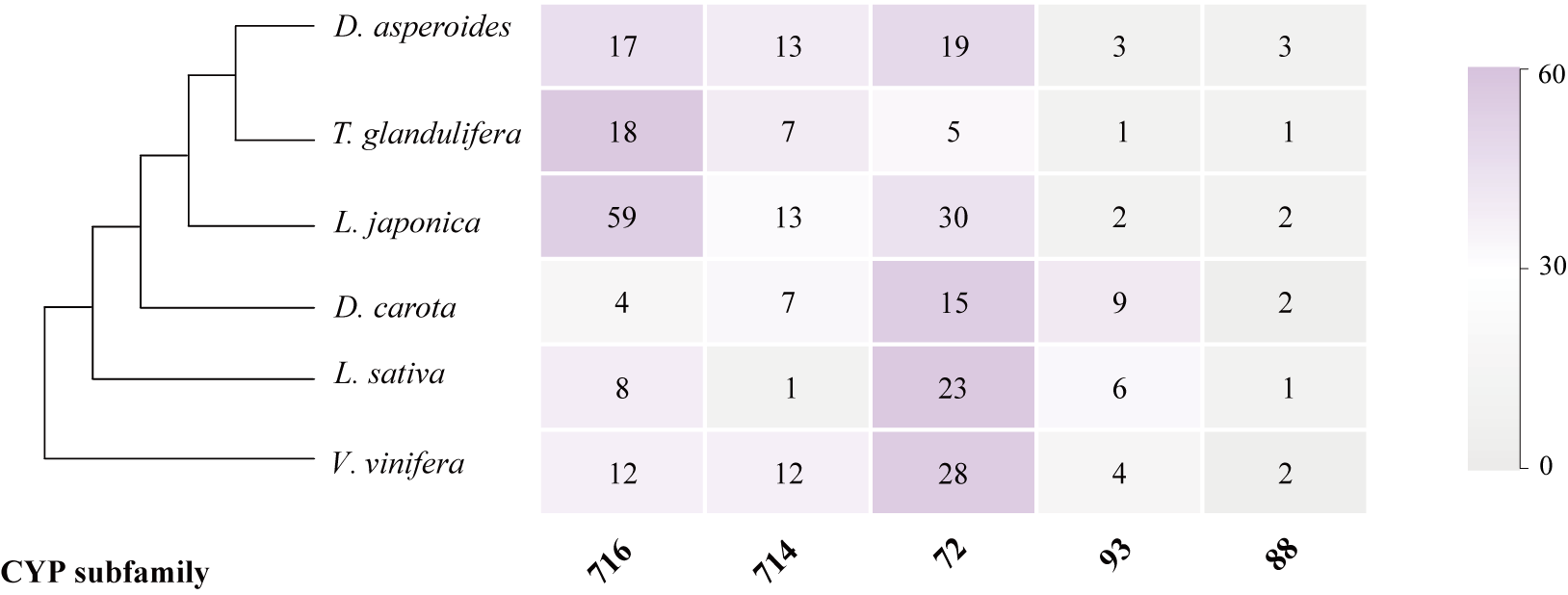

**Supplemental Figure 59. The number of members in five CYP families of *D. asperoides* and five other species**.

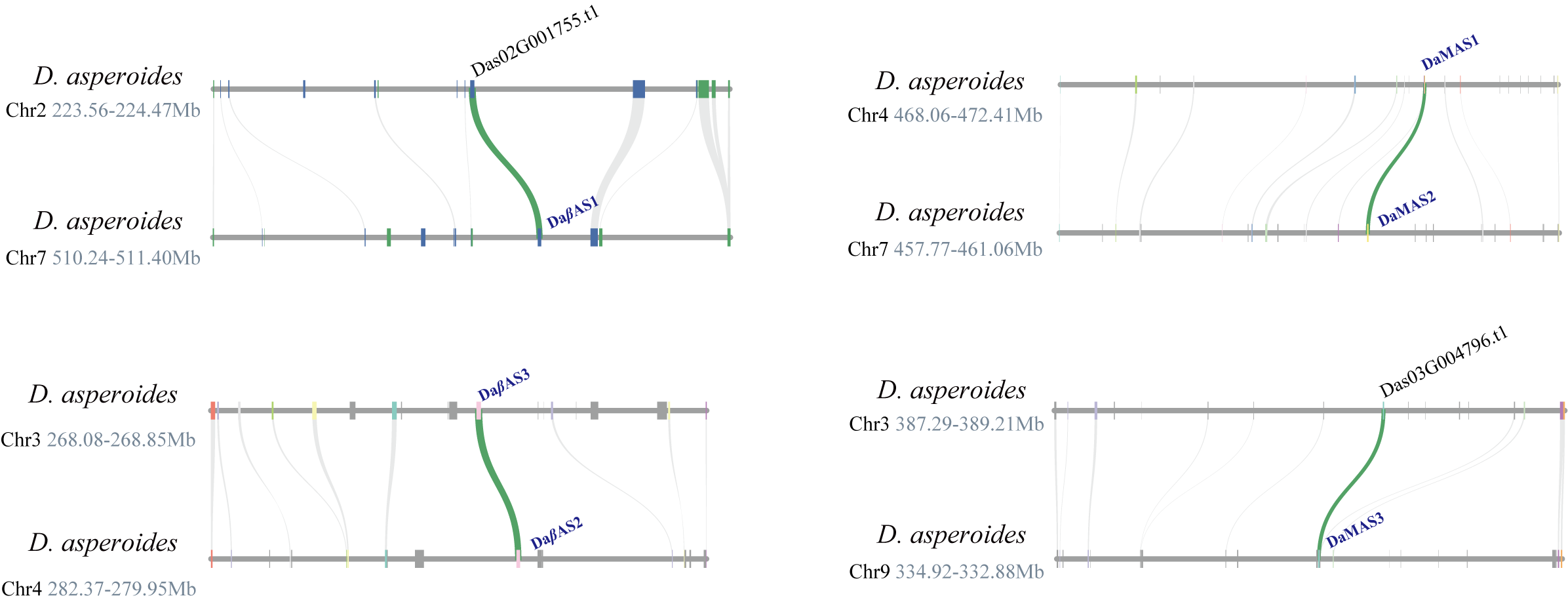

**Supplemental Figure 60. Intraspecific micro-synteny relations of OSC genes. Direct collinear relations for OSC genes were highlighted in green**.

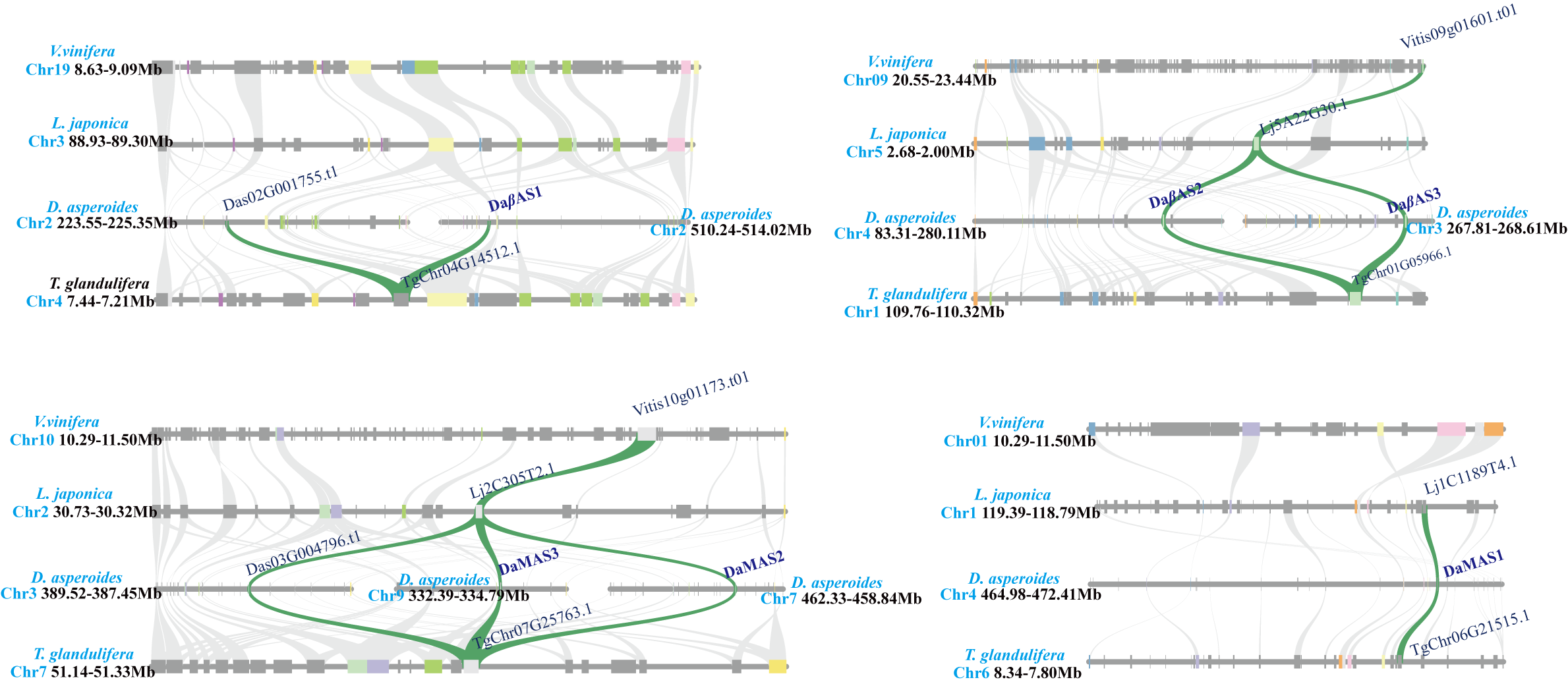

**Supplemental Figure 61. Collinearity analysis of *β*AS and MAS**.

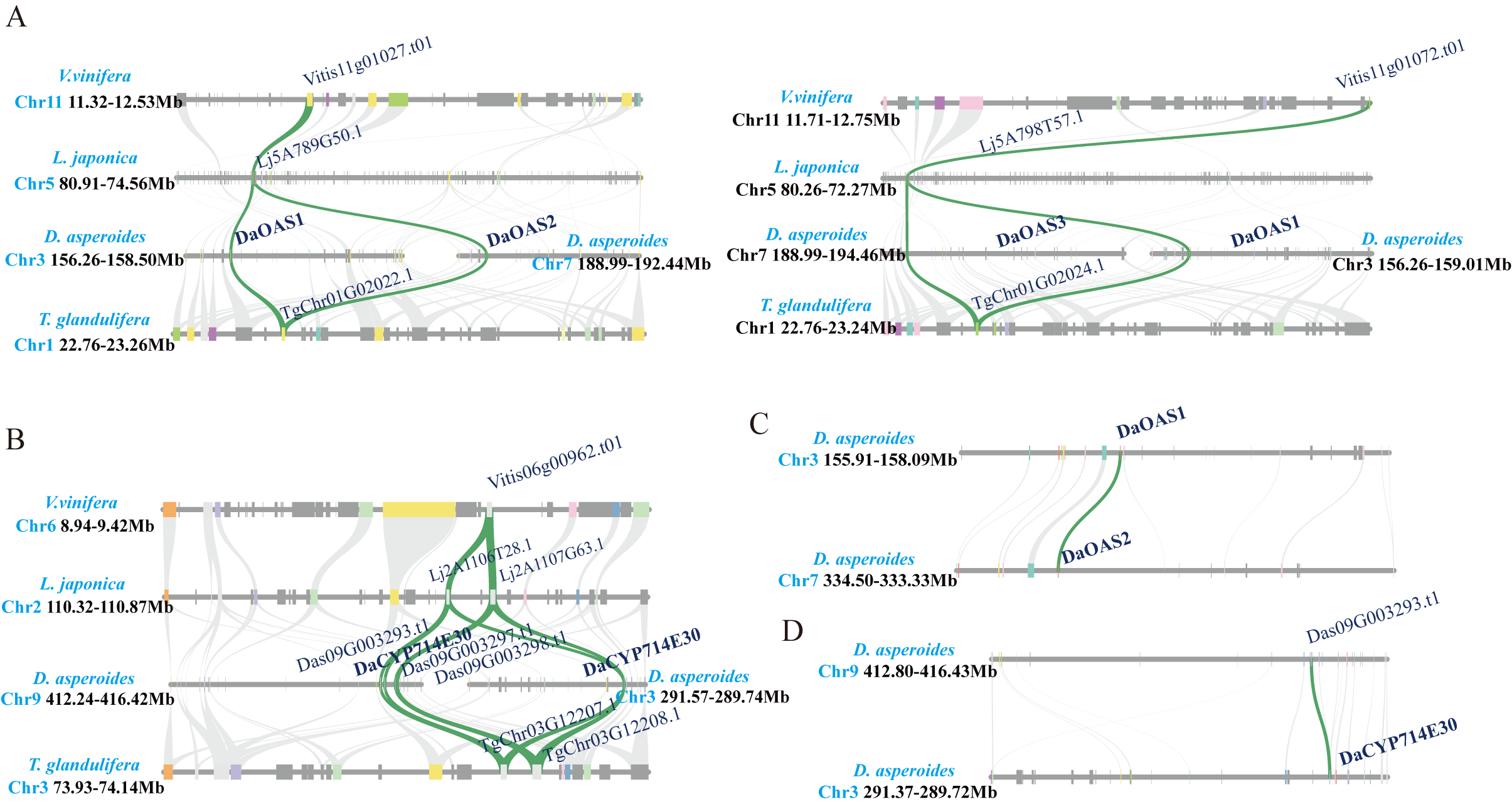

**Supplemental Figure 62.** **Analysis of micro-syntenic relations of functional DaCYPs genes. (A)** and **(B)** Interspecific synteny analysis relations of functional DaCYPs genes. (**C** – **D)** Intraspecific micro-synteny relations of functional DaCYPs genes.

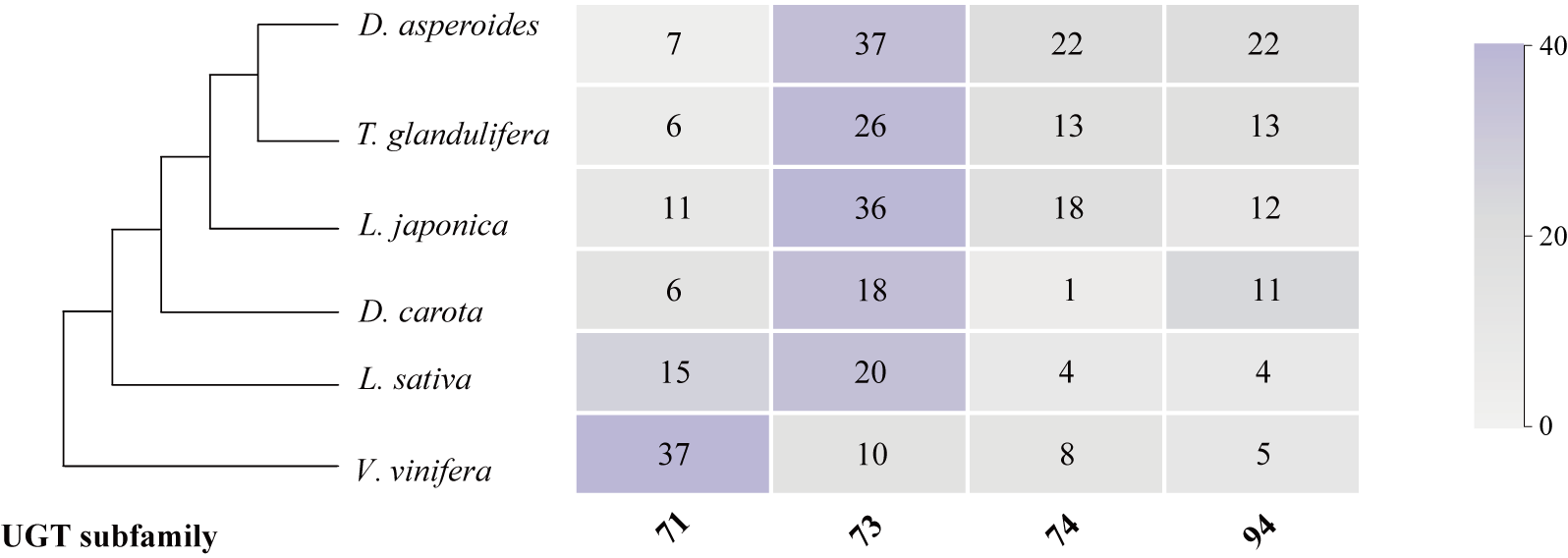

**Supplemental Figure 63.** **The number of members in UGT71/73/74/94 families of *D. asperoides* and five other species**.

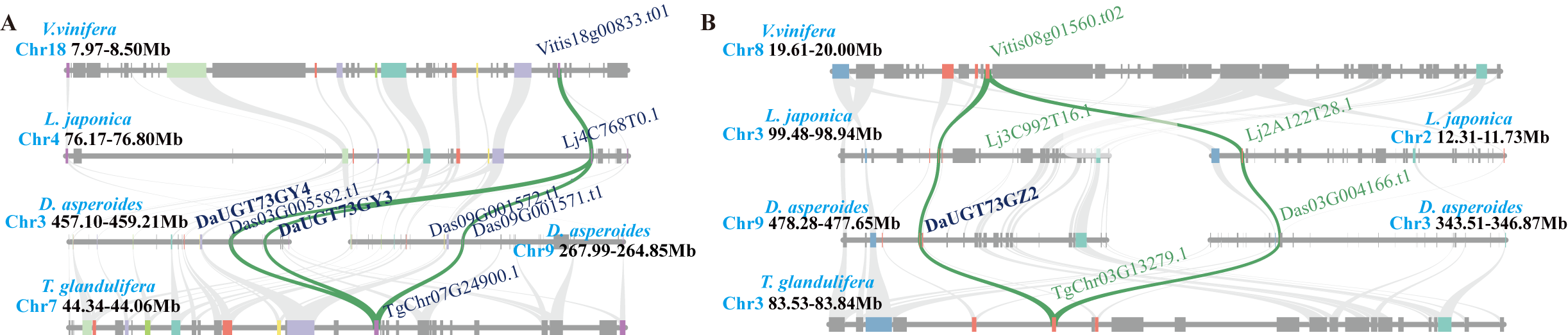

**Supplemental Figure 64. Syntenic relationships of functional UGT73 genes involved in hederagenin-based saponin biosynthesis**.

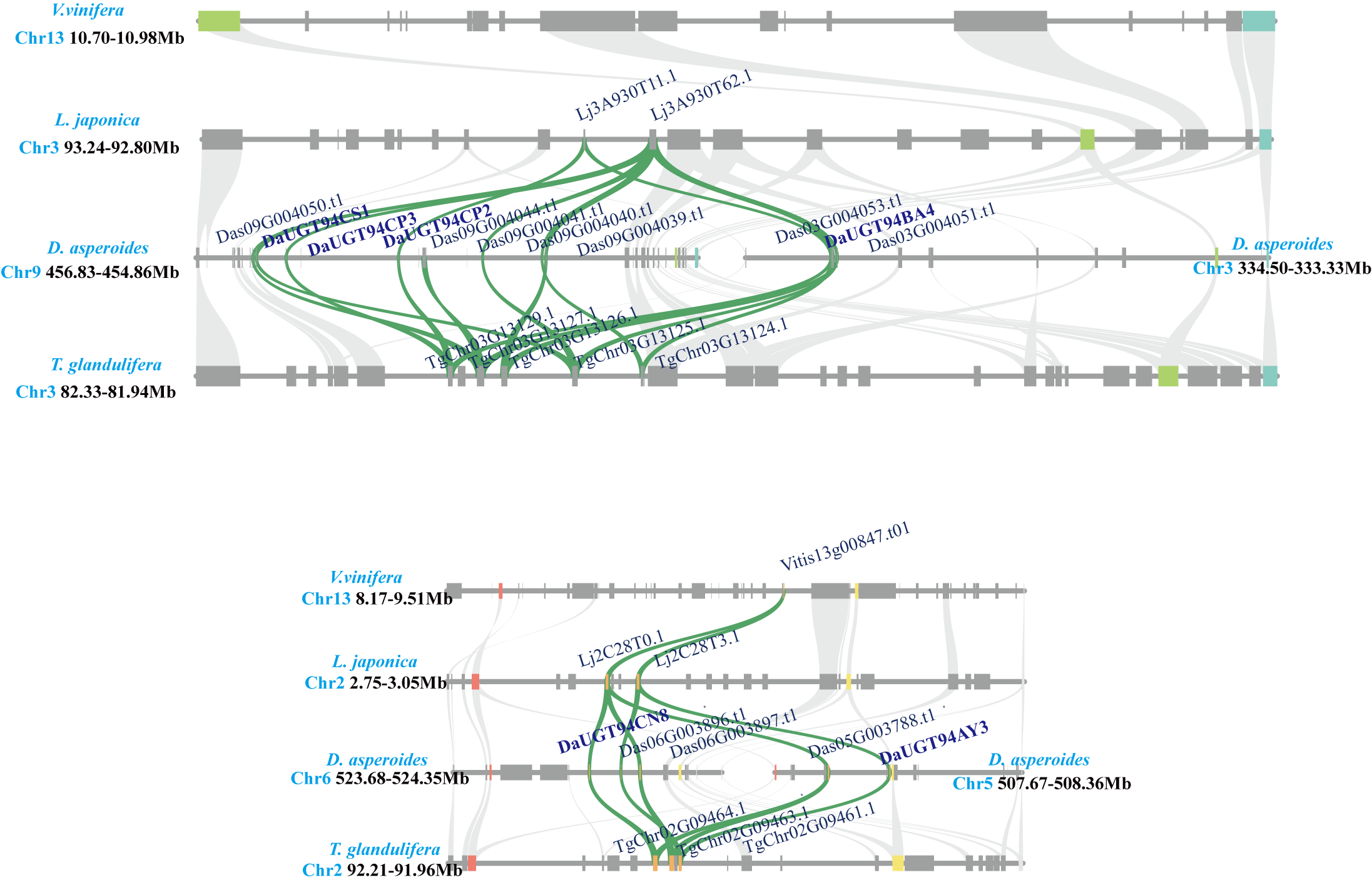

**Supplemental Figure 65. Syntenic relationships of functional UGT94 genes involved in hederagenin-type saponin biosynthesis**.

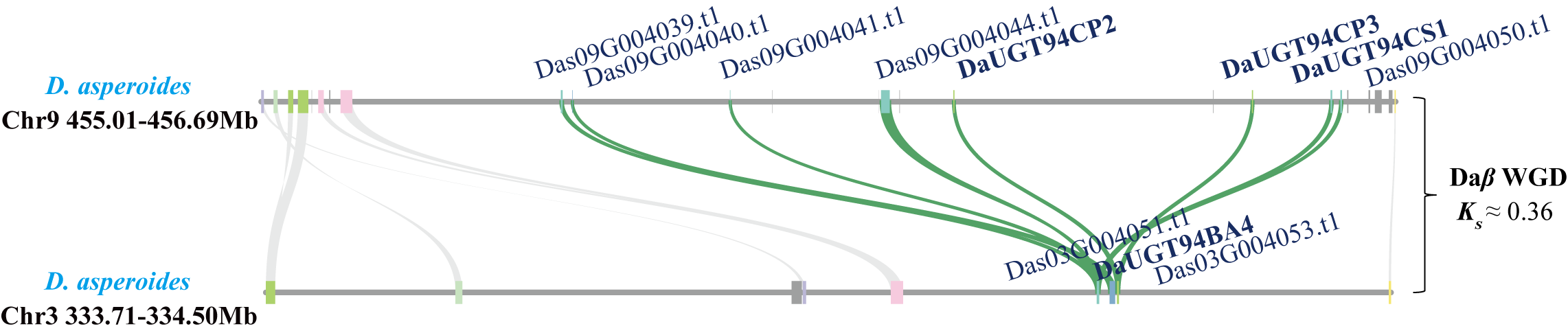

**Supplemental Figure 66. Synteny analysis of UGT94 family genes on chromosomes 3 and 9 in *D. asperoides***.

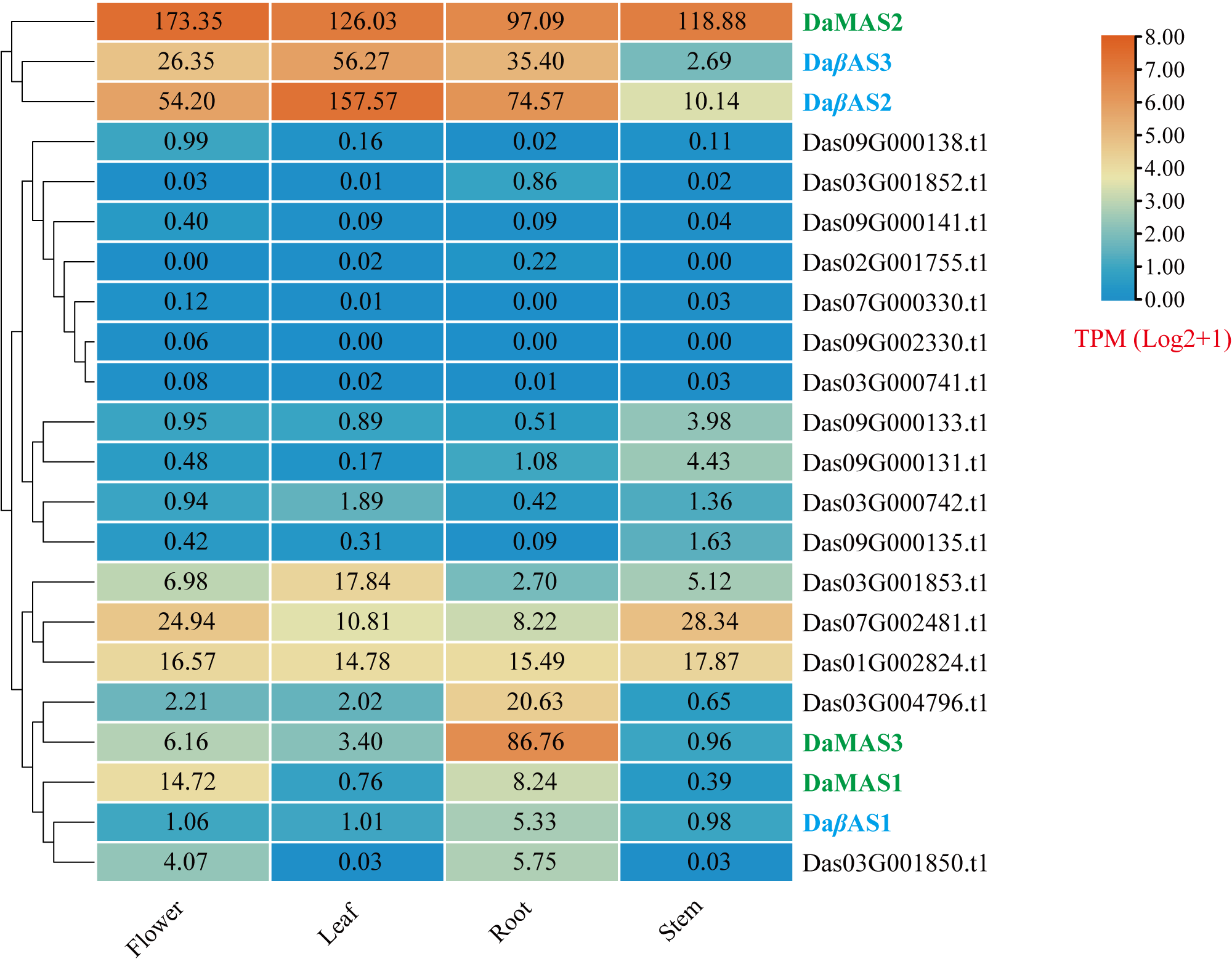

**Supplemental Figure 67.** **Heatmap of the OSC gene family in different tissues of *D. asperoides****.* Cyan and red represent the OSC genes of the *β*AS and MAS clades, respectively.

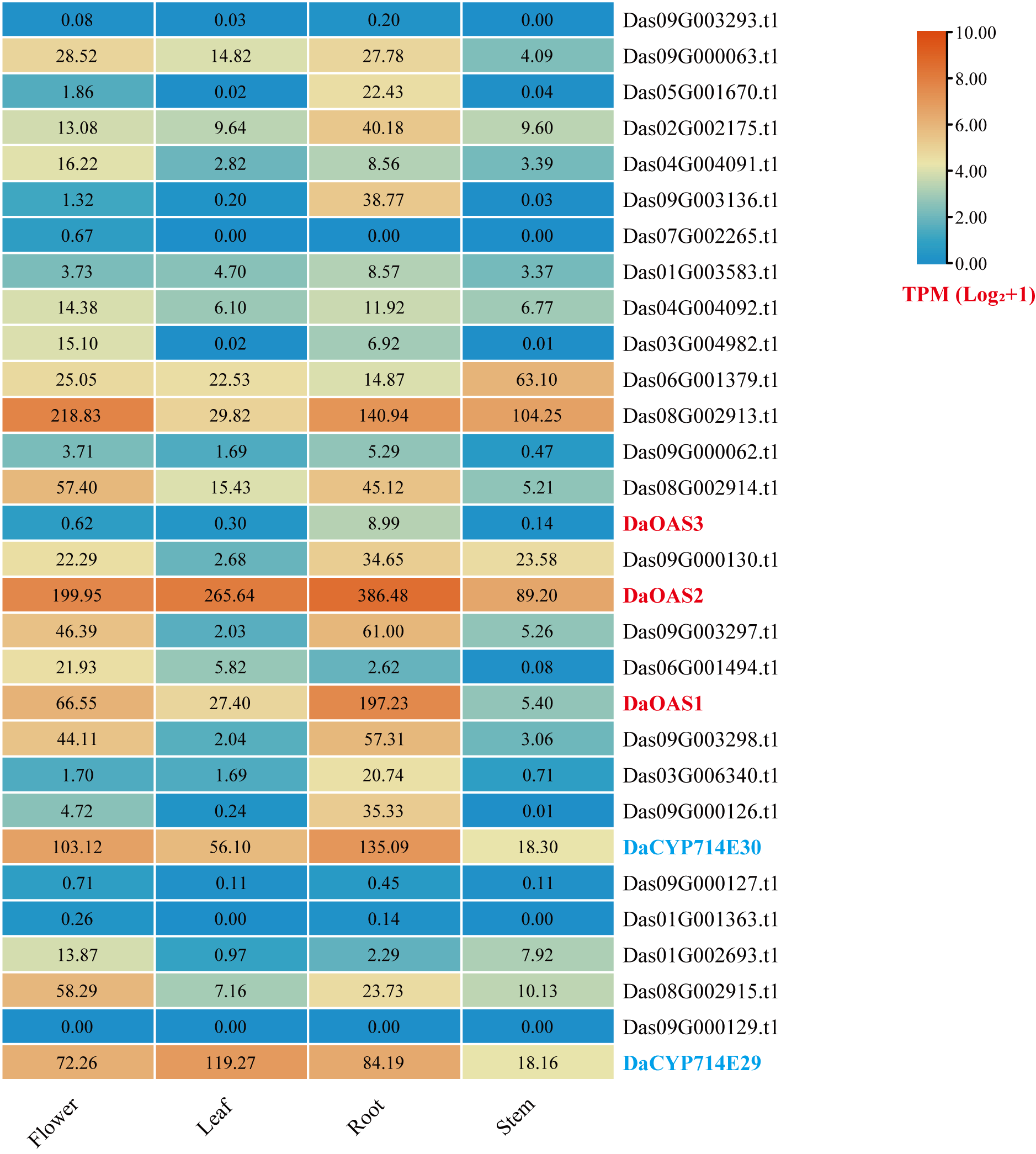

**Supplemental Figure 68.** **Heatmap of the CYP716/714 gene family in different tissues of *D. asperoides***.

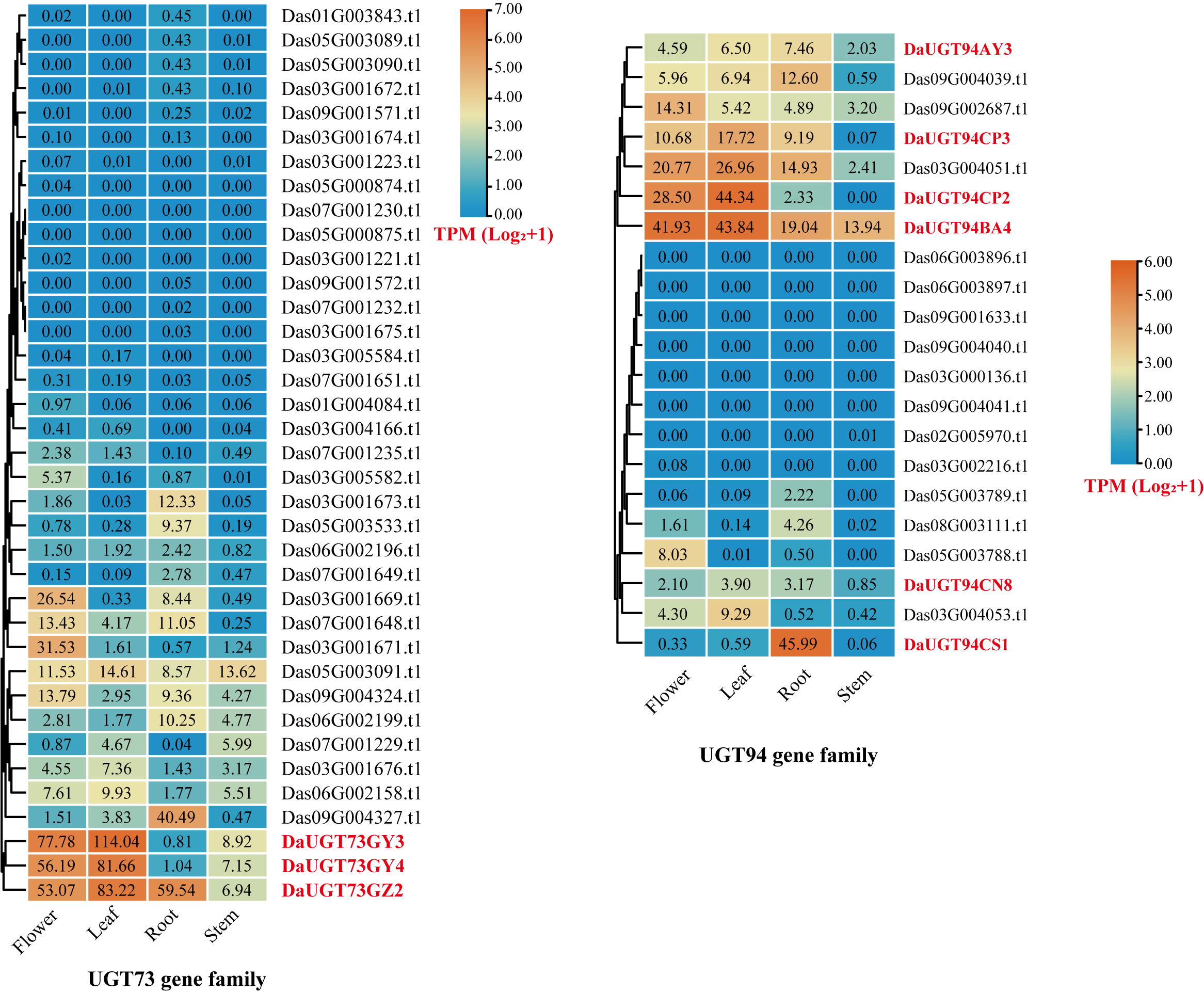

**Supplemental Figure 69.** **Heatmap of the UGT 73/94 gene family in different tissues of *D. asperoides***.

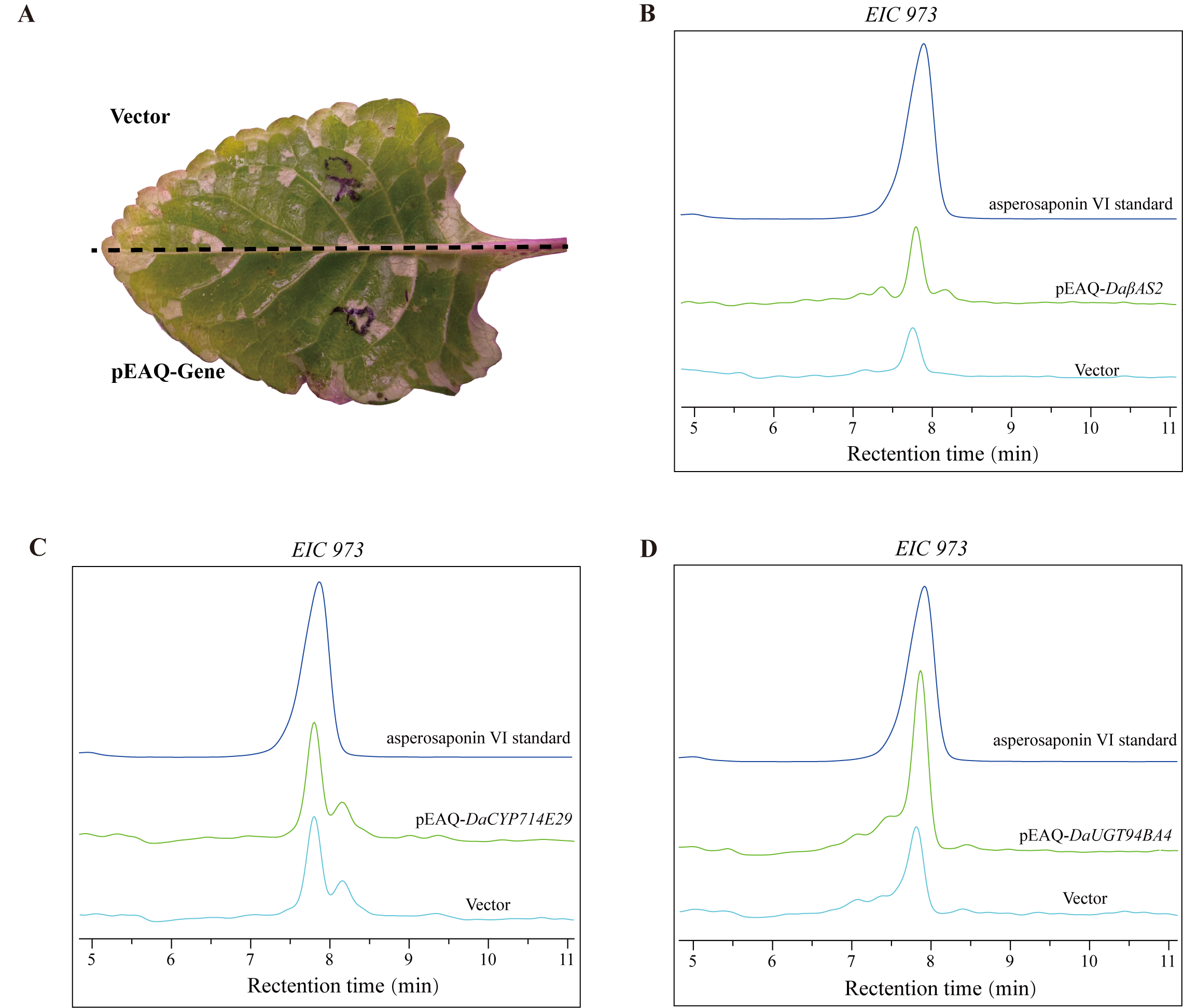

**Supplemental Figure 70. LC-MS analysis for the molecular function of key pathway genes by Agrobacterium-mediated transient expression in *D. asperoides* leaves**. **(A)**, schematic representation of the transient expression assay. One half of each leaf was infiltrated with Agrobacterium carrying the empty vector, while the other half was infiltrated with Agrobacterium carrying the target gene. **(B–D)**, extracted ion chromatograms (EICs) at m/z 973 showing asperosaponin VI signals in leaves transiently overexpressing *DaβAS2*, *DaCYP714E29* or *DaUGT94BA4*, respectively, compared with the vector control and asperosaponin VI standard.

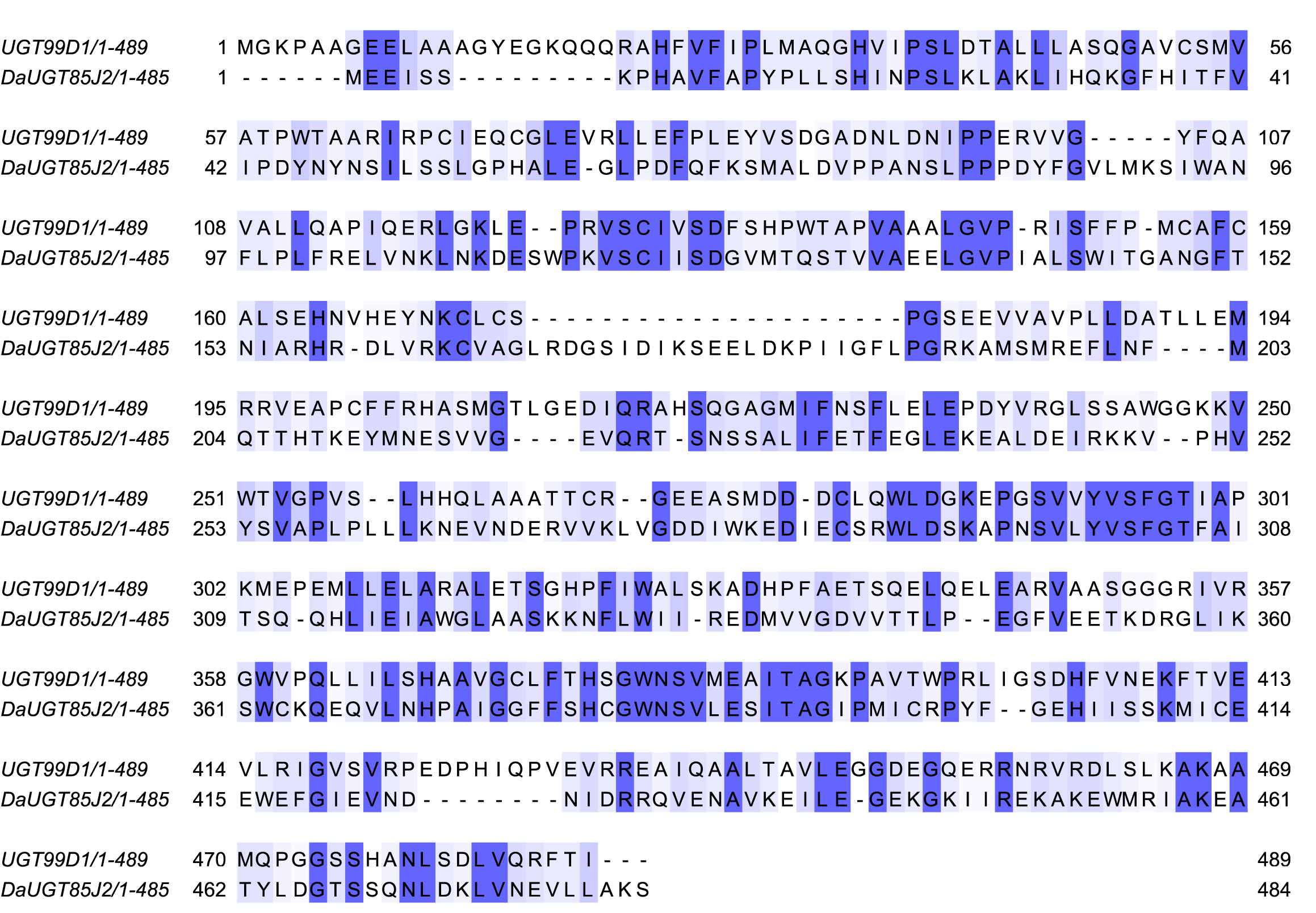

**Supplemental Figure 71. Sequence alignment of DaUGT85J2 and UGT99D1**.

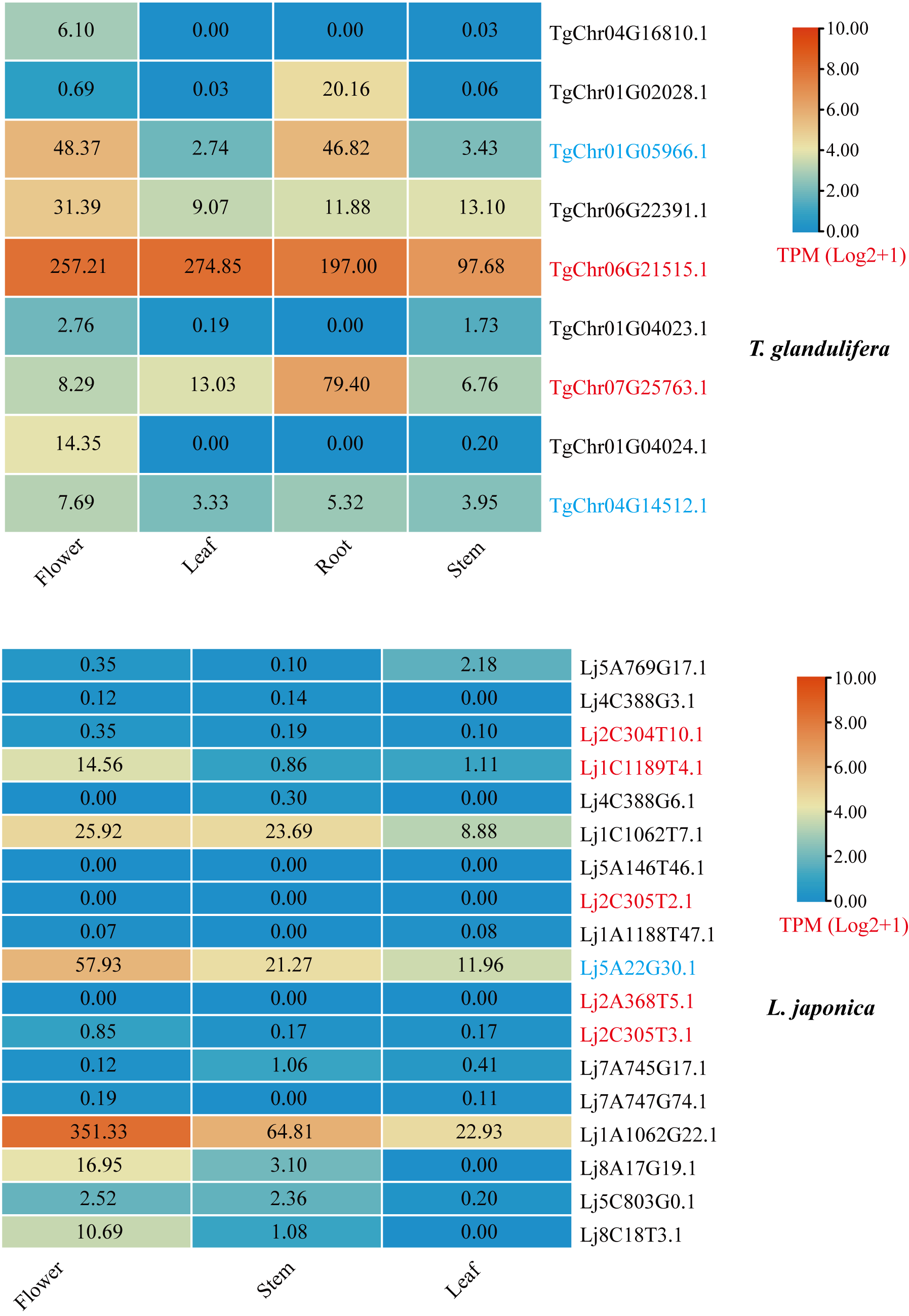

**Supplemental Figure 72. Heatmap of the OSC gene family in different tissues of *L. japonica* and *T. glandulifera****.* Cyan and red represent the OSC genes of the *β*AS and MAS clades, respectively.

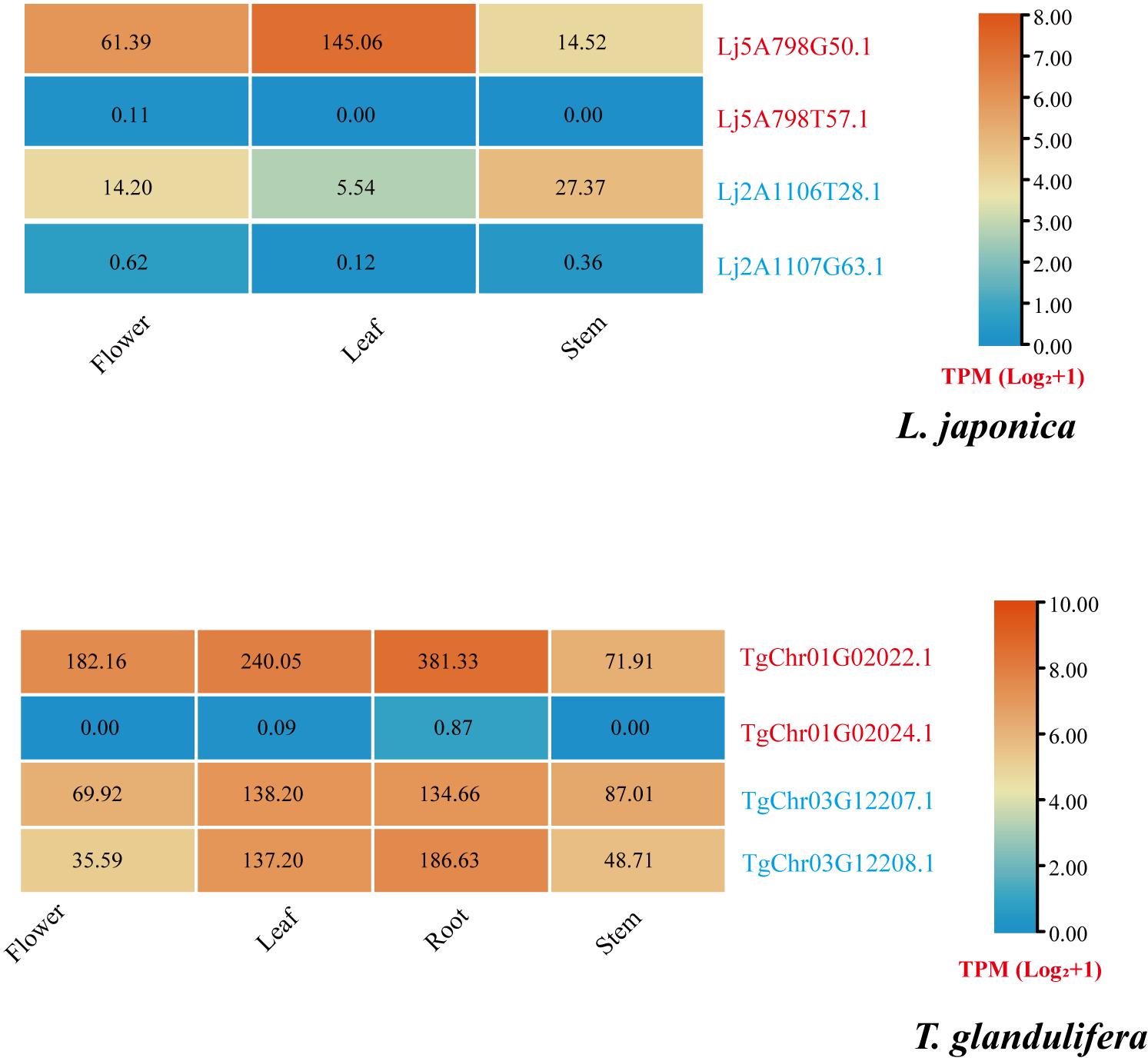

**Supplemental Figure 73. Expression profiles of CYP genes in *L. japonica* and *T. glandulifera* syntenic to functional *D. asperoides* CYPs**.

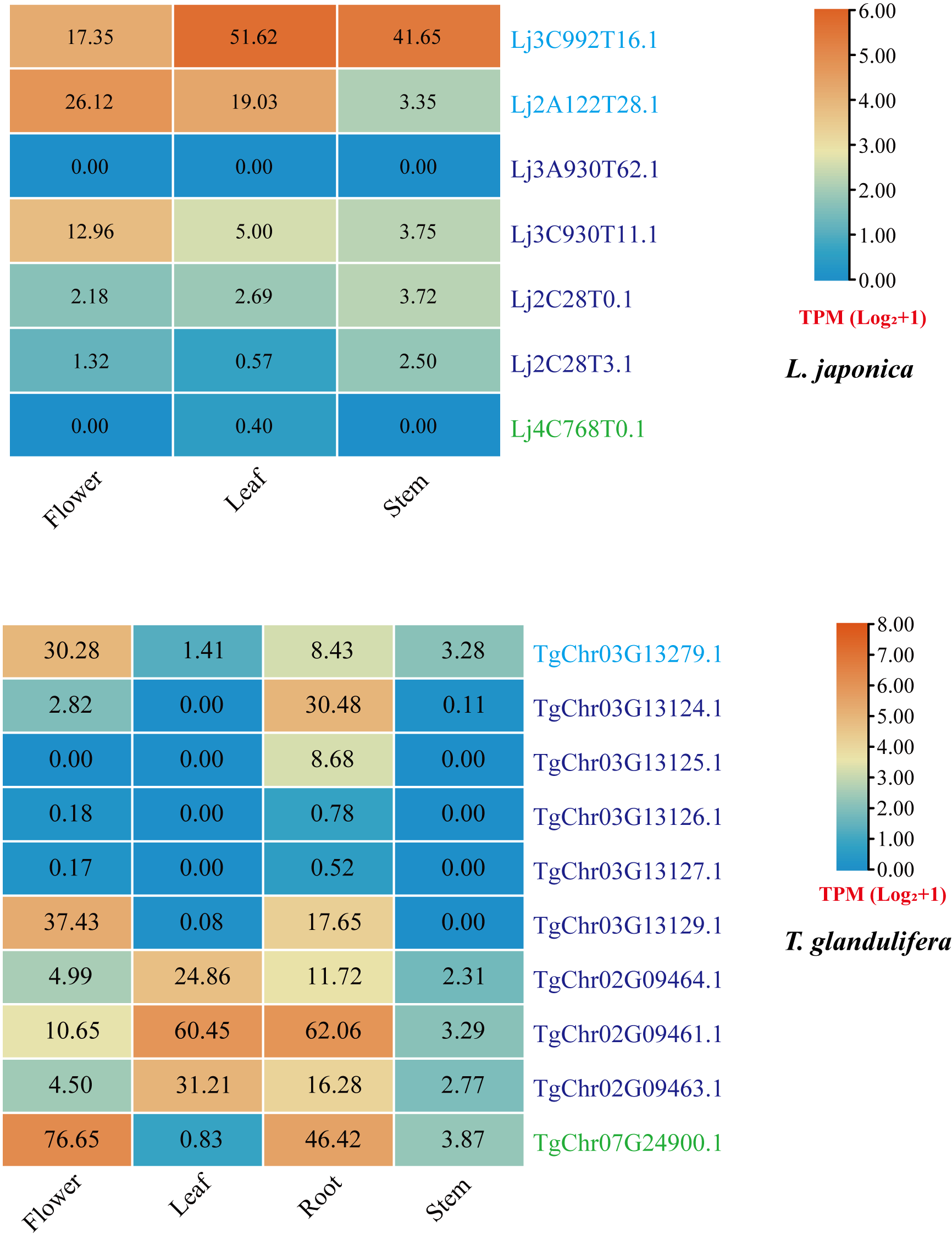

**Supplemental Figure 74. Expression profiles of UGT genes in *L. japonica* and *T. glandulifera* syntenic to functional *D. asperoides* UGTs**.

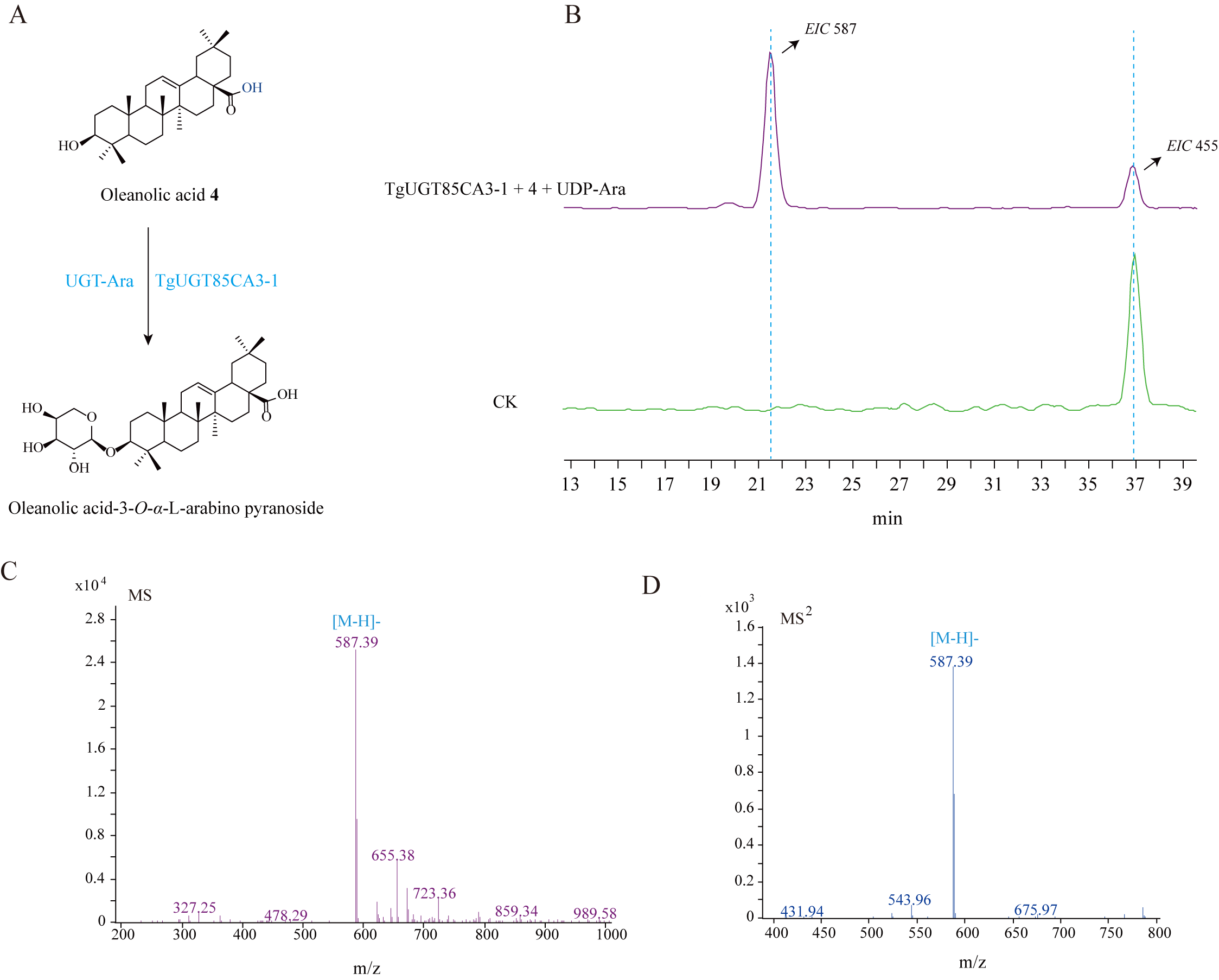

**Supplemental Figure 75**. **In vitro functional characterization of TgUGT85CA3-1 using oleanolic acid as substrate**. **(A)** Reaction scheme of oleanolic acid glycosylation. **(B)** LC-MS chromatograms of the enzymatic product and control. **(C)** Mass spectrum and **(D)** MS/MS fragmentation of the newly formed saponin.

**Supplementary Table 1. LC–MS analysis of triterpenoids and triterpenoid saponins in enzymatic reaction mixtures.**

| No. | CAS No. | Compound | Formula | Pseudomolecular  ion | Retention time  (min) | Standard *m/z* | Detected *m/z* in enzyme assays |
| --- | --- | --- | --- | --- | --- | --- | --- |
| **4** | 508-02-1 | Oleanolic acid | C₃₀H₄₈O₃ | [M-H]- | 36.87 | 455.35 | 455.35 |
| **5** | 465-99-6 | Hederagenin | C₃₀H₄₈O₄ | [M-H]- | 17.40 | 471.34 | 471.34 |
| **6** | 631-01-6 | Cauloside A | C_35_H_56_O_8_ | [M+HCOO]- | 13.30 | 649.39 | 649.39 |
| **7** | 39524-13-5 | HN saponin F | C_41_H_66_O_13_ | [M+HCOO]- | 9.53 | 811.45 | 811.45 |
| **8** | 39524-08-8 | Asperosaponin Ⅵ | C_47_H_76_O_18_ | [M+HCOO]- | 8.58 | 973.50 | 973.50 |
| **9** | 27013-91-8 | α-Hederin | C_41_H_66_O_12_ | [M+HCOO]- | 12.50 | 795.45 | 795.50 |
| **10** | 33289-85-9 | Dipsacoside B | C_53_H_86_O_22_ | [M+HCOO]- | 7.12 | 1119.56 | 1119.56 |
| **23** | 96990-19-1 | Momordin IIc | C_41_H64O_13_ | [M-H]- | 12.65 | 763.18 | 763.18 |
| **24** | 20853-58-1 | Hederoside D2 | C_41_H_66_O_13_ | [M+HCOO]- | 10.84 | 811.25 | 811.25 |
| **25** | 78285-90-2 | Ecliptasaponin A | C_36_H_58_O_9_ | [M-H]- | 11.04 | 633.22 | 633.22 |
| **26** | 35286-59-0 | Ziyuglycoside II | C_35_H_56_O_8_ | [M-H]- | 12.52 | 603.31 | 603.31 |
| **27** | 129724-84-1 | Anemoside A3 | C_41_H_66_O_12_ | [M-H]- | 11.08 | 749.37 | 749.37 |

**Supplementary Table 2. Sampling information and asperosaponin Ⅵ content in roots of 209 *D. asperoides* accessions.**

| **Code** | **Content (mg/g)** | **Origin** | **Growth age (year)** | **Collection time** | **Code** | **Content (mg/g)** | **Origin** | **Growth age (year)** | **Collection time** | **Code** | **Content (mg/g)** | **Origin** | **Growth age (year)** | **Collection time** |
| --- | --- | --- | --- | --- | --- | --- | --- | --- | --- | --- | --- | --- | --- | --- |
| **14-4** | 177.13632 | Baoshan,Yunnan Province, China | 1Y | 2022.10 | **9-4** | 80.38560 | Baoshan,Yunnan Province, China | 1Y | 2022.10 | **XD-1** | 60.63612 | Kunming,Yunnan Province, China | 2Y | 2022.07 |
| **10-2F2** | 171.20736 | Baoshan,Yunnan Province, China | 1Y | 2022.10 | **8-6F** | 79.69020 | Baoshan,Yunnan Province, China | 1Y | 2022.10 | **Yl-17** | 60.48342 | Kunming,Yunnan Province, China | 1Y | 2022.07 |
| **15-5** | 169.62252 | Baoshan,Yunnan Province, China | 1Y | 2022.10 | **10-18** | 77.19936 | Baoshan,Yunnan Province, China | 1Y | 2022.10 | **Lq-7** | 48.74460 | Kunming,Yunnan Province, China | 2Y | 2022.07 |
| **14-3** | 167.82504 | Baoshan,Yunnan Province, China | 1Y | 2022.10 | **5-13F** | 77.04162 | Baoshan,Yunnan Province, China | 1Y | 2022.10 | **Lq-3** | 46.43394 | Kunming,Yunnan Province, China | 2Y | 2022.07 |
| **6-12F** | 157.95000 | Baoshan,Yunnan Province, China | 2Y | 2022.10 | **11-17** | 76.48752 | Baoshan,Yunnan Province, China | 1Y | 2022.10 | **XD-8** | 42.15714 | Kunming,Yunnan Province, China | 2Y | 2022.07 |
| **8-27F** | 155.32800 | Baoshan,Yunnan Province, China | 2Y | 2022.10 | **6-6F** | 75.23034 | Baoshan,Yunnan Province, China | 1Y | 2022.10 | **XD-17** | 42.05772 | Kunming,Yunnan Province, China | 2Y | 2022.07 |
| **15-5** | 151.93554 | Baoshan,Yunnan Province, China | 2Y | 2022.10 | **5-14F2** | 73.95486 | Baoshan,Yunnan Province, China | 1Y | 2022.10 | **Lq-9** | 40.76502 | Kunming,Yunnan Province, China | 2Y | 2022.07 |
| **8-7F** | 143.02146 | Baoshan,Yunnan Province, China | 1Y | 2022.10 | **12-8F** | 72.06816 | Baoshan,Yunnan Province, China | 1Y | 2022.10 | **Yl-10** | 31.80840 | Kunming,Yunnan Province, China | 1Y | 2022.07 |
| **16-12** | 142.63458 | Baoshan,Yunnan Province, China | 1Y | 2022.10 | **11-14** | 71.47074 | Baoshan,Yunnan Province, China | 1Y | 2022.10 | **Lj-8** | 171.64512 | Lijiang,Yunnan Province, China | 1Y | 2022.11 |
| **13-6** | 140.42346 | Baoshan,Yunnan Province, China | 1Y | 2022.10 | **13-9** | 70.35684 | Baoshan,Yunnan Province, China | 1Y | 2022.10 | **Lj-23** | 160.61844 | Lijiang,Yunnan Province, China | 1Y | 2022.11 |
| **13-6F** | 138.82056 | Baoshan,Yunnan Province, China | 1Y | 2022.10 | **6-5F** | 68.59854 | Baoshan,Yunnan Province, China | 1Y | 2022.10 | **Lj-31** | 153.77142 | Lijiang,Yunnan Province, China | 1Y | 2022.11 |
| **16-2** | 132.73050 | Baoshan,Yunnan Province, China | 1Y | 2022.10 | **11-20** | 67.77396 | Baoshan,Yunnan Province, China | 1Y | 2022.10 | **Lj-4** | 147.04800 | Lijiang,Yunnan Province, China | 1Y | 2022.11 |
| **10-23** | 130.32000 | Baoshan,Yunnan Province, China | 1Y | 2022.10 | **8-13F** | 65.12340 | Baoshan,Yunnan Province, China | 1Y | 2022.10 | **Lj-32** | 131.98800 | Lijiang,Yunnan Province, China | 1Y | 2022.11 |
| **8-8F** | 129.78600 | Baoshan,Yunnan Province, China | 1Y | 2022.10 | **14-1** | 61.72518 | Baoshan,Yunnan Province, China | 1Y | 2022.10 | **Lj-6** | 131.63550 | Lijiang,Yunnan Province, China | 1Y | 2022.11 |
| **3-15** | 128.45220 | Baoshan,Yunnan Province, China | 1Y | 2022.10 | **8-25F** | 61.27320 | Baoshan,Yunnan Province, China | 1Y | 2022.10 | **Lj-13** | 130.98000 | Lijiang,Yunnan Province, China | 1Y | 2022.11 |
| **14-9** | 128.40384 | Baoshan,Yunnan Province, China | 1Y | 2022.10 | **4-3** | 52.91424 | Baoshan,Yunnan Province, China | 1Y | 2022.10 | **Lj-5** | 124.71270 | Lijiang,Yunnan Province, China | 1Y | 2022.11 |
| **10-19** | 125.92200 | Baoshan,Yunnan Province, China | 1Y | 2022.10 | **9-2** | 52.46556 | Baoshan,Yunnan Province, China | 1Y | 2022.10 | **Lj-14** | 122.41800 | Lijiang,Yunnan Province, China | 1Y | 2022.11 |
| **13-14** | 125.90442 | Baoshan,Yunnan Province, China | 1Y | 2022.10 | **3-11** | 50.62860 | Baoshan,Yunnan Province, China | 1Y | 2022.10 | **Lj-12** | 120.89400 | Lijiang,Yunnan Province, China | 1Y | 2022.11 |
| **11-2** | 125.57010 | Baoshan,Yunnan Province, China | 1Y | 2022.10 | **4-9F** | 48.99870 | Baoshan,Yunnan Province, China | 1Y | 2022.10 | **Lj-22** | 112.89240 | Lijiang,Yunnan Province, China | 1Y | 2022.11 |
| **2-9F** | 118.82928 | Baoshan,Yunnan Province, China | 1Y | 2022.10 | **4-4** | 42.11532 | Baoshan,Yunnan Province, China | 1Y | 2022.10 | **Lj-24** | 104.76600 | Lijiang,Yunnan Province, China | 1Y | 2022.11 |
| **6-9F** | 118.55598 | Baoshan,Yunnan Province, China | 1Y | 2022.10 | **13-11** | 42.08502 | Baoshan,Yunnan Province, China | 1Y | 2022.10 | **Lj-30** | 103.59816 | Lijiang,Yunnan Province, China | 1Y | 2022.11 |
| **2-9F** | 114.63588 | Baoshan,Yunnan Province, China | 1Y | 2022.10 | **11-7** | 40.31352 | Baoshan,Yunnan Province, China | 1Y | 2022.10 | **Lj-26** | 97.83000 | Lijiang,Yunnan Province, China | 1Y | 2022.11 |
| **3-1** | 114.63588 | Baoshan,Yunnan Province, China | 1Y | 2022.10 | **3-6** | 36.78102 | Baoshan,Yunnan Province, China | 1Y | 2022.10 | **Lj-7** | 97.48632 | Lijiang,Yunnan Province, China | 1Y | 2022.11 |
| **12-9F2** | 112.84200 | Baoshan,Yunnan Province, China | 1Y | 2022.10 | **10-11** | 35.00514 | Baoshan,Yunnan Province, China | 1Y | 2022.10 | **Lj-29** | 97.04520 | Lijiang,Yunnan Province, China | 1Y | 2022.11 |
| **3-4** | 110.73684 | Baoshan,Yunnan Province, China | 1Y | 2022.10 | **10-9** | 27.25638 | Baoshan,Yunnan Province, China | 1Y | 2022.10 | **Lj-18** | 93.59724 | Lijiang,Yunnan Province, China | 1Y | 2022.11 |
| **8-14F** | 110.58186 | Baoshan,Yunnan Province, China | 1Y | 2022.10 | **14-10** | 22.03590 | Baoshan,Yunnan Province, China | 1Y | 2022.10 | **Lj-15** | 93.53976 | Lijiang,Yunnan Province, China | 1Y | 2022.11 |
| **7-8F** | 110.19276 | Baoshan,Yunnan Province, China | 1Y | 2022.10 | **L1** | 7.47054 | Baoshan,Yunnan Province, China | 1Y | 2022.10 | **Lj-10** | 91.93800 | Lijiang,Yunnan Province, China | 1Y | 2022.11 |
| **8-26F** | 110.12802 | Baoshan,Yunnan Province, China | 1Y | 2022.10 | **Lq-5** | 190.65000 | Kunming,Yunnan Province, China | 2Y | 2022.07 | **Lj-21** | 90.53322 | Lijiang,Yunnan Province, China | 1Y | 2022.11 |
| **5-8F** | 110.04552 | Baoshan,Yunnan Province, China | 1Y | 2022.10 | **XD-3** | 158.71704 | Kunming,Yunnan Province, China | 2Y | 2022.07 | **Lj-28** | 90.36000 | Lijiang,Yunnan Province, China | 1Y | 2022.11 |
| **5-10F** | 107.04000 | Baoshan,Yunnan Province, China | 1Y | 2022.10 | **XD-7** | 149.36400 | Kunming,Yunnan Province, China | 2Y | 2022.07 | **Lj-19** | 87.67068 | Lijiang,Yunnan Province, China | 1Y | 2022.11 |
| **15-13** | 105.94860 | Baoshan,Yunnan Province, China | 1Y | 2022.10 | **Lq-6** | 132.44400 | Kunming,Yunnan Province, China | 2Y | 2022.07 | **Lj-34** | 85.71636 | Lijiang,Yunnan Province, China | 1Y | 2022.11 |
| **7-5F** | 105.32400 | Baoshan,Yunnan Province, China | 1Y | 2022.10 | **Lq-1** | 132.36834 | Kunming,Yunnan Province, China | 2Y | 2022.07 | **Lj-17** | 84.84306 | Lijiang,Yunnan Province, China | 1Y | 2022.11 |
| **12-4F** | 105.17574 | Baoshan,Yunnan Province, China | 1Y | 2022.10 | **Yl-6** | 128.02242 | Kunming,Yunnan Province, China | 1Y | 2022.07 | **Lj-9** | 78.64800 | Lijiang,Yunnan Province, China | 1Y | 2022.11 |
| **2-13F** | 105.09000 | Baoshan,Yunnan Province, China | 1Y | 2022.10 | **Yl-11** | 125.93916 | Kunming,Yunnan Province, China | 1Y | 2022.07 | **Lj-33** | 78.62694 | Lijiang,Yunnan Province, China | 1Y | 2022.11 |
| **10-13** | 104.93400 | Baoshan,Yunnan Province, China | 1Y | 2022.10 | **Yl-14** | 124.81200 | Kunming,Yunnan Province, China | 1Y | 2022.07 | **Lj-2** | 74.94000 | Lijiang,Yunnan Province, China | 1Y | 2022.11 |
| **11-18** | 102.76068 | Baoshan,Yunnan Province, China | 1Y | 2022.10 | **Yl-1** | 119.17200 | Kunming,Yunnan Province, China | 1Y | 2022.07 | **Lj-1** | 74.19786 | Lijiang,Yunnan Province, China | 1Y | 2022.11 |
| **4-1** | 102.42816 | Baoshan,Yunnan Province, China | 1Y | 2022.10 | **XD-9** | 111.97182 | Kunming,Yunnan Province, China | 2Y | 2022.07 | **Lj-11** | 68.60046 | Lijiang,Yunnan Province, China | 1Y | 2022.11 |
| **8-24F** | 102.23400 | Baoshan,Yunnan Province, China | 1Y | 2022.10 | **Lq-13** | 111.25800 | Kunming,Yunnan Province, China | 2Y | 2022.07 | **Lj-16** | 67.16658 | Lijiang,Yunnan Province, China | 1Y | 2022.11 |
| **7-9F** | 102.08400 | Baoshan,Yunnan Province, China | 1Y | 2022.10 | **XD-11** | 110.55600 | Kunming,Yunnan Province, China | 2Y | 2022.07 | **Lj-3** | 65.98819 | Lijiang,Yunnan Province, China | 1Y | 2022.11 |
| **8-19F** | 100.72434 | Baoshan,Yunnan Province, China | 1Y | 2022.10 | **XD-15** | 109.45614 | Kunming,Yunnan Province, China | 2Y | 2022.07 | **Lj-25** | 59.99676 | Lijiang,Yunnan Province, China | 1Y | 2022.11 |
| **4-10F** | 100.38096 | Baoshan,Yunnan Province, China | 1Y | 2022.10 | **Lq-11** | 109.05888 | Kunming,Yunnan Province, China | 2Y | 2022.07 | **Lj-20** | 55.91712 | Lijiang,Yunnan Province, China | 1Y | 2022.11 |
| **9-3** | 100.31460 | Baoshan,Yunnan Province, China | 1Y | 2022.10 | **Yl-15** | 107.18400 | Kunming,Yunnan Province, China | 1Y | 2022.07 | **Lj-27** | 37.12356 | Lijiang,Yunnan Province, China | 1Y | 2022.11 |
| **5-11F** | 100.23996 | Baoshan,Yunnan Province, China | 1Y | 2022.10 | **Lq-17** | 106.08660 | Kunming,Yunnan Province, China | 2Y | 2022.07 | **LC-8** | 142.49016 | Puer,Yunnan Province, China | 2Y | 2021.03 |
| **12-9F** | 99.79800 | Baoshan,Yunnan Province, China | 1Y | 2022.10 | **Lq-12** | 105.77244 | Kunming,Yunnan Province, China | 2Y | 2022.07 | **LC-13** | 141.34710 | Puer,Yunnan Province, China | 2Y | 2021.03 |
| **6-18F** | 99.60294 | Baoshan,Yunnan Province, China | 1Y | 2022.10 | **Lq-14** | 104.89800 | Kunming,Yunnan Province, China | 2Y | 2022.07 | **LC-1** | 128.10948 | Puer,Yunnan Province, China | 2Y | 2021.03 |
| **8-8F** | 99.32400 | Baoshan,Yunnan Province, China | 1Y | 2022.10 | **Lq-15** | 100.59000 | Kunming,Yunnan Province, China | 2Y | 2022.07 | **LC-15** | 125.90580 | Puer,Yunnan Province, China | 2Y | 2021.03 |
| **10-12** | 97.30800 | Baoshan,Yunnan Province, China | 1Y | 2022.10 | **XD-16** | 98.69394 | Kunming,Yunnan Province, China | 2Y | 2022.07 | **LC-5** | 120.67200 | Puer,Yunnan Province, China | 2Y | 2021.03 |
| **15-4** | 96.91344 | Baoshan,Yunnan Province, China | 1Y | 2022.10 | **Yl-5** | 98.27844 | Kunming,Yunnan Province, China | 1Y | 2022.07 | **LC-16** | 120.19800 | Puer,Yunnan Province, China | 2Y | 2021.03 |
| **2-7F** | 96.68490 | Baoshan,Yunnan Province, China | 1Y | 2022.10 | **Lq-10** | 97.97688 | Kunming,Yunnan Province, China | 2Y | 2022.07 | **LC-22** | 115.30200 | Puer,Yunnan Province, China | 2Y | 2021.03 |
| **10-14** | 96.67800 | Baoshan,Yunnan Province, China | 1Y | 2022.10 | **XD-10** | 95.12226 | Kunming,Yunnan Province, China | 2Y | 2022.07 | **LC-21** | 110.43696 | Puer,Yunnan Province, China | 2Y | 2021.03 |
| **5-2F** | 95.44584 | Baoshan,Yunnan Province, China | 1Y | 2022.10 | **Yl-3** | 95.10000 | Kunming,Yunnan Province, China | 1Y | 2022.07 | **LC-25** | 109.53240 | Puer,Yunnan Province, China | 2Y | 2021.03 |
| **3-8** | 94.90200 | Baoshan,Yunnan Province, China | 1Y | 2022.10 | **XD-12** | 93.08040 | Kunming,Yunnan Province, China | 2Y | 2022.07 | **LC-26** | 106.40400 | Puer,Yunnan Province, China | 2Y | 2021.03 |
| **8-21F** | 94.84878 | Baoshan,Yunnan Province, China | 1Y | 2022.10 | **XD-4** | 92.91318 | Kunming,Yunnan Province, China | 2Y | 2022.07 | **LC-20** | 106.31700 | Puer,Yunnan Province, China | 2Y | 2021.03 |
| **7-6F** | 93.90600 | Baoshan,Yunnan Province, China | 1Y | 2022.10 | **Lq-4** | 92.75400 | Kunming,Yunnan Province, China | 1Y | 2022.07 | **LC-27** | 105.86400 | Puer,Yunnan Province, China | 2Y | 2021.03 |
| **3-7** | 93.89400 | Baoshan,Yunnan Province, China | 1Y | 2022.10 | **Lq-7** | 91.62672 | Kunming,Yunnan Province, China | 1Y | 2022.07 | **LC-4** | 105.18000 | Puer,Yunnan Province, China | 2Y | 2021.03 |
| **5-7F** | 93.77178 | Baoshan,Yunnan Province, China | 1Y | 2022.10 | **Lq-9** | 91.50000 | Kunming,Yunnan Province, China | 1Y | 2022.07 | **LC-2** | 101.49600 | Puer,Yunnan Province, China | 2Y | 2021.03 |
| **6-4F** | 93.70200 | Baoshan,Yunnan Province, China | 1Y | 2022.10 | **Lq-8** | 87.45840 | Kunming,Yunnan Province, China | 2Y | 2022.07 | **LC-10** | 100.10208 | Puer,Yunnan Province, China | 2Y | 2021.03 |
| **9-5** | 92.54400 | Baoshan,Yunnan Province, China | 1Y | 2022.10 | **XD-14** | 85.95444 | Kunming,Yunnan Province, China | 2Y | 2022.07 | **LC-9** | 95.62122 | Puer,Yunnan Province, China | 2Y | 2021.03 |
| **6-15F** | 92.50110 | Baoshan,Yunnan Province, China | 1Y | 2022.10 | **XD-2** | 84.14112 | Kunming,Yunnan Province, China | 2Y | 2022.07 | **LC-18** | 95.50266 | Puer,Yunnan Province, China | 2Y | 2021.03 |
| **6-2F** | 92.45514 | Baoshan,Yunnan Province, China | 1Y | 2022.10 | **Yl-16** | 81.23646 | Kunming,Yunnan Province, China | 1Y | 2022.07 | **LC-6** | 92.59284 | Puer,Yunnan Province, China | 2Y | 2021.03 |
| **6-19F** | 91.17900 | Baoshan,Yunnan Province, China | 1Y | 2022.10 | **Lq-4** | 80.62980 | Kunming,Yunnan Province, China | 2Y | 2022.07 | **LC-23** | 90.30000 | Puer,Yunnan Province, China | 2Y | 2021.03 |
| **4-13** | 90.97092 | Baoshan,Yunnan Province, China | 1Y | 2022.10 | **Lq-16** | 77.83548 | Kunming,Yunnan Province, China | 2Y | 2022.07 | **LC-24** | 85.89102 | Puer,Yunnan Province, China | 2Y | 2021.03 |
| **5-18F** | 90.81546 | Baoshan,Yunnan Province, China | 1Y | 2022.10 | **XD-13** | 77.66874 | Kunming,Yunnan Province, China | 2Y | 2022.07 | **LC-17** | 84.18348 | Puer,Yunnan Province, China | 2Y | 2021.03 |
| **6-21F** | 90.80640 | Baoshan,Yunnan Province, China | 1Y | 2022.10 | **Yl-8** | 75.34440 | Kunming,Yunnan Province, China | 1Y | 2022.07 | **LC-11** | 81.92166 | Puer,Yunnan Province, China | 2Y | 2021.03 |
| **7-2F** | 89.93226 | Baoshan,Yunnan Province, China | 1Y | 2022.10 | **Yl-12** | 75.26616 | Kunming,Yunnan Province, China | 1Y | 2022.07 | **LC-3** | 77.88936 | Puer,Yunnan Province, China | 2Y | 2021.03 |
| **8-5F** | 87.75096 | Baoshan,Yunnan Province, China | 1Y | 2022.10 | **XD-5** | 73.07730 | Kunming,Yunnan Province, China | 2Y | 2022.07 | **LC-19** | 71.04000 | Puer,Yunnan Province, China | 2Y | 2021.03 |
| **5-24F** | 87.52344 | Baoshan,Yunnan Province, China | 1Y | 2022.10 | **Yl-13** | 72.12750 | Kunming,Yunnan Province, China | 1Y | 2022.07 | **LC-14** | 57.44916 | Puer,Yunnan Province, China | 2Y | 2021.03 |
| **6-6F2** | 83.97114 | Baoshan,Yunnan Province, China | 1Y | 2022.10 | **XD-6** | 67.16796 | Kunming,Yunnan Province, China | 2Y | 2022.07 | **LC-12** | 55.50900 | Puer,Yunnan Province, China | 2Y | 2021.03 |
| **7-16F** | 82.38816 | Baoshan,Yunnan Province, China | 1Y | 2022.10 | **Lq-2** | 65.88540 | Kunming,Yunnan Province, China | 2Y | 2022.07 | **LC-7** | 37.73676 | Puer,Yunnan Province, China | 2Y | 2021.03 |
| **7-1F** | 80.52150 | Baoshan,Yunnan Province, China | 1Y | 2022.10 | **Yl-2** | 65.78988 | Kunming,Yunnan Province, China | 1Y | 2022.07 |  |  |  |  |  |

**Supplementary Table 3.** **Statistics of the contig-level and chromosome-level** ***D. asperoides* genome assembly**.

| Item | The contig-level genome assembly | | The contig-level genome assembly (purged) | | | The chromosome-level genome assembly |
| --- | --- | --- | --- | --- | --- | --- |
| Total bases | 5,303,701,501 | | | 5,248,957,368 | 5,227,414,451 | |
| Max sequence length (bp) | 173,897,702 | | | 173,897,702 | 647,486,188 | |
| Average length | | 2953063 | | 6686569 | 12626604 | |
| N20 length | | 80,939,096 | | 80,939,096 | 646,935,175 | |
| N30 length | | 54,511,083 | | 54,511,083 | 613,228,167 | |
| N50 length | | 31,394,640 | | 32,451,949 | 545,804,658 | |
| N70 length | | 17,350,956 | | 17,750,568 | 520,105,756 | |
| N90 length | | 6,491,918 | | 7,175,527 | 500,811,633 | |

**Supplementary Table 4. Statistics of** ***D. asperoides* genome assembly.**

| **Hi-C** | **Length(bp)** | **congtifs** | **Genes** | **Telomere** |
| --- | --- | --- | --- | --- |
| Chr1 | 647,486,188 | 31 | 4,380 | Right |
| Chr2 | 646,935,175 | 26 | 6,850 | Right |
| Chr3 | 613,228,167 | 32 | 6,554 | Left |
| Chr4 | 569,350,932 | 27 | 4,670 | Both |
| Chr5 | 545,804,658 | 36 | 4,557 | Both |
| Chr6 | 541,499,446 | 29 | 4,344 | Right |
| Chr7 | 520,105,756 | 53 | 3,637 | Both |
| Chr8 | 516,718,050 | 36 | 3,385 | Both |
| Chr9 | 500,811,633 | 13 | 4,479 | Left |
| Total | 5,101,940,005 | 283 | 42,856 | 13 |
| un-anchored | 125,474,446 | 412 | 0 |  |

**Supplementary Table 5. Assessment of the** ***D. asperoides* genome quality using Benchmarking Universal Single-Copy Orthologs (BUSCO).**

| **Type** | **Number** | **Percent (%)** |
| --- | --- | --- |
| Complete (C) | 1600 | 99.1 |
| Complete and single-copy (S) | 1292 | 80.0 |
| Complete and duplicated (D) | 308 | 19.1 |
| Fragmented (F) | 7 | 0.4 |
| Missing (M) | 7 | 0.5 |
| Total BUSCO groups searched | 1614 | 100 |

**Supplementary Table 6. Repeated sequence annotations of *D. asperoides.***

| Species | *D. asperoides* | | | |
| --- | --- | --- | --- | --- |
| total length: | 5,227,414,451 | | | |
| GC level: | 36.72% | | | |
| bases masked: | | 4,763,581,268 | | |
|  | | number of elements | length(bp) | percentage |
| Retroelements | | 4,675,927 | 4,148,436,563 | 79.36% |
| SINEs: | | 0 | 0 | 0% |
| Penelope | | 0 | 0 | 0% |
| LINEs: | | 13715 | 4543592 | 0.09% |
| CRE/SLACS | | 0 | 0 | 0% |
| L2/CR1/Rex | | 0 | 0 | 0% |
| R1/LOA/Jockey | | 0 | 0 | 0% |
| R2/R4/NeSL | | 0 | 0 | 0% |
| RTE/Bov-B | | 0 | 0 | 0% |
| L1/CIN4 | | 0 | 0 | 0% |
| LTR elements: | | 4,662,212 | 4,143,892,971 | 79.27% |
| BEL/Pao | | 0 | 0 | 0% |
| Ty1/Copia | | 732,108 | 361,620,113 | 6.92% |
| Gypsy/DIRS1 | | 2,066,577 | 2,836,732,904 | 54.27% |
| Retroviral | | 0 | 0 | 0% |
| DNA transposons | | 1,759,489 | 371,837,923 | 7.11% |
| hobo-Activator | | 0 | 0 | 0% |
| Tc1-IS630-Pogo | | 0 | 0 | 0% |
| En-Spm | | 0 | 0 | 0% |
| MuDR-IS905 | | 0 | 0 | 0% |
| PiggyBac | | 0 | 0 | 0% |
| Other (Mirage, Pelement, Transib) | | 0 | 0 | 0% |
| Rolling-circles | | 0 | 0 | 0% |
| Unclassified: | | 1,291,305 | 243,306,782 | 4.65% |
| Total interspersed repeats: | |  | 4,763,581,268 | 91.13% |
| Small RNA: | | 0 | 0 | 0% |
| Satellites: | | 0 | 0 | 0% |
| Simple repeats | | 0 | 0 | 0% |
| Low complexity | | 0 | 0 | 0% |

**Supplementary Table 7. Repeated sequence annotations of *T. glandulifera.***

| Species | *T. glandulifera* | | | |
| --- | --- | --- | --- | --- |
| total length: | 680,371,130 bp | | | |
| GC level: | 37.16 % | | | |
| bases masked: | | 463,556,361 bp | | |
|  | | number of elements | length(bp) | percentage |
| Retroelements | | 333,420 | 349,504,857 | 51.37% |
| SINEs: | | 0 | 0 | 0% |
| Penelope | | 0 | 0 | 0% |
| LINEs: | | 0 | 0 | 0% |
| CRE/SLACS | | 0 | 0 | 0% |
| L2/CR1/Rex | | 0 | 0 | 0% |
| R1/LOA/Jockey | | 0 | 0 | 0% |
| R2/R4/NeSL | | 0 | 0 | 0% |
| RTE/Bov-B | | 0 | 0 | 0% |
| L1/CIN4 | | 0 | 0 | 0% |
| LTR elements: | | 333,420 | 349,504,857 | 51.37% |
| BEL/Pao | | 0 | 0 | 0% |
| Ty1/Copia | | 91,713 | 85,662,083 | 12.59% |
| Gypsy/DIRS1 | | 136,566 | 182,704,258 | 26.85% |
| Retroviral | | 0 | 0 | 0% |
| DNA transposons | | 375,895 | 104,569,392 | 15.37% |
| hobo-Activator | | 0 | 0 | 0% |
| Tc1-IS630-Pogo | | 0 | 0 | 0% |
| En-Spm | | 0 | 0 | 0% |
| MuDR-IS905 | | 0 | 0 | 0% |
| PiggyBac | | 0 | 0 | 0% |
| Other (Mirage, Pelement, Transib) | | 0 | 0 | 0% |
| Rolling-circles | | 0 | 0 | 0% |
| Unclassified: | | 65,225 | 9,482,112 | 1.39% |
| Total interspersed repeats: | |  | 463,556,361 | 68.13% |
| Small RNA: | | 0 | 0 | 0% |
| Satellites: | | 0 | 0 | 0% |
| Simple repeats | | 0 | 0 | 0% |
| Low complexity | | 0 | 0 | 0% |

**Supplementary Table 8. Repeated sequence annotations of *L. japonica.***

| Species | *L. japonica* | | | |
| --- | --- | --- | --- | --- |
| total length: | 903,813,177 bp | | | |
| GC level: | 34.11 % | | | |
| bases masked: | | 480,966,400 bp | | |
|  | | number of elements | length(bp) | percentage |
| Retroelements | | 656,023 | 385,669,060 | 42.67% |
| SINEs: | | 0 | 0 | 0% |
| Penelope | | 0 | 0 | 0% |
| LINEs: | | 0 | 0 | 0% |
| CRE/SLACS | | 0 | 0 | 0% |
| L2/CR1/Rex | | 0 | 0 | 0% |
| R1/LOA/Jockey | | 0 | 0 | 0% |
| R2/R4/NeSL | | 0 | 0 | 0% |
| RTE/Bov-B | | 0 | 0 | 0% |
| L1/CIN4 | | 0 | 0 | 0% |
| LTR elements: | | 656023 | 385,669,060 | 42.67% |
| BEL/Pao | | 0 | 0 | 0% |
| Ty1/Copia | | 259,429 | 139,090,689 | 15.39% |
| Gypsy/DIRS1 | | 137,745 | 146,158,853 | 16.17% |
| Retroviral | | 0 | 0 | 0% |
| DNA transposons | | 491,946 | 95,296,230 | 10.54% |
| hobo-Activator | | 4,581 | 1,220,417 | 0.14% |
| Tc1-IS630-Pogo | | 0 | 0 | 0% |
| En-Spm | | 0 | 0 | 0% |
| MuDR-IS905 | | 0 | 0 | 0% |
| PiggyBac | | 0 | 0 | 0% |
| Tourist/Harbinger | | 35 | 2,357 | 0.00 % |
| Other (Mirage, Pelement, Transib) | | 0 | 0 | 0% |
| Rolling-circles | | 0 | 0 | 0% |
| Unclassified: | | 3 | 135 | 0% |
| Total interspersed repeats: | |  | 480,965,425 | 53.22% |
| Small RNA: | | 0 | 0 | 0% |
| Satellites: | | 0 | 0 | 0% |
| Simple repeats | | 0 | 0 | 0% |
| Low complexity | | 0 | 0 | 0% |

**Supplementary Table 9. Analysis of CDS, exons, and introns in *D. asperoides.***

| State | Total | Average |
| --- | --- | --- |
| CDS lengths | 48569910 | 1,133.38 |
| Exon lengths | NA | 302.89 |
| Intron lengths | NA | 850.89 |

**Supplementary Table 10. Number of predicted protein-coding genes with homology or functional classification by different methods for *Dipsacus asperoides*.**

| Annotation Database | Number of Annotations | Annotation Percentage |
| --- | --- | --- |
| Nr | 40,438 | 94.36% |
| KEGG | 40,231 | 93.88% |
| KOG | 38,005 | 88.68% |
| TrEMBL | 40,392 | 94.25% |
| SwissProt | 32,282 | 75.33% |
| Pfam | 32,532 | 75.91% |
| InterProScan | 40,579 | 94.69% |
| KO | 15,801 | 36.87% |
| GO | 26,578 | 62.02% |
| **All** | **41,558** | **96.98%** |
| Total | 42,856 | 100.00% |

**Supplementary Table 11. Non-coding RNAs predicted of the *D. asperoides* genome.**

| Non-coding RNAs | Number |
| --- | --- |
| tRNA | 2,863 |
| rRNA | 19,627 |
| snRNA | 843 |
| miRNA | 172 |

**Supplementary Table 12. Annotation-BUSCO of *D. asperoides*.**

| **Type** | **Percent (%)** |
| --- | --- |
| Complete (C) | 97.1 |
| Complete and single-copy (S) | 80.1 |
| Complete and duplicated (D) | 17.0 |
| Fragmented (F) | 0.9 |
| Missing (M) | 2.0 |

**Supplementary Table 13. LC–MS analysis of triterpenoids and triterpenoid saponins in plant samples.**

| No. | CAS No. | Compound | Formula | Pseudomolecular  ion | Retention time  (min) | Standard *m/z* | Detected in plant samples |
| --- | --- | --- | --- | --- | --- | --- | --- |
| **4** | 508-02-1 | Oleanolic acid | C₃₀H₄₈O₃ | [M-H]- | 23.50 | 455.35 | 455.35 |
| **5** | 465-99-6 | Hederagenin | C₃₀H₄₈O₄ | [M-H]- | 18.19 | 471.34 | 471.34 |
| **6** | 631-01-6 | Cauloside A | C_35_H_56_O_8_ | [M+HCOO]- | 16.58 | 649.39 | 649.39 |
| **7** | 39524-13-5 | HN saponin F | C_41_H_66_O_13_ | [M+HCOO]- | 14.71 | 811.44 | 811.44 |
| **8** | 39524-08-8 | Asperosaponin Ⅵ | C_47_H_76_O_18_ | [M+HCOO]- | 13.94 | 973.50 | 973.50 |
| **9** | 27013-91-8 | α-Hederin | C_41_H_66_O_12_ | [M+HCOO]- | 15.94 | 795.45 | 795.45 |
| **10** | 33289-85-9 | Dipsacoside B | C_53_H_86_O_22_ | [M+HCOO]- | 13.72 | 1119.56 | 1119.55 |
| **11** | 140360-29-8 | Macranthoidin A | C_59_H_96_O_27_ | [M-H]- | 11.30 | 1235.60 | 1235.60 |
| **12** | 136849-88-2 | Macranthoidin B | C_65_H_106_O_32_ | [M-H]- | 8.97 | 1397.65 | ND |
| **13** | 510-30-5 | Echinocystic acid | C_30_H_48_O_4_ | [M-H]- | 18.37 | 471.34 | 471.34 |
| **14** | 4373-41-5 | Maslinic acid | C₃₀H₄₈O₄ | [M-H]- | 19.34 | 471.34 | 471.34 |
| **15** | 639-14-5 | Gypsogenin | C₃₀H₄₆O_4_ | [M-H]- | 18.98 | 469.33 | 469.33 |
| **16** | 77-52-1 | Ursolic acid | C30H48O3 | [M-H]- | 23.80 | 455.35 | 455.35 |
| **17** | 4547-24-4 | Corosolic acid | C₃₀H₄₈O₄ | [M-H]- | 19.10 | 471.34 | 471.34 |
| **18** | 464-92-6 | Asiatic acid | C₃₀H₄₈O₅ | [2M-H]- | 16.33 | 975.69 | 975.68 |
| **19** | 34540-22-2 | Asiaticoside A | C₄₈H₇₈O_20_ | [M+HCOO]- | 5.02 | 1019.50 | ND |
| **20** | 26020-14-4 | Calenduloside E | C₃₆H₅_6_O_9_ | [M-H]- | 16.80 | 631.38 | ND |
| **21** | 80930-74-1 | Zingibroside R1 | C_42_H_66_O_14_ | [M-H]- | 15.72 | 793.43 | ND |
| **22** | 34367-04-9 | Ginsenoside Ro | C₄₈H₇₆O₁₉ | [M-H]- | 14.28 | 955.49 | ND |

**Supplementary Table 14. Pairwise sequence identity matrix of UGT85CA family gene**

| **Gene ID** | **Lj1A1075T77.1** | **Lj1C1075T1.1** | **Lj1C1074T6.1** | **Lj1A1074T44.1** | **Das07G002907.t1** | **Das07G002596.t1** | **TgChr04G16777.1** | **TgChr04G16778.1** |
| --- | --- | --- | --- | --- | --- | --- | --- | --- |
| **Lj1A1075T77.1** | **100.00%** |  |  |  |  |  |  |  |
| **Lj1C1075T1.1** | **61.38%** | **100.00%** |  |  |  |  |  |  |
| **Lj1C1074T6.1** | **57.29%** | **76.72%** | **100.00%** |  |  |  |  |  |
| **Lj1A1074T44.1** | **58.75%** | **77.75%** | **93.79%** | **100.00%** |  |  |  |  |
| **Das07G002907.t1** | **54.26%** | **64.86%** | **69.29%** | **70.95%** | **100.00%** |  |  |  |
| **Das07G002596.t1** | **55.93%** | **67.78%** | **72.41%** | **73.65%** | **78.67%** | **100.00%** |  |  |
| **TgChr04G16777.1** | **57.80%** | **72.56%** | **77.59%** | **78.84%** | **74.74%** | **80.33%** | **100.00%** |  |
| **TgChr04G16778.1** | **57.80%** | **72.56%** | **77.80%** | **79.05%** | **74.53%** | **80.54%** | **99.79%** | **100.00%** |

**Supplementary Table 15. Enrichment of TEs and LTRs within the genomic neighborhood of *DaUGT85CA7.***

| Type | count | Length (bp) | Total length (bp) | sequence ratio (%) |
| --- | --- | --- | --- | --- |
| LTR | 25 | 17438 | 20000 | 87.19% |
| TE | 42 | 18456 | 20000 | 92.28% |
| LTR | 50 | 36220 | 40000 | 90.55% |
| TE | 81 | 37592 | 40000 | 93.98% |
| LTR | 78 | 50842 | 60000 | 84.74% |
| TE | 128 | 55520 | 60000 | 92.53% |
| LTR | 112 | 80537 | 100000 | 80.54% |

**Supplementary Table 16. *Ka* and *Ks* values of *βAS* and *MAS* gene pairs calculated using the WGDI.**

| id1 | id2 | *Ka* | *Ks* |
| --- | --- | --- | --- |
| Das04G003910 | Das07G002904 | 0.0084 | 0.3852 |
| Das03G003102 | Das04G003055 | 0.0205 | 0.3738 |
| Das02G004901 | Das02G001755 | 0.0275 | 0.3572 |
| Das03G004796 | Das09G002328 | 0.0107 | 0.2451 |

**Supplementary Table 17. *Ka* and *Ks* values of *CYP* gene pairs calculated using the WGDI.**

| id1 | id2 | *Ka* | *Ks* |
| --- | --- | --- | --- |
| Das07G001840 | Das07G001837 | 0.0948 | 1.201 |
| Das03G001840 | Das07G001837 | 0.0214 | 0.4424 |
| Das03G003375 | Das09G003295 | 0.0541 | 0.3766 |

**Supplementary Table 18. Primers used in this study.**

| ***Name*** | **Primer sequences** |
| --- | --- |
| *pYES2-DaβAS1-F* | gcttggtaccgagctcggatccATGTGGAGACTGAAGATAGCAGAAG |
| *pYES2-DaβAS1-R* | gcacactggcggccgttactagTTAAGAAAGACATTTTTGGTTGGATG |
| *pYES2-DaβAS2-F* | gcttggtaccgagctcggatccATGTGGAGATTGAAGATAGCTGA |
| *pYES2-DaβAS2-R* | cactggcggccgttactagTTATTAAGAGAGTTGTTCTTTACTGGAC |
| *pYES2-DaβAS3-F* | gcttggtaccgagctcggatccATGTGGAGACTGAAGATAGCTGA |
| *pYES2-DaβAS3-R* | cacactggcggccgttactagCTAGTTCTTATTGGAGGGTAAAGGAA |
| *pYES2-DaMAS1-F* | gcttggtaccgagctcggatccATGTGGAAATTGAAGATTGCAGA |
| *pYES2-DaMAS1-R* | gcacactggcggccgttactagTTATATCTTTTGGCTCGGCAACCAGA |
| *pYES2-DaMAS2-F* | gcttggtaccgagctcggatccATGTGGAAGTTGAAGATTGCTGA |
| *pYES2-DaMAS2-R* | gcacactggcggccgttactagTTAGATCTTTTGGCTTGGCAACCAAA |
| *pYES2-DaMAS3-F* | gcttggtaccgagctcggatccATGTGGAAGCTGAAAATTGCTGA |
| *pYES2-DaMAS3-R* | agcacactggcggccgttactagTCAGATTAATTGATTAGGCACCCAA |
| *y33-DaOAS1-F* | cagtcgacctcgaatctagaATGGAGGTCTTCTATGTCTCTCTC |
| *y33-DaOAS1-R* | tctagagggccgcatcatgTTAAGGTTTGTAGGGAAATAGGCG |
| *y33-DaOAS2-F* | cagtcgacctcgaatctagaATGGAAATCTTCTATGTATCTCTCCTAT |
| *y33-DaOAS2-R* | tctagagggccgcatcatgCAGAAGGCCAAGCGGATTAAGGTTTGTT |
| *y33-DaOAS3-F* | cagtcgacctcgaatctagaATTCATCTCAAACTACAAACACAAGAC |
| *y33-DaOAS3-R* | tctagagggccgcatcatgTTATGGTTTGTGTGGATAGAGGC |
| *pET28a-DaUGT73GZ2-F* | cagcaaatgggtcgcggatccATGGCTTCCAAAACCCAAAAC |
| *pET28a-DaUGT73GZ2-R* | ctcgaattcggatcggatccTCAGTGTAAAATTTCAGCATTCTCAAC |
| *pET28a-DaUGT94CS1-F* | ctggtgccgcgcggcagccatatgTCCTCCATTACCATTTCTAGTTCC |
| *pET28a-DaUGT94CS1-R* | gtggtggtggtggtggtgctcgagCTACTTATTGTTCTTGATAAACTC |
| *pET28a-DaUGT94BA4-F* | ggacagcaaatgggtcgcggatccATGGCCATCAACGAACACCACCGT |
| *pET28a-DaUGT94BA4-R* | tcgacggagctcgaattcggatccTAACACATTGGCATCATTTTTTTC |
| *pET28a-DaUGT94CP2-F* | ggacagcaaatgggtcgcggatccATGGCATCTGAAAAAGCCGATCGT |
| *pET28a-DaUGT94CP2-R* | tcgacggagctcgaattcggatccATTTTTTGGACAAAGTTGCAACAA |
| *pET28a-DaUGT94CP3-F* | ggacagcaaatgggtcgcggatccATGGCATCTGAAAAAGCCGATCGT |
| *pET28a-DaUGT94CP3-R* | tcgacggagctcgaattcggatccATTTTTTGGACAAAGTTGCAACAA |
| *pET28a-DaUGT94CN8-F* | ctggtgccgcgcggcagccatatgATGGATGCTGGTGAGGGCAAATTG |
| *pET28a-DaUGT94CN8-R* | gtggtggtggtggtggtgctcgagTTATTTGGTCTTGTTGCCACAAAG |
| *pET28a-DaUGT94AY3-F* | ctggtgccgcgcggcagccatatgATGGATCCCAAGGAAAAAAGATTA |
| *pET28a-DaUGT94AY3-R* | gtggtggtggtggtggtgctcgagCTATTCTTGTTCTTTGCTTTTCTC |
| *pET28a-DaUGT73GY4-F* | ggacagcaaatgggtcgcggatccATGGCAACTGATGACTTGAAGCTC |
| *pET28a-DaUGT73GY4-R* | tcgacggagctcgaattcggatccAGCTGAAGAAATTTTTTGTTTCTT |
| *pET28a-DaUGT73GY3-F* | ggacagcaaatgggtcgcggatccATGGCAACTGATGACTTGAAGCTC |
| *pET28a-DaUGT73GY3-R* | tcgacggagctcgaattcggatccAGCTGAAGAAATTTTTTGTTTCTT |
| *pET28a-DaUGT2596-F* | cagcaaatgggtcgcggatccATGGAAGCAATTTCAAAGCCAC |
| *pET28a-DaUGT2596-R* | gacggagctcgaattcggatccTTAGGGTTTTGCGAGCAAAAC |
| *pET28a-DaUGT85A245-F* | ggacagcaaatgggtcgcggatccATGAATTCCATTTCAAGAAGACAA |
| *pET28a-DaUGT85A245-R* | tcgacggagctcgaattcggatccGTGCTCTTAGCAAAGTTGATTAATT |
| *pET28a-ljUGT85CA2-F* | ggacagcaaatgggtcgcggatccATGGAAGCCACTTCAAAGC |
| *pET28a-ljUGT85CA2-R* | tcgacggagctcgaattcggatccGTTAACGAAGTTTTACTAAAAAAA |
| *pET28a-ljUGT85CA5-F* | ggacagcaaatgggtcgcggatccATGGACGCAACTTCAAAGCC |
| *pET28a-ljUGT85CA5-R* | tcgacggagctcgaattcggatccCTATCGCAAAAACGTAAGAGCTAA |
| *pET28a-TgUGT85CA3-1-F* | ggacagcaaatgggtcgcggatccATGGAAGCAACTTCACAGCC |
| *pET28a-TgUGT85CA3-1-R* | tcgacggagctcgaattcggatccGAAGTTCTGCTAGCAAAACCTTAA |
| *pET28a-TgUGT85CA3-2-F* | ggacagcaaatgggtcgcggatccATGGAAGCAACTTCACAGCC |
| *pET28a-TgUGT85CA3-2-R* | tcgacggagctcgaattcggatccGAAGTTCTGCTAGCAAAACCTTAA |
| *pEAQ-tHMGR-F* | cccaaattcgcgaccggtATGCCCGAGGAGGACGAGG |
| *pEAQ-tHMGR-R* | tcgaggcctttaactctggGGACATGTCCAAGATCGCCTGCTGA |
| *pEAQ-βAS2-F* | cccaaattcgcgaccggtATGTGGAGATTGAAGATAGCTGA |
| *pEAQ-βAS2-R* | tcgaggcctttaactctggTTATTAAGAGAGTTGTTCTTTACTGGACG |
| *pEAQ-OAS2-F* | cccaaattcgcgaccggtATGGAAATCTTCTATGTATCTCTCCTAT |
| *pEAQ-OAS2-R* | tcgaggcctttaactctggCGCCTAATTCCAAACAAAGCATAA |
| *pEAQ-CYP714E29-F* | cccaaattcgcgaccggtATGGAGATTGCTAGTGTGATG |
| *pEAQ-CYP714E29-R* | tcgaggcctttaactctggTCACAACTTTTTCACCAAAAGATTAAC |
| *pEAQ-DaUGT73GZ2-F* | cccaaattcgcgaccggtATGGCTTCCAAAACCCAAAAC |
| *pEAQ-DaUGT73GZ2-R* | tcgaggcctttaactctggTCAGTGTAAAATTTCAGCATTCTCAAC |
| *pEAQ-UGT85J2-F* | cccaaattcgcgaccggtATGGAAGAAATCAGTTCAAAGCC |
| *pEAQ-UGT85J2-R* | tcgaggcctttaactctggGAAGTTTTGCTAGCAAAATCCTAA |
| *pEAQ-DaUGT94BA4-F* | cccaaattcgcgaccggtATGGCCATCAACGAACACCACCGT |
| *pEAQ-DaUGT94BA4-R* | tcgaggcctttaactctggTAACACATTGGCATCATTTTTTTC |
| *pET28a-anc156* | Synthesized |
| *pET28a-anc160* | Synthesized |

**Supplementary Table 19. Primers used in the site-mutations of DaUGT73GZ2 for heterologous expression.**

| **Name** | **Primer sequences** |
| --- | --- |
| L15A-F | gccATGAGCCAGAGCCATGTTATTCCGATGAC |
| L15A-R | TGGCTCTGGCTCATggcCGGCAGCAGCAGAAAATGC |
| H20A-F | AGCgccGTTATTCCGATGACCGATTTTGGTAA |
| H20A-R | ATCGGAATAACggcGCTCTGGCTCATCAGCGGC |
| M87A-F | CGAAAATgccGATGAACTGACCAGTCAGGATCTG |
| M87A-R | GTTCATCggcATTTTCGCAACCCTGCGGCAGA |
| Q93A-F | AACTGACCAGTgccGATCTGAGCCATAAATTTGTGCAG |
| Q93A-R | ATCggcACTGGTCAGTTCATCCATATTTTCGC |
| S96A-F | GGATCTGgccCATAAATTTGTGCAGGGTAGCAGC |
| S96A-R | ATTTATGggcCAGATCCTGACTGGTCAGTTCATC |
| S127A-F | TTATTAGTgccACCGCCCTGTATTGGGCAGGT |
| S127A-R | GGCGGTggcACTAATAATGCAACTAATTTTCGGCT |
| T128A-F | ATTAGTagtgccgccCTGTATTGGGCAGGTGAACTGG |
| T128A-R | AGggcggcactACTAATAATGCAACTAATTTTCGGCT |
| Y150A-F | TGTTTTCCAGACCgccAGCTGCTTTACCAGCCTGTGTG |
| Y150A-R | TggcGGTCTGGAAAACATAACGCGGAATTTTA |
| S162A-F | CCTGattgccggtGGCCCGCTGCAGGAAGTGA |
| S162A-R | GGCCaccggcaatCAGGCCGGCACACAGGCTG |
| Q195A-F | GAAgccATGCGCATGATGCTGAGCGATGTTCG |
| Q195A-R | ATCATGCGCATggcTTCCGGCATCTGGGCGCG |
| V207A-F | TGCCATGgccATGAAAATGGGTGATGCCATGC |
| V207A-R | TTTTCATggcCATGGCACGAACATCGCTCAGC |
| M210A-F | GATGAAAgccGGTGATGCCATGCGTAGCAGCC |
| M210A-R | CATCACCggcTTTCATCACCATGGCACGAACA |
| R239A-F | CTATCGCaaagccgaaAATAATCTGTGGTGCATTGGTCC |
| R239A-R | TttcggctttGCGATAGGCTTCAACATAGCTGC |
| F389A-F | ATGgccGCAGATCAGTTTCTGAATGAAAAATT |
| F389A-R | AACTGATCTGCggcCATCGGCCAGGTAATCATCG |

**Supplementary Table 20. Primers used in the site-mutations of DaUGT85CA7 for heterologous expression.**

| **Name** | **Primer sequences** |
| --- | --- |
| I31L-F | CGCAAAACTActaCATCAAAAGGGTTTCCACATAACA |
| I31L-R | GATGtagTAGTTTTGCGAGTTTTAGTGATGGA |
| L51D-F | TAgatTCTTCTCTCGGCCCCCACGCACTGGAA |
| L51D-R | GCCGAGAGAAGAatcTATGCTATTGTAGTTGTAATCTGGTATAACA |
| D73G-F | TATGGCACTCggcGTTCCTCCGGCCAACTCGC |
| D73G-R | GAACgccGAGTGCCATACTCTTGAACTGGAAA |
| D84Q-F | TCCACCAcagTACTTTGGCGTGCTAATGAAGAGC |
| D84Q-R | CAAAGTActgTGGTGGAGGCAGCGAGTTGGCC |
| F86I-F | AGACTACattGGCGTGCTAATGAAGAGCATTT |
| F86I-R | GCACGCCaatGTAGTCTGGTGGAGGCAGCGAG |
| Q128F-F | GATGACAttcAGCACTGTGGTGGCTGAAGAGC |
| Q128F-R | CAGTGCTgaaTGTCATCACCCCATCTGATATAATG |
| G147S-F | GATCACTagcGCAAATGGCTTTACAAACATTGC |
| G147S-R | CATTTGCgctAGTGATCCAAGATAGTGCAATAGGAA |
| N149C-F | TGGGGCAtgtGGCTTTACAAACATTGCCCGTC |
| N149C-R | TAAAGCCacaTGCCCCAGTGATCCAAGATAGT |
| C164G-F | TCAGGAAAggcGTAGCTGGACTTAGAGATGGCAGTAT |
| C164G-R | AGCTACgccTTTCCTGACAAGATCCCGATGAC |
| G171M-F | GGACTTAGAGATatgAGTATCGATATCAAAAGCGAAGAATT |
| G171M-R | CTcatATCTCTAAGTCCAGCTACACATTTCCT |
| E244A-F | GCGTTGGATgcaATTAGAAAGAAAGTTCCGCATGTATATAG |
| E244A-R | CTAATtgcATCCAACGCCTCCTTTTCTAGGCC |
| S254T-F | TactGTAGCACCTCTTCCACTCCTACTAAAGA |
| S254T-R | GAAGAGGTGCTACagtATATACATGCGGAACTTTCTTTCTAATT |
| D277S-F | GTCGGAagcGATATATGGAAAGAAGACGTTGAGTGTT |
| D277S-R | CATATATCgctTCCGACTAATTTTAAAACTCTTTCGT |
| S302N-F | aatTTCGGTACTTTTGCAATTACAAGCCAGCA |
| S302N-R | GCAAAAGTACCGAAattAACATAAAGAACCGAGTTTGGTGC |
| S310T-F | GCAATTACAactCAGCAACATCTTATAGAAATTGCATG |
| S310T-R | TGCTGagtTGTAATTGCAAAAGTACCGAAGCT |
| H313Q-F | CCAGCAAcagCTTATAGAAATTGCATGGGGACTTG |
| H313Q-R | CTATAAGctgTTGCTGGCTTGTAATTGCAAAA |
| E366I-F | CTGGTGCAAACAGattCAAGTTCTAAATCACCCGGCA |
| E366I-R | GaatCTGTTTGCACCAGCTCTTTATTAGTCCA |
| Q367I-F | GTGCAAACAGGAGataGTTCTAAATCACCCGGCAATAGG |
| Q367I-R | CtatCTCCTGTTTGCACCAGCTCTTTATTAGT |
| V386T-F | GAACTCGactCTCGAAAGCATTACTGCAGGAA |
| V386T-R | TTTCGAGagtCGAGTTCCACCCACAATGGCTA |
| L387I-F | GTGGAACTCGGTAattGAAAGCATTACTGCAGGAATTCC |
| L387I-R | CaatTACCGAGTTCCACCCACAATGGCTAAAG |
| G403A-F | CCATACTTTgcaGAGCATATAATAAGTAGCAAAATGATTTG |
| G403A-R | TGCTCtgcAAAGTATGGCCGACAAATCATAGG |
| H405Q-F | CATACTTTGGAGAGcaaATAATAAGTAGCAAAATGATTTGTGAAGAA |
| H405Q-R | ttgCTCTCCAAAGTATGGCCGACAAATCATAG |

**Supplementary Table 21. Primers used for qRT-PCR analysis.**

| **Name** | **Primer sequences** |
| --- | --- |
| *ACT-F* | GGTAACATTGTGCTCAGTGGTGG |
| *ACT-R* | AACGACCTTAATCTTCATGCTGC |
| *DaβAS2-F* | ACAGCCATTCCTTCTTGGGG |
| *DaβAS2-R* | CCTCAACCCAACAAGCAAGC |
| *DaCYP714E29-F* | TACAGCAGTTTCAGCCACGT |
| *DaCYP714E29-R* | GCTTCTCTTGACACCACGGA |
| *DaUGT94BA4-F* | GGCTCGACAAGAAGAACCCA |
| *DaUGT94BA4-R* | CCGCTAAGCTCTAGTCCGTG |

**Supplementary Table 22. Gene identifiers and source genome information of sequences analyzed in this study.**

| Renamed_ID | Source_species | Gene_ID |
| --- | --- | --- |
| *DaβAS1* | *D. asperoides* | *Das02G004901.t1* |
| *DaβAS2* | *D. asperoides* | *Das04G003055.t1* |
| *DaβAS3* | *D. asperoides* | *Das03G003102.t1* |
| *DaMAS1* | *D. asperoides* | *Das04G003910.t1* |
| *DaMAS2* | *D. asperoides* | *Das07G002904.t1* |
| *DaMAS3* | *D. asperoides* | *Das09G002328.t1* |
| *DaOAS1* | *D. asperoides* | *Das03G001840.t1* |
| *DaOAS2* | *D. asperoides* | *Das07G001837.t1* |
| *DaOAS3* | *D. asperoides* | *Das07G001840.t1* |
| *DaCYP714E29* | *D. asperoides* | *Das09G003295.t1* |
| *DaCYP714E30* | *D. asperoides* | *Das03G003375.t1* |
| *DaUGT85CA7/J2* | *D. asperoides* | *Das07G002907.t1* |
| *DaUGT73GZ2* | *D. asperoides* | *Das09G004326.t1* |
| *DaUGT73GY3* | *D. asperoides* | *Das03G005585.t1* |
| *DaUGT73GY4* | *D. asperoides* | *Das03G005581.t1* |
| *DaUGT94CP2* | *D. asperoides* | *Das09G004046.t1* |
| *DaUGT94CP3* | *D. asperoides* | *Das09G004048.t1* |
| *DaUGT94BA4* | *D. asperoides* | *Das03G004052.t1* |
| *DaUGT94CS1* | *D. asperoides* | *Das09G004049.t1* |
| *DaUGT94AY3* | *D. asperoides* | *Das05G003790.t1* |
| *DaUGT94CN8* | *D. asperoides* | *Das06G003895.t1* |
| *LjUGT85CA2* | *L. japonica* | *Lj1C1074T6.1* |
| *TgUGT85CA3-1* | *T. glandulifera* | *TgChr04G16778.1* |
| *TgUGT85CA3-2* | *T. glandulifera* | *TgChr04G16777.1* |

**Supplementary Table 23. Amino acid sequences of ancestral enzymes reconstructed from phylogenetic internal nodes.**

| **Ancestral enzyme nodes** | **Amino acid sequences** |
| --- | --- |
| **anc156** | MGSMSSSQKPHAVCIPFPAQGHINPMLKLAKLLHNKGFHITFVNTEYNHNRLLKSRGPNSLDGLPDFRFETIPDGLPPSNALDATQDIPSLCESTSKNCLAPFRELLRKLNDSSSSNVPPVSCIVSDGVMSFTVEAAEEFGIPEVLFWTTSACGFLCYAQYRQLVERGLIPLKDASYLTNGYLDNTTIDWIPGMKDIRLRDFPSFIRTTDSNDFMLNFVIGEIDRASKASAIIFNTFDALEQDVLNALSSIYPRIYSIGPLHLLLKNQLPDNNKLKSIGSNLWKEDTECLEWLNSKEPNSVVYVNFGSITVMTPQQLVEFAWGLANSKHSFLWIIRPDLVTGDSVAVLPPEFVEETKERGLLASWCPQEQVLNHPSVGGFLTHSGWNSTLESISSGVPMICWPFFAEQQTNCRYSCNEWGVGMEIDNDVKRDEVEKLVRELMEGEKGKEMKKKAMEWKKKAEEATNSGGSSSMNLDKLIDEVLLAKR* |
| **anc160** | MGSMASSSKPHAVFIPFPISTHINPMLKLAKLIHQKGFHITFITTEYNHNSILSSRGPHALDGLPDFRFETIPIDVPPSNARHPAEYFASLLKSIRKNFLPLFRELLRKLNDDSSSGMPPVSCIVSDGVMTQSVVAAEELGIPVVLFWVTGASGFMSIARHRDLVRRCLAAVKDGTYEVNEDLDNTTIDFIPGKKGIRMRDFLNFMRTTHSNDFMIDSCVGEVERTSKASAIIFNTFEALERDVLNELSAMYSRVYSIGPLPLLLKNQVKDENVLKLVGDNIWKEDTECVEWLNSKEPNSVIYVSFGSLTVTSPKHLVEFAWGLAKSKHSFLWIIRPDMVVGETVMALPQEFIEETKERGLIVSWCQQEQVLNHPAVGGFLTHCGWNSVLESISAGVPMICWPFFGEHMTNCKYVCNEWGVGIEIDEDVKRGQVEKVVRELMEGEKGKKIREKAKEWKKIAQEATYPSGISSQNLDKLVDEVLLSKP* |
